# High-resolution lineage tracking of within-host evolution and strain transmission in a human gut symbiont across ecological scales

**DOI:** 10.1101/2024.02.17.580834

**Authors:** Kimberly S. Vasquez, Daniel P.G.H. Wong, Miguel F. Pedro, Feiqiao Brian Yu, Sunit Jain, Xiandong Meng, Steven K. Higginbottom, Brian C. DeFelice, Norma Neff, Ami Bhatt, Carolina Tropini, Karina B. Xavier, Justin L. Sonnenburg, Benjamin H. Good, Kerwyn Casey Huang

## Abstract

Gut bacteria rapidly evolve *in vivo*, but their long-term success requires dispersal across hosts. Here, we quantify this interplay by tracking >50,000 genomically barcoded lineages of the prevalent commensal *Bacteroides thetaiotaomicron* (*Bt*) among co-housed mice. We find that adaptive mutations rapidly spread between hosts, overcoming the natural colonization resistance of resident *Bt* strains. Daily transmission rates varied >10-fold across hosts, but shared selection pressures drove predictable engraftment of specific lineages over time. The addition of a 49-species community shifted the adaptive landscape relative to mono-colonized *Bt* without slowing the rate of evolution, and reduced transmission while still allowing specific mutants to engraft. Whole-genome sequencing uncovered diverse modes of adaptation involving complex carbohydrate metabolism. Complementary *in vitro* evolution across 29 carbon sources revealed variable overlap with *in vivo* selection pressures, potentially reflecting synergistic and antagonistic pleiotropies. These results show how high-resolution lineage tracking enables quantification of commensal evolution across ecological scales.

## Introduction

The mammalian gut harbors a diverse microbial community that provides myriad benefits to its host. The residents of this community contend with frequent changes in their host environment, which cause rapid shifts in the abundances of different species and strains^1^. While many studies have focused on these ecological dynamics, there is a growing recognition that rapid evolution of gut commensals can also play an important role^2–4^. Longitudinal sequencing of human and murine microbiomes has revealed that strains can evolve within individual hosts via *de novo* mutations that sweep to large frequencies over weeks and even days^1,2,4^, hinting at a rich landscape of local adaptation. The contingencies of this landscape across host and community contexts are only starting to be explored^5–10^, and even less is known about how these adaptive mutations spread across multiple host communities. Previous work suggests that horizontal transmission is often constrained by the colonization resistance of a healthy microbiota^11,12^, which hinders the invasion of strains that compete for occupied ecological niches. The interplay between local adaptation, transmission, and colonization resistance is important for the design of microbiome-based therapies, and for understanding the long-term evolution of gut commensals across the broader host population^13,14^. However, the relative impacts of each of these factors, operating across ecological scales, remain challenging to tease apart in uncontrolled, natural settings^15^.

Experimental evolution of gut commensals in model hosts such as mice has been a powerful tool for quantifying *in vivo* evolution^5,7–10,16–18^. Most studies have focused on the model enteric bacterium *Escherichia coli*, and have elucidated *in vivo* selection pressures^19–21^ and the effects of other community members on the mutations that drive evolution^10,20,22,23^. Some mutations repeatedly arose in independent evolution experiments and distinct *E. coli* strains^9,10,17^, suggesting strong and consistent selective pressures in the mouse gut. Recent work has started to extend these findings to species from the more prevalent and abundant gut genus *Bacteroides*^6,7,24,25^, finding parallel evolution across hosts but strong dependence on host diet^5^ or inflammation state^26^.

The ability of locally adaptive mutants to spread across hosts is intimately connected with the tradeoffs that they encounter in other environmental conditions. Gut bacteria inhabit a complex environment, which is shaped by interactions with their host and other members of their local community. Our limited understanding of this landscape leaves many basic questions unresolved. Does the presence of a diverse community slow down evolution by closing off adaptive niches^20,27^, or speed up evolution due to cross-feeding and other opportunities for ecological diversification^28,29^? Are mutations that arise in a single host transmitted broadly, stochastically, or selectively across hosts? How does transmission and engraftment depend on the host, the focal species, and the presence of other community members?

Traditional approaches based on isolate sequencing or shotgun metagenomics are ill-suited to address these questions, because low throughput and low temporal resolution make evolutionary trajectories difficult to track. These limitations are especially apparent in diverse communities like the gut microbiota, in which the species of interest may comprise only a small fraction of cells. Previous studies were able to distinguish small numbers of lineages with high temporal frequency using fluorescent proteins as labels, and whole genome sequencing of clones to identify adaptive mutations^5,8,10,17,22^. Genomic barcoding increases the throughput of lineage tracking, enabling quantification of evolutionary forces such as mutation, drift, selection, transmission, and engraftment both *in vitro*^30,31^ and *in vivo*^9,26^. For instance, in a previous study, we cross-housed gnotobiotic mice colonized with ∼200 barcoded *E. coli* strains to study the interplay between within-host selection and inter-host transmission, and used mathematical modeling to estimate that the transmission rate was ∼10% per day^9^. However, it remains unclear how these observations in *E. coli*^8–10,16,17,20,22,32,33^ – a facultative anaerobe typically at low abundance in mammalian feces – generalize to the obligate anaerobes that dominate the gut microbiota and provide a larger contribution to its total metabolic potential.

In this study, we developed a genomic barcoding platform for high-resolution lineage tracking of the model human gut commensal *Bacteroides thetaiotaomicron* (hereafter *Bt*). We introduced a library of >50,000 barcoded *Bt* strains into germ-free mice under various diets, housing conditions, and microbiota diversities – including a synthetic community of 49 strains derived from a single human host – to track evolutionary dynamics within and across hosts over ∼2 months. Our high-resolution view allowed us to distinguish transient passage of transmitted cells from long-term engraftment, and to quantify the dependence of these processes, as well as the landscape of adaptive evolution, on microbiota diversity. We combined barcode lineage tracking with isolate and metagenomic sequencing to uncover targets of selection largely related to complex carbohydrate metabolism. Comparisons with *in vitro* evolution experiments across 29 diverse carbon sources probed the replicability of *in vivo* selection pressures, and suggested that *Bt* must navigate a complex pleiotropic landscape in adapting to a new host. Collectively, our results establish the utility of high-resolution lineage tracking to address gut commensal evolution at various scales of spatial and community complexity.

## Results

### Creation of a highly diverse genetic barcoding system in a model human gut symbiont

To investigate the evolution of a representative commensal bacterium prevalent in human guts, we constructed a library of barcoded *Bt* strains arrayed across 192 pools of separately transformed cells (Fig. 1A). Separation of strains into small pools allowed us to assemble distinct combinations of barcodes into larger pools as inocula for hosts, enabling detection of transmission between hosts. Each pool was transformed by aerobic conjugation with *E. coli* carrying plasmids with unique 26-nucleotide DNA barcodes (Methods). The 192 pools of barcoded *Bt* strains were maintained separately in two 96-well plates, and sequencing demonstrated that each well contained ∼300 barcodes on average, for a total of ∼50,000-60,000 unique barcodes. These highly diverse inocula allowed us to quantify the adaptation of lineages as they grew and competed from very low frequencies^30^.

**Figure 1:**
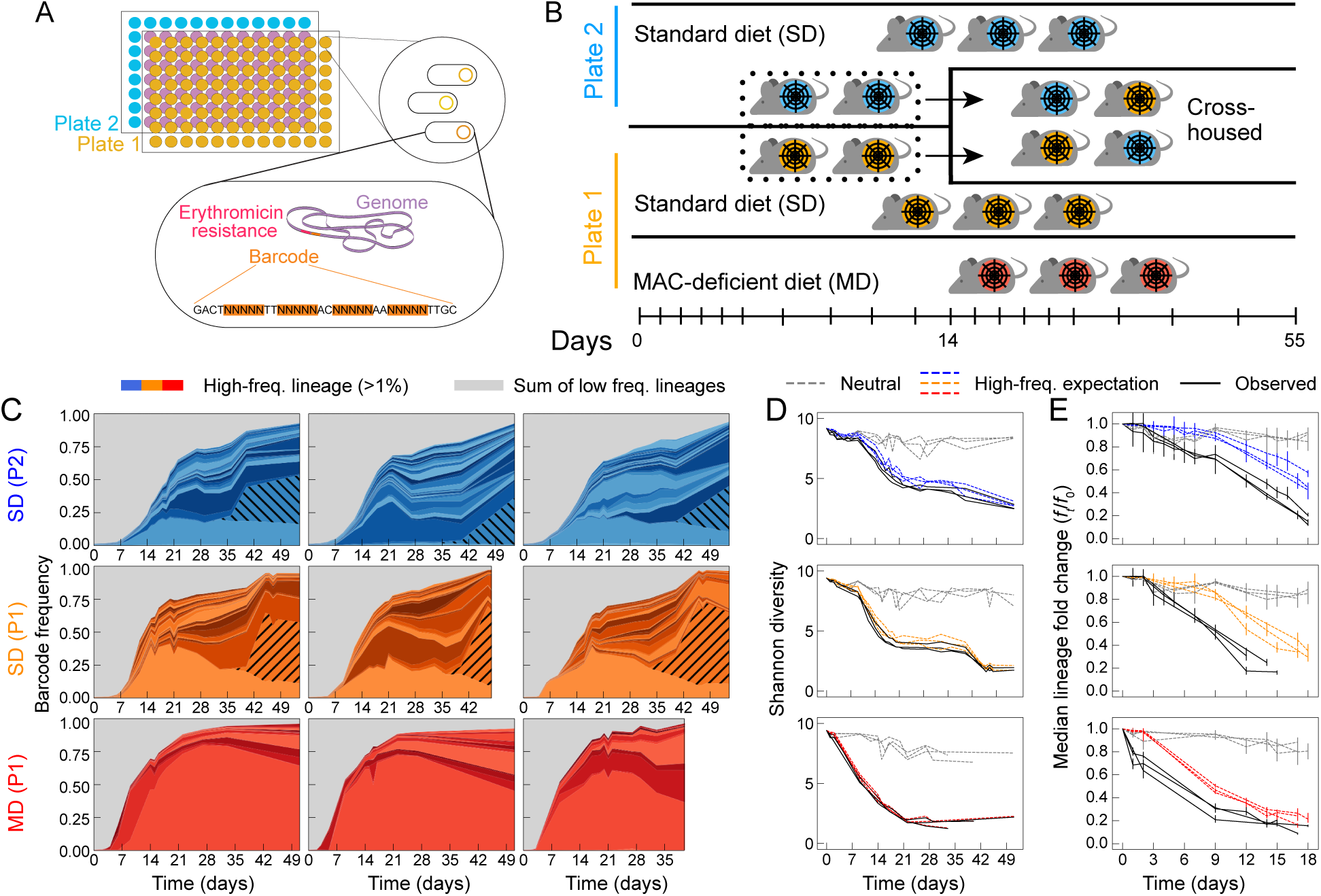
Genomic barcoding enables high-resolution lineage tracking of *Bacteroides thetaiotaomicron*. A) Construction of a library of strains with a genomically integrated DNA barcode and an antibiotic resistance cassette (Methods). Separate pools, each containing ∼200-400 barcoded strains, were maintained across two 96-well plates to permit downstream assembly of distinct inocula. B) Plate 1 (P1)- and plate 2 (P2)-colonized mice fed a standard diet (SD) were initially kept in separate isolators (5 co-housed mice per isolator). On day 14, two P1 mice were transferred to a cross-housing cage with two P2 mice in the P2 isolator. Three P1 mice fed a MAC-deficient diet were cohoused in a third isolator. C) Barcode lineage abundances in continually co-housed mice (rows; cross-housed mice are shown in Fig. 2). Strongly adaptive barcodes reaching >1% at some time point in a cage are color-coded, and all others are coarse-grained in gray. Hatched lineages are late-expanding barcodes LE1 and LE2. D,E) Dynamics of lineage diversity (one curve per mouse in (C)) compared to simulated expectations under neutral drift or under expansion of the high-frequency color-coded lineages in (C) alone (Supplementary Text). Bars in (E) represent the median and interquartile range of median lineage fold-changes (MFC) computed across a range of initial frequencies (Supplementary Text). High-frequency lineages drive the decline of observed Shannon diversity (D), but do not entirely account for the observed MFC decline (E), suggesting the presence of many more adaptive lineages at intermediate frequencies.

We grew each 96-well plate of barcoded *Bt* strains overnight and mixed the cultures within each plate at equal volume, creating distinct pools that we will refer to as plate 1 (P1) and plate 2 (P2). Barcode sequencing analysis identified ∼28,000 unique barcodes in P1 and ∼25,000 unique barcodes in P2 (Fig. S1A). Independent amplifications and sequencing of the same DNA preparation from the P1 pool (Fig. S1B) or from feces sampled on consecutive days in the same mouse (Fig. S1C) revealed strong correlation of barcode abundance down to a relative abundance of ∼10^-5^. Read dereplication with unique molecular identifiers (UMIs)^30^ revealed that some fecal samples had barcode relative abundances that were less well resolved (Methods). Nonetheless, the frequency resolution of most samples was ≤ 10^−4^ (Supplementary Text, Fig. S2), at least two orders of magnitude better than whole-genome sequencing-based approaches. Thus, our barcoded strain library enables high-resolution measurements of lineage dynamics.

### Barcoding provides a high-resolution view of adaptation to different diets

To demonstrate the utility of our barcoded strain library, we first investigated evolution of barcoded *Bt* populations upon mono-colonization of germ-free mice (Figure 1B). We gavaged the P1 and P2 strain pools into 13 mice housed in three cages across three isolators. In each of two isolators, five co-housed, standard diet-fed (SD) mice were inoculated with either the P1 or P2 pool. In the same isolator as the P1-inoculated mice, we transitioned three co-housed, germ-free mice to a diet lacking microbiota accessible carbohydrates (MACs) two weeks prior to gavage with the P1 strains. These MAC-deficient diet-fed (MD) mice were used to quantify the dependence of *Bt* evolutionary dynamics on diet, which has previously been shown to influence the targets of selection in *Bt*^5^. For all mice, we sampled feces and tracked *Bt* barcode populations (Methods) over ∼50 days, daily for the first week and twice per week thereafter. In all mice, *Bt* colonized and remained at high abundance (∼10^11^ CFUs/mL) over the course of the experiment (Fig. S1D). On day 14, four mice (two from each SD isolator) were cross-housed in a separate cage for the remainder of the experiment to study transmission (influx) and engraftment of external strains, as discussed in a subsequent section. This cross-housing design left three cages of three mice each to study the effects of diet.

In the two sets of SD mice, we observed the rapid expansion of a handful of barcodes to frequencies >1% over the first three weeks (Fig. 1C), consistent with previous observations of positive selection on *Bt* within weeks of colonization^5^. Among these high-frequency barcodes, expansion rates – a proxy for their relative fitness – ranged from 17-242% per day (median 46%, IQR 32-82%; Supplementary Text) and early expansion was accompanied by a decrease in the Shannon diversity of the larger library (Fig. 1D). Later expansion of other barcodes suggested multiple waves of selection, due to subsequent mutations in the barcoded lineages and/or shifting selection pressures within the mice. Such dynamics were most clearly illustrated by the delayed expansion of a pair of barcodes from each of the two isolators, which grew from nearly undetectable frequencies (<0.01%) on day 30 to ∼30-40% of the entire co-housed *Bt* population by day 55 (Fig. 1C, hatched lineages). The genetic bases of these distinctive barcodes (LE1 and LE2) are examined in a subsequent section. In contrast, other lineages rose to high frequencies before LE1 or LE2 but failed to expand in other co-housed mice (Fig. S3). Such localized expansions could be driven by host-specific adaptation, pleiotropic tradeoffs to transmission and engraftment, or neutral processes like colonization of a privileged spatial niche.

While the overall dynamics were highly reproducible across the two sets of standard diet (SD) mice, we observed high-frequency lineages emerging even faster in the MAC-deficient (MD) mice (Fig. 1C, S4. These lineages had significantly larger expansion rates (median 98% and IQR 62-140% per day) than the high-frequency lineages in SD mice (*p*=7×10^-3^, Mann-Whitney U-test), and a single barcode reached >40% frequency in all MD mice by day 14. This rapid expansion in MD mice was mirrored by the more rapid loss of Shannon diversity across the larger library compared to SD mice (Fig. 1D). Shannon diversity is strongly biased toward the frequencies of the largest barcodes, and indeed the expansion of the ∼10-20 highest frequency barcode lineages in each mouse (Fig. 1C) almost entirely accounted for the decrease in Shannon diversity over time (Fig. 1D).

An alternative measure, the median fold change (MFC) across many barcodes, captures the dynamics of more typical, low-frequency lineages. In a simple population genetic model in which most lineages have equal fitness, the MFC approximates the negative integral of mean fitness of the population over time (Fig. 1E, Supplementary Text), providing a more direct measure of the overall rate of adaptation of the population. We found that the MFC steadily declined after day 2, reflecting the competitive disadvantage of the bulk population as it competes against a smaller subset of adaptive lineages. However, this decline in the MFC was faster than expected based on growth of the high-frequency lineages (Fig. 1C). This discrepancy enabled us to infer that the high-frequency lineages (Fig. 1C) accounted for ≲1/2 of the total adaptation of the population and <1/10 of the total number of adaptive lineages at day 14 (Supplementary Text). Consistent with this inference, we found that >100 barcodes persisted at substantial frequencies (>10^-4^) in most SD mice at day 51-54 (Fig. S4). While these intermediate-frequency barcode lineages could in principle have survived replacement by residing in spatially protected regions of the gut, our quantitative analysis suggests that adaptative mutations also played a role in their persistence. These findings demonstrate that our diverse library of barcoded *Bt* strains can reveal features of within-host dynamics and evolution that are not accessible with existing sequencing methodologies.

### Colonization resistance is overcome by ongoing evolution and transmission

In addition to tracking lineages within hosts, our library design also enables us to measure the transmission of *Bt* strains between hosts by cross-housing mice inoculated with different barcode pools (Fig. 1B). Two P1 mice and two P2 mice from the SD isolators were cross-housed 14 days after initial colonization (in a separate cage within the isolator containing the other P2 mice), and *Bt* barcode dynamics were tracked for a further 40 days. We reasoned that this two-week delay between initial colonization and cross-housing would allow us to quantify *Bt* transmission well after any transient colonization dynamics within the mice, and after the resident *Bt* populations had already begun to adapt to their particular host^5^.

While high rates of inter-host transmission have previously been observed for the facultative anaerobe *E. coli* (∼10% per day)^9^, it remains unclear whether such high rates can be maintained in obligate anaerobes like *Bt*. Previous studies have shown that *Bacteroides* species exhibit colonization resistance upon single challenges with ∼10^8^ CFUs of invading strains^34,35^, suggesting that even a large influx of cells does not guarantee invasion. Nonetheless, we found that adaptive barcodes at high frequency in their originating mice were able to reciprocally invade the other populations and collectively expanded to >10% frequency by day 55 (Fig. 2A). These rapid invasions demonstrate that the rates of *Bt* transmission were high enough to overcome the inherent colonization resistance imposed by the locally adapted resident strains. However, these invasion dynamics were also highly asymmetric between mice. The P2 barcodes expanded in both P1-inoculated mice within a week of cross-housing, with 8-9 barcodes reaching >1% frequency in their new hosts by day 55 (Fig. 2A). On the other hand, the P1 barcodes did not substantially colonize the P2-inoculated mice until ∼3 weeks after cross-housing, with only 3-4 barcodes reaching >1% frequency in their new hosts (Fig. 2A).

**Figure 2:**
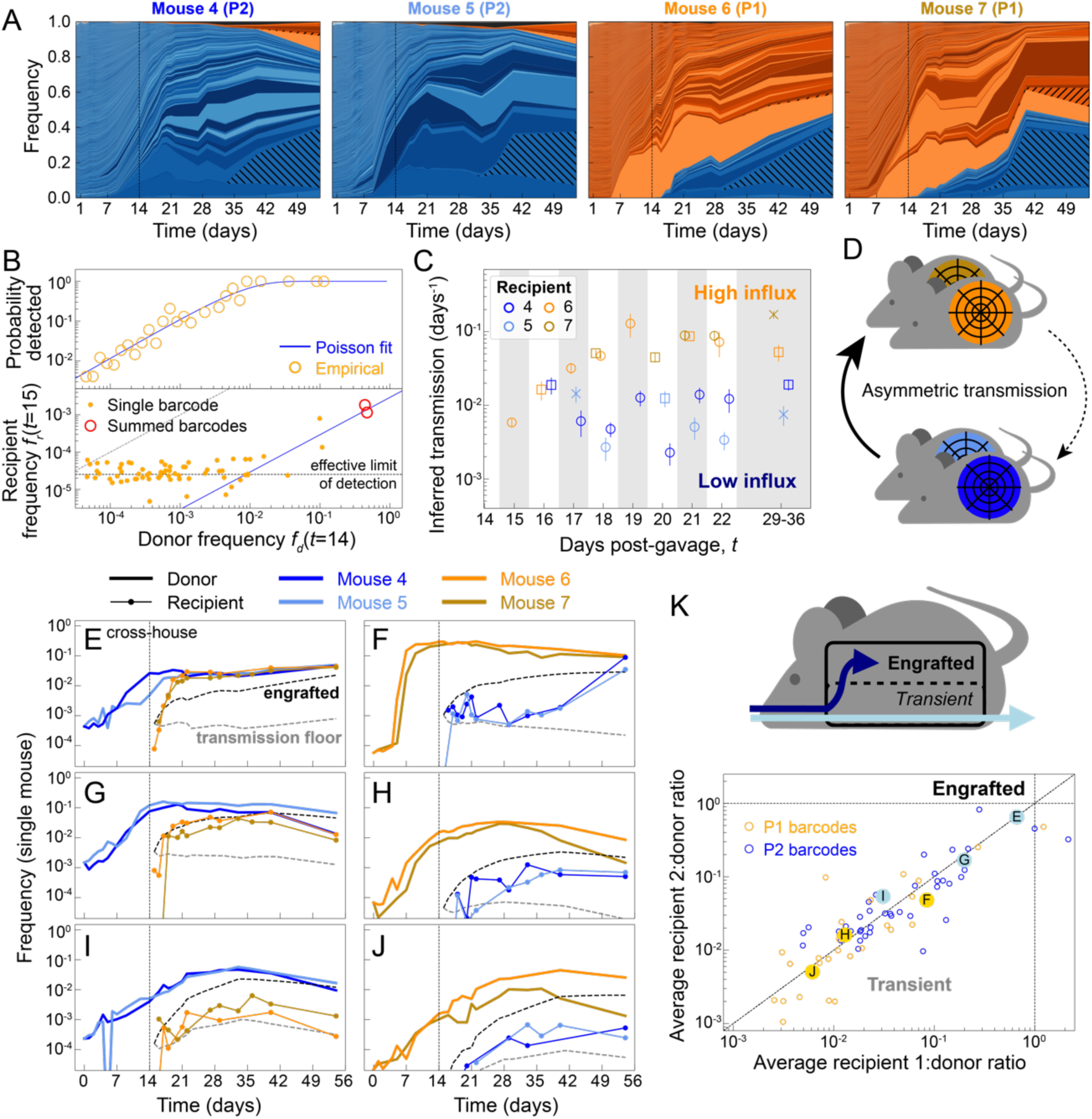
Variability in transmission and engraftment of lineages among cross-housed mice. A) Barcode lineage dynamics of cross-housed mice. Lineages reaching above 10^-5^ in at least one mouse are highlighted in shades of orange (P1) and blue (P2), reflecting their originating inoculum (Supplementary Text). A small percentage of lineages (gray) have unspecified inocula. LE1 and LE2 lineages are hatched. B) Transmission of individual P2 barcodes from donor mice (#4, 5) to recipient mouse #6 the day after cross-housing (Supplementary Text). Top: probability of detection in recipient on day 15 as a function of donor frequency at day 14 is consistent with a uniform transmission rate of ∼0.6%. Bottom: measured frequencies of barcodes in the recipient. While most barcodes were undetected or measured at the effective limit of detection, their summed frequencies (red circles, non-overlapping sets of barcodes) were consistent with estimates inferred from the frequency-resolved probability of detection. C) Extending (B) to measure single-day transmission rates over time (Supplementary Text), with maximum likelihood estimates ±1 standard error of the mean (SEM) fitted from the probability of detection of *n* donor lineages in the recipient on day *t* (*n* > 30 per estimate). For most points (circles), donor lineages used for fitting were identified on day *t* − 1. When single-day intervals (circles) were not sampled, donor lineages were identified on day *t* − 2 (squares) or *t* − 3 (crosses), since these intervals provided reasonable approximations of single day transmission (Supplementary Text). D) Measurements in (C) suggest strongly asymmetric transmission rates driven by host behavior. E-J) Trajectories of barcode lineages in donor and recipient mice. Two theoretical expectations of recipient trajectories, from a minimal model of inter- and intra-host dynamics, were parametrized using a donor trajectory and transmission estimates from (C). Gray represents daily transmission without engraftment in recipients, while black represents complete engraftment of transmitted cells (Supplementary Text). K) Top: The diversity of recipient trajectories in (E-J) support a strong, lineage-dependent distinction between transient passage and engraftment. Bottom: Average recipient:donor ratio (Supplementary Text) was highly correlated between paired recipients (mice #4,5 in orange, or #6,7 in blue).

To disentangle whether these asymmetric cross-invasion rates were driven by differences in barcode transmissibility, barcode fitness, or mouse behavior (e.g., differences in coprophagy), we focused on the transmission dynamics in the initial days after cross-housing. For example, in P1 mouse #6, many P2 barcodes were immediately detected after the first day of cross-housing (and undetected in the last fecal sample prior to cross-housing, as expected), indicating transmission of barcodes between mice (Fig. 2B). However, since most of these barcodes were measured close to the limit of detection (∼10^-5^), it is difficult to quantify their individual transmission rates on a barcode-by-barcode basis. Instead, we assumed that barcodes were uniformly transmitted and fit a simple population-genetic model to the likelihood of detection as a function of donor frequency (Supplementary Text). The estimated transmission rate of 0.6 ± 0.1 % over this single day interval provides a good fit to the data, and agrees with the average transmission rate found by aggregating frequencies across multiple barcodes (Fig. 2B). These findings suggest that any variability in transmission over this interval is small.

Building on this analysis, we used the same approach to measure transmission rates over subsequent time intervals during the first week of cross-housing (Fig. 2C,D, S5). In each interval, we quantified the fraction of lineages that were not present at the first time point and were present in the second, as a proxy for transmission in that interval. Note that beyond the first day of cross-housing, non-detection of a donor barcode in the recipient mouse could be due to finite sampling rather than true absence. If engraftment and persistence is sufficiently high, observation of a barcode at a subsequent time point could be dominated by past transmission events, rather than new transmission in the current time interval. This scenario would predict a steadily increasing apparent transmission rate as cells that arrived in earlier intervals accumulate in the recipient, as we observed in mouse #6 over the first few days of cross-housing (Fig. 2C). However, this >10-fold increase is quantitatively inconsistent with the measured transmission rate from day 14 to 15 or the strength of selection in these populations (<100% per day for the majority of lineages). Similarly, in mouse #4, the decrease in transmission rate over the first week was qualitatively inconsistent with persistence of previously transmitted cells. These results suggest that our estimates predominantly reflect transmission within a single measurement interval, and that this instantaneous rate strongly fluctuates over time.

Even more striking than the variation in transmission rate within a mouse was the systematic variation across mice, with lower rates into both P2 mice (#4 and 5, 0.2%-2%/day) compared with both P1 mice (#6 and 7, 0.6%-10%/day). While these differences in transmissibility could be driven by the evolution of the *Bt* populations in the two weeks prior to cross-housing, this scenario is unlikely to explain our data, since it would require the evolved transmissibility to systematically differ between P1 and P2 barcodes. Instead, our data suggest that the variation in transmission rates over time and across hosts was due to behavioral variation in the mice (e.g., differences in the frequency, timing, and/or preferences of coprophagy; Fig. 2C,D).

A crucial question is whether the transmitted lineages are able to successfully engraft in the recipient community, or whether they are only transiently passing through (e.g., due to spatial or metabolic niche priority effects^36^). In classical experiments involving simulated transmission events (“challenges”), engraftment can be distinguished from transient passage by long-term persistence of the transmitted strain^12,34,35^. To distinguish these scenarios in our continuous cross-housing conditions, we considered a simple dynamical model that incorporates the joint effects of transmission, engraftment, and selection, in which the fitness of each lineage can vary in a mouse-independent fashion (Supplementary Text). This model predicts that for a barcode under continual transmission without engraftment, its frequency in a recipient mouse should be set by the product of its frequency in the donor mouse and the daily transmission rate. This product defines a “transmission floor” that is consistent with transient passage alone. In contrast, an engrafted lineage will expand in the recipient mouse and will eventually approach the frequency in the donor population at long times. By comparing the observed barcode trajectories to these two extremes, we can classify the transmitted lineages along a continuum ranging from purely transient to fully engrafted.

We found that unlike daily transmission, which was relatively uniform across barcode strains, the engraftment rate varied widely across strains. We identified individual examples of adaptive barcodes (Fig. 2E-J) that spanned the full range of behaviors predicted by our dynamical model (Fig. 2K; Supplementary Text). Individual strains exhibited strikingly consistent dynamics across recipient mice (Fig. 2K), implying that evolved differences in engraftment ability, rather than stochasticity in the engraftment process, drove variation across barcoded strains. For instance, one P2 barcode rapidly approached the donor frequency in both recipient mice within one week of cross-housing (Fig. 2E), while a P1 barcode remained at very low frequencies in recipient mice (Fig. 2F) until day 40, after which it rapidly expanded to match the donor frequency. Other barcodes spanned two representative behaviors: partial convergence to the donor frequency (Fig. 2G,H) or never rising above the transmission floor (Fig. 2I,J), signifying transient passage without engraftment. Importantly, the degree of engraftment in a recipient mouse was not predicted by the rate of contemporaneous expansion in the donor mouse (compare Fig. 2G versus 2I, and Fig. S6), suggesting that relative growth rates alone cannot account for differences in engraftment. Instead, these data imply that barcoded strains strongly varied in their ability to engraft due to evolved physiological differences, or their ability to fill an underutilized niche (physical or metabolic) in recipient mice.

A prototypical example of this niche-filling behavior is shown in Fig. 2E. This lineage rapidly expanded in both recipient mice—more quickly than expected if it had equal fitness in the donor and recipient—and saturated once it reached an approximately equal frequency ∼2% with the donor. This frequency was then maintained over ∼40 days despite ongoing sweeps among other barcoded strains. These biphasic invasion dynamics are consistent with frequency-dependent selection to fill a niche supporting a fixed fraction of the population. In contrast, the more variable engraftment trajectories of other strains (Fig. 2G,H) are consistent with directional selection within the original niche of the *Bt* population. These examples illustrate how high-resolution lineage tracking enables the quantification of a diverse range of selection and transmission dynamics co-occurring within a single population.

In our experiments, *Bt* transmission strongly varied across murine hosts, potentially due to host behavior and sensitivity of *Bt* to the external environment. On the other hand, distinct hosts reliably selected the same subset of adaptive barcodes for engraftment and long-term persistence. Thus, although the rate of engraftment is fundamentally bounded by host-dependent rates of transmission, consistent *in vivo* selection pressures were sufficient to drive similar lineage dynamics across hosts. These findings provide further evidence that ongoing within-host evolution, whether directional or diversifying, can overcome colonization resistance imposed by already colonized and simultaneously evolving populations of the same strain.

The distinct engraftment dynamics in Fig. 2 suggest that the complex spatial structure of the gut^37,38^ may support the emergence and stable persistence of minimally diverged but spatially adapted strains. For instance, adaptive lineages could localize differentially along the intestinal tract following nutrient or pH gradients^39^, or to the mucus (walls) versus the lumen (interior) of the intestine^40^. To probe this within-host spatial variation, we dissected the cecum (beginning of the large intestine) of each mouse at the time of sacrifice to compare the relative abundances in homogenized samples of this region with a fecal sample from the same day (Fig. S7A). We reasoned that the cecum should reflect distinct environmental conditions^41^ and spatial niches^37,38^, compared to feces. Nonetheless, we found that the relative abundance of barcoded strains was broadly correlated between the cecum and feces across all mice that we examined (Fig. S7B). This correlation was maintained even among transmitted barcodes in recipient mice (Fig. S7C), regardless of engraftment rate (Fig. S7D-F). Thus, variable engraftment did not appear to be related to pre-empted colonization of the cecum, at least at the spatial resolution we probed.

### Diet-dependent emergence of structurally diverse mutations during later stages of adaptation

To interrogate the genetic drivers of the evolutionary dynamics in our experiments, we complemented our barcode analyses with sequencing of the whole genome of isolates and fecal metagenomes. To link the targets of adaptation to specific barcodes, we isolated and sequenced 479 colonies (Methods) obtained from all 10 SD mice and two MD mice at various time points, at least 9 days after gavage. These 479 isolates were associated with 161 distinct barcodes. Although only a minority (*n=*62) of these barcodes reached >1% frequency in any of the mice, our MFC analysis (Fig. 1E) suggests that most sampled strains had adaptive mutations by the time of isolate collection (Supplementary Text).

We identified a mixture of simple point mutations, short indels, and complex structural variants in these adaptive lineages (Fig. 3A, Table S1, Supplementary Text). Eight barcode lineages across SD and MD mice harbored point mutations at a specific site in *BT0867*, a gene in polysaccharide utilization locus (PUL) 12 associated with digestion of host mucins^42^. Both variants (T756I and T756A) also arose in separate experiments involving co-colonization of *E. coli* and unbarcoded *Bt* (Methods, Fig. S8), as well as in a previous study of *Bt* colonization of specific pathogen-free (SPF) mice^5^. In that previous study^5^, the T756I mutation was more strongly selected for in SPF mice fed the Western (MAC-deficient-like) compared with a standard diet^5^, consistent with our observation that the dominant barcode in MD mice (Fig. 1C) carried the T756I mutation. However, we also found that multiple lineages in SD mice (Fig. 2F,G) carried the T756I mutation and reached >10% frequency as well. Notably, different barcode lineages in different cages emerged with *BT0867* mutations, implying that these variants arose *de novo* in individual mice rather than in the library prior to colonization. These findings show how strong selection pressures and large population sizes *in vivo* can reliably and repeatedly generate single nucleotide variants. All other point mutations were associated with single barcode lineages (Fig. 3A, Table S1), and like *BT0867* were broadly associated with metabolic pathways. GO-enrichment analysis revealed a significant excess (*q*<2×10^-3^) of nonsynonymous mutations in *susC-* and *susD*-like genes (representing eight distinct PULs), which encode outer membrane proteins involved in uptake of large metabolites like starch^43^.

**Figure 3:**
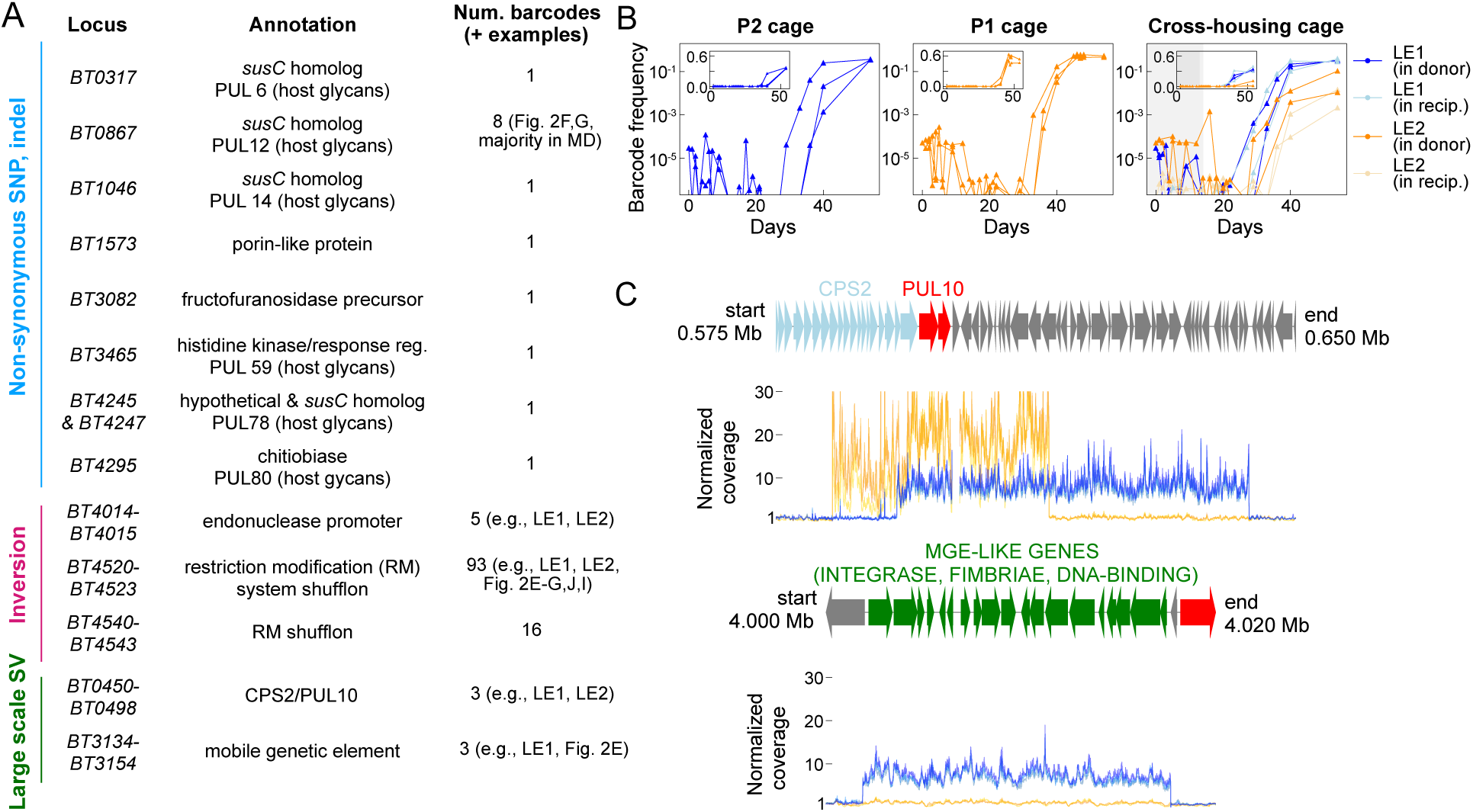
Isolate sequencing reveals diverse genetic bases of adaptation. A) Summary of the genetic mutations detected in >50% of isolates in at least two barcode lineages with 3 or more isolates. Many mutations are related to metabolism. B) The LE1 and LE2 barcodes shared similar dynamics, declining over the first 3-4 weeks before rapidly emerging after 30 days. C) Sequencing coverage of LE1 (blue) and LE2 isolates (orange), normalized by local coverage in a metagenome of the ancestor, shows extensive amplification of the same metabolic locus, CPS2/PUL10 (*n*=3 isolates per barcode). In LE1, this amplification is linked to the mobile genetic element (MGE)-like *BT3134*-*BT3154* locus.

Structural variants were more commonly shared among barcode lineages. One class of structural variants involved genomic spans of 180 to 2,000 bases flanked by inverted repeats (Table S2). These variants were likely instances of frequent and reversible RecA-mediated phase variation^44^. To rule out high-rate inversions arising during colony growth of the isolates, we applied additional statistical filters (Supplementary Text) and then validated their presence in fecal metagenomes (Fig. S9). Inversions passing our quality filters were primarily located in the binding domains and promoters of restriction modification systems involved in defense against bacteriophages and mobile genetic elements^45^. We observed 42 barcode lineages (8 supported by multiple isolates) with inverted regions and no other detected mutations, suggesting that this phase variation is sufficiently adaptive to drive the expansion of strains on its own.

Another structural variant was found in the two late-emerging barcoded strains, LE1 and LE2 (Fig. 1C, 2A), which were notable in both their frequency trajectories across mice and in the genetic bases of their adaptations. These barcode lineages exhibited striking and consistent behaviors across mice in the same cage (Fig. 3B), declining in frequency over the first ∼30 days before rapidly expanding at rates >90% per day over the next ∼10 days to reach intermediate frequencies of 30-60%. The expansion rate of these strains then dramatically declined over the final ∼10 days of the experiment (Fig. 3B). These dynamics were even synchronized across cages, and suggest that these adaptations arose prior to cross-housing and were transmitted across mice while persisting at very low frequencies prior to expansion. These observations suggest that these late-emerging barcoded strains experienced negative frequency-dependent selection and/or transient fitness benefits, in addition to clonal interference.

All 51 isolates associated with LE1 and LE2 barcodes shared an 8- to 20-fold amplification of a >20-kb region at the end of CPS locus 2, which contains PUL10 (of unknown function^42^, Fig. 3C, S10). Metagenomic sequencing confirmed the presence of these variants in feces from the mice with high abundance of the late-emerging strains, and in 4 of 5 mice in a second cohort of mono-colonized SD mice (Methods, Supplementary Text). In contrast, we did not observe this structural variant in isolate genomes from MD mice, nor in metagenomic sequencing of stool from mice co-colonized with a diverse synthetic community (Fig. S10). These findings suggest that the combination of mono-colonization and a standard diet pose a specific selection pressure for the amplification of this locus. A previous study of *Bt* in mono-colonized mice fed a standard diet observed repeated duplication and selection of a different PUL-associated locus, *BT1871*^25^. In our study, we detected complex structural variants in three other barcodes involving PUL79, PUL9 and 10, and PUL59, in addition to the LE1 and LE2 barcodes discussed above. In three of the five barcodes (including LE1 and the barcode in Fig. 2E), the complex structural variants involved transposition and/or amplification of a 21-gene mobile genetic element, *BT3134*-*BT3154* (Fig. 3C, S9). Taken together, these and previous observations^25^ suggest that structural variation, including and beyond canonical phase variation, can be an important source of adaptive genetic variation in a novel *in vivo* environment.

### Stable colonization of barcoded *Bt* with a diverse synthetic community of gut commensals

We next investigated whether *Bt* evolutionary dynamics were influenced by the presence of a phylogenetically diverse microbiota. To do so, we assembled a synthetic community composed of 49 strains isolated from a single human donor with representatives of all major phyla and most major families (Table S3). Notably, this community did not include a *Bt* strain, to avoid direct competition with the barcoded population, but it did include 7 other *Bacteroides* or *Phocaeicola* species (*B. fragilis*, *B. ovatus, B. intestinalis, B. uniformis, B. stercoris, B. caccae*, and *P. vulgatus*).

We introduced barcoded *Bt* populations into SD mice under three colonization conditions (Fig. 4A). In the control condition, mice were colonized with barcoded *Bt* alone. In the co-colonization condition, mice were gavaged with a 1:1 mixture of the synthetic community and a barcoded *Bt* pool. Finally, in the established community condition, the barcoded *Bt* pool was introduced two weeks after initial gavage with the community. This final condition enabled identification of ecological priority effects or transient effects between the host and community, and their consequences on *Bt* engraftment and evolution. Each colonization condition was introduced to three co-housed and two singly housed mice in a single isolator, so that we could observe the evolutionary dynamics with or without transmission. Within each condition, each of the mice were gavaged with a distinct set of barcodes, so that we could study the effects of transmission at the earliest stages of colonization. Four sets of barcodes included ∼2,700-3,100 barcodes each, while the fifth set was dominated by 109 barcodes at higher initial frequencies (to minimize the potential for pre-existing adaptive mutations). The same five sets of barcodes were used across all three colonization conditions, so that barcode behaviors across conditions could be compared.

**Figure 4:**
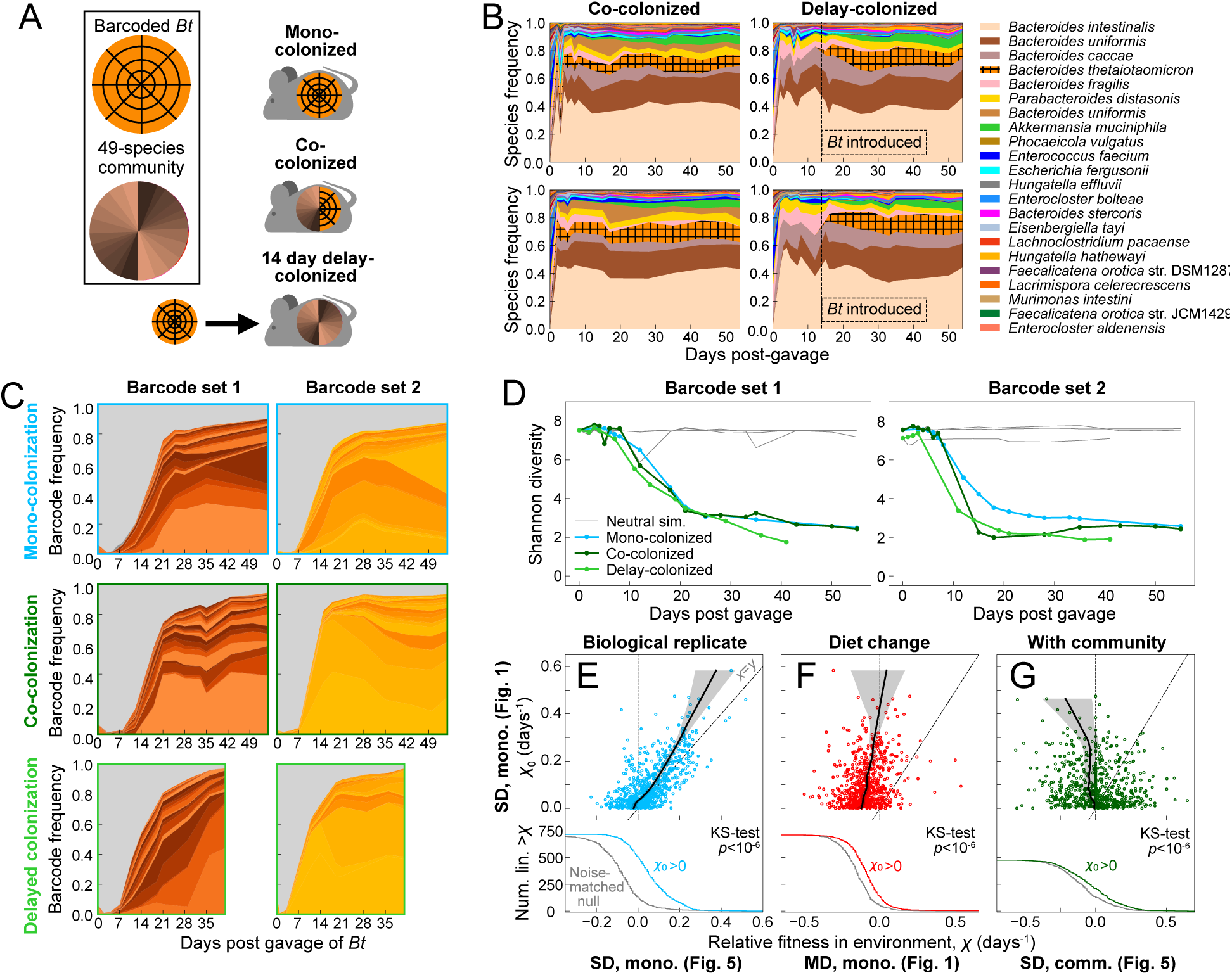
Rapid adaptation of barcoded *Bt* during stable co-colonization with a diverse 49-species community. A) Barcoded *Bt* populations were introduced to a cohort of SD mice under mono-colonization, simultaneous co-colonization with a 49-species community, or 14 days after colonization with the community. B) 16S rRNA gene sequencing in four singly-housed mice reveals stable, robust colonization, with similar species abundances under co- or delayed-*Bt* colonization, except for the extinction of *A. muciniphila* when *Bt* was delayed. C) Barcoded *Bt* lineage trajectories of singly housed mice in all three colonization conditions plotted as in Fig. 1C. Columns represent shared barcode inocula. D) Shannon diversity decreased similarly over time in each colonization condition. E-G) Individual lineage fitness varied across environments. For ∼718 low-noise lineages (Supplementary Text), relative fitness was measured over days 2-9 in mono-colonized SD mice from the first cohort (Fig. 1C) and compared to relative fitness over a similar interval (days 2-3 to 7-9) in (E) mono-colonized SD mice from the second cohort in (B), (F) mono-colonized MD mice (Fig. 1C), and (G) community co-colonized SD mice in (B). Top: points are individual lineages, and black curves with shaded regions are LOWESS fits (regressing *x*-axes on the shared *y*-axis, 30% of data for each point) with bootstrapped 95% confidence intervals (Supplementary Text). (E) Lineage fitnesses were highly correlated in replicate (SD-fed, mono-colonized) mice, owing to shared pre-existing adaptive variation. (F,G) Fitnesses were less correlated or weakly anti-correlated across diets (F) or community-colonization conditions (G), but were still typically fitter than a noise-matched null set of lineages (bottom, Supplementary Text).

In all mice gavaged with the synthetic community, 16S rRNA gene sequencing showed that the microbiota equilibrated within ∼2-3 days, reaching a stable composition similar to germ-free mice humanized with feces of the donor from which the community species were isolated^46,47^. The relative abundance of the *Bt* populalation was similar for the co-colonization or established community conditions, as was the rest of the community with the prominent exception of *Akkermansia muciniphila*, which was at ∼5% in the co-colonization condition but was undetectable in the delayed colonization condition, and did not recover once the barcoded *Bt* population was introduced (Fig. 4B, S11). Of the 49 strains in the community inoculum, 32 were detected at relative abundance >10^-4^ in a majority of the mice at ≥2 time point >14 days after colonization. These data indicate that a large fraction of the 49 strains successfully and stably colonized the mouse gut along with the barcoded *Bt* population, motivating their use as a model synthetic community.

### The presence of a diverse gut microbiota does not slow the pace of *Bt* evolution

Members of ecological communities are frequently conceptualized as occupying distinct niches provided by the environment. From this perspective, the evolution of a focal species into empty niches available during mono-colonization – as suggested by the dynamics of some lineages (Fig. 2E,F and LE1,2) – should be slowed or even prevented in a diverse community when the niches are already filled by other species^48,49^. Alternatively, a diverse community could provide a more complex metabolic environment and network of microbial interactions, generating new niches and maintaining or even accelerating evolution of a focal species, as has been previously observed in certain cases^28,29^. Comparing our singly housed mice under both mono-colonization and community colonization conditions enabled us to test these competing hypotheses. We found that the dynamics of high-frequency (>1%) barcodes were qualitatively similar across the three community colonization conditions: a few barcodes expanded to become a majority of the population within three weeks, regardless of the presence or absence of the community (Fig. 4C). Moreover, all *Bt* populations experienced similar rates of lineage diversity loss (Fig. 4D, S12) over the same time interval. These findings suggest that the presence of a community containing several closely related *Bacteroides* species did not substantially alter the pace of *Bt* evolution.

While this consistency of evolutionary dynamics across community conditions could be consistent with a balanced mixture of niche pre-emption and niche creation, it could also be consistent with community-independent evolution of *Bt* within its own isolated niche. The latter scenario predicts that individual mutations should confer the same fitness advantage independent of community context. To test this hypothesis, we focused on the subset of adaptive variants that were present in the initial library prior to colonization. Such pre-existing variation was previously observed to drive initial adaptation in other lineage-tracking experiments^7,17,36^, and provides a convenient way to quantify how the fitness benefits of a lineage vary across different environmental conditions. We identified pre-existing variants by observing barcode fold-changes (relative fitnesses) over days 2-9 across two cohorts of independently housed SD mono-colonized mice (represented in Fig. 1 and Fig. 4, respectively). A barcode’s behavior across independent mice should be uncorrelated, unless deterministic selective forces act on genetic variation shared between separately colonized mice^7^ (Supplementary Text). Indeed, we found a strong correlation in fitnesses within replicate mice sharing the same conditions (Fig. S13), in particular among the ∼718 fittest well-measured strains that expanded in the first cohort of mice (Fig. 4E, Supplementary Text).

In contrast, fitnesses among this cohort of SD-adaptive barcodes during mono-colonization were less well correlated with their fitnesses in MD mice (Fig. 4F), and virtually uncorrelated with those during co-colonization with the community (Fig. 4G). Indeed, most of these strains declined in frequency in these other host diet or community conditions. Nonetheless, as a group, the SD-adaptive barcodes were collectively enriched for higher fitness in both MD and co-colonized mice compared to noise-matched null collections of barcodes (Fig. 4F,G, bottom; Supplementary Text). This enrichment suggests that there is some overlap in the adaptive landscape across SD and MD mice (Fig. 4F, bottom), consistent with the presence of *BT0867* mutations in isolates derived from both SD and MD mice. On the other hand, there was more limited genomic corroboration of overlap between monocolonization and community-colonization conditions: metagenomic analyses of fecal samples from community-colonized mice provided no evidence of mutations in *BT0867* (Fig. S9) or amplifications of the CPS2/PUL10 locus that was present in the LE1/2 lineages (Fig. S10). Together, these observations suggest that community complexity influences the adaptive landscape of *Bt* through a mixture of niche preemption and niche creation, and that similar apparent rates of adaptation can mask broader shifts in the underlying targets of adaptation.

### A diverse microbiota slows migration between hosts

Our cross-housing experiments (Fig. 2) showed that adaptive lineages can rapidly overcome the colonization resistance of an established and locally adapted *Bt* population. The colonization resistance of the mammalian gut microbiota is known to positively correlate with community diversity^50,51^, as well as the timing of challenges with respect to transient environmental perturbations^52^. We sought to test both these aspects of colonization resistance using our co-housing experiments with the three community colonization conditions.

To test how colonization resistance depends on the timing of invasions relative to initial host colonization, we examined the co-housed mono-colonized mice that contained distinct *Bt* barcode pools (Fig. 5A). This immediate co-housing scenario differed from the experiments in Fig. 2, in which mice were cross-housed 14 days after mono-colonization. Among the three co-housed mice in the mono-colonization condition, we identified one pair of mice in which barcodes were significantly enriched on day 3 in their mouse of origin relative to the other mouse (Fig. 5B,C). Over the next 10 days, thousands of barcodes rapidly equilibrated in abundance across these two mice, consistent with an effective transmission with engraftment rate of ∼29% per day over the two weeks post-gavage (Fig. 5B,C, Supplementary Text). These dynamics are consistent with the hypothesis that there is little colonization resistance in the initial days after colonization of germ-free mice. However, this global convergence contrasted with more limited transmission we observed to and from the third co-housed mouse (Fig. 5B), as well as the lower engraftment we detected in the cross-housed mice in Fig. 2. The maintenance of high diversity argues against aberrant colonization dynamics among the first pair of mice that might permit such high rates of invasion, and measured transmission rates were high even after day 14. Instead, this variation across specific pairs of mice provides further support that transmission is strongly modulated by mouse-dependent social or coprophagic behaviors, which could be explained by the previously demonstrated propensity for mice to ingest feces directly from the anus of other mice rather than from the cage floor^53^. Nonetheless, as in the cross-housing experiment (Fig. 2A), we found that all three co-housed mono-colonized mice converged in their high-frequency barcode composition over the course of the experiment (Fig. 5B), suggesting that shared selection pressures can eventually overwhelm variation in the rate of migration.

**Figure 5:**
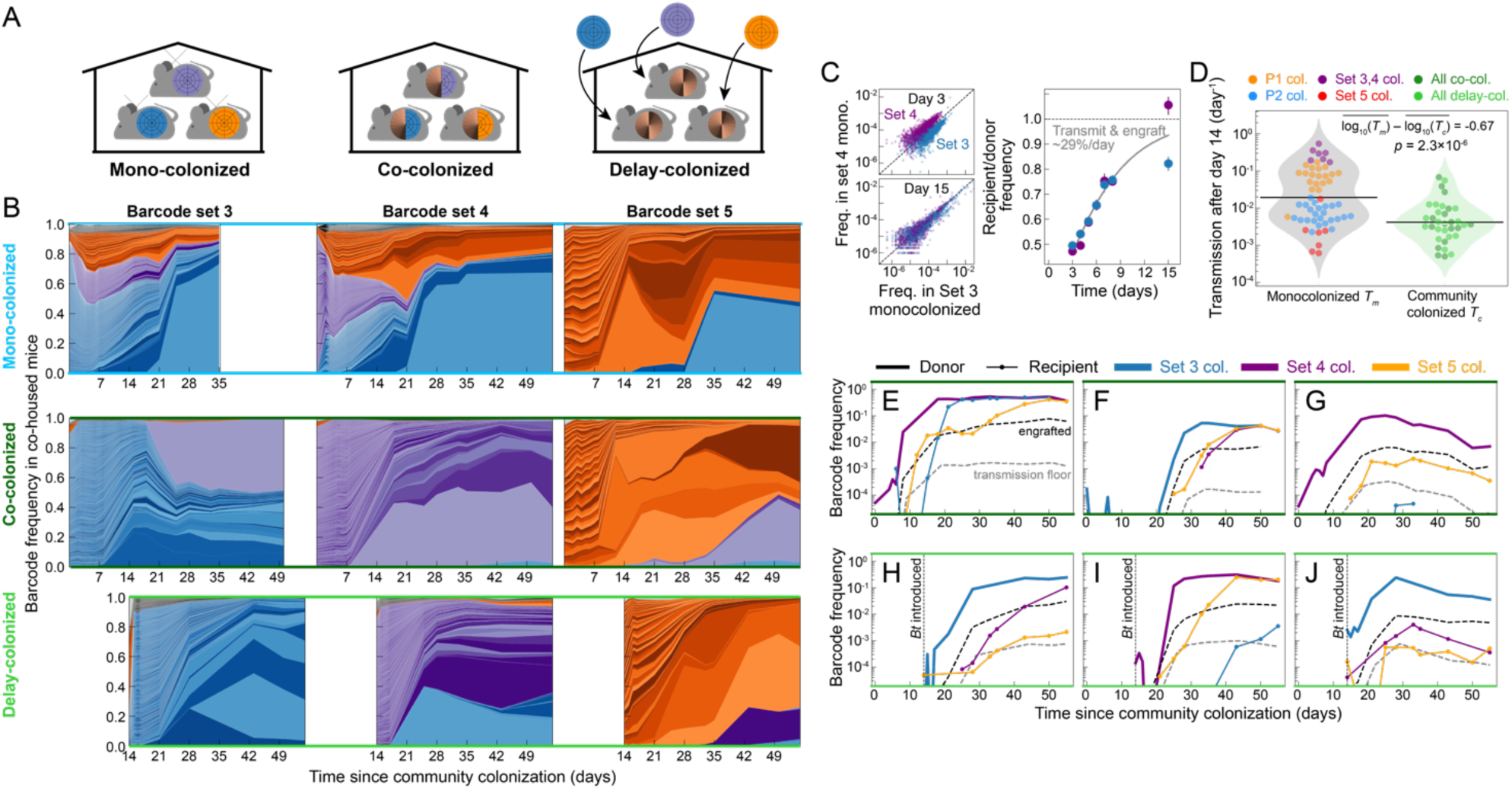
Transmission and engraftment are altered but not prevented by the presence of a diverse community. A) For each community colonization condition, three co-housed mice were gavaged with distinct barcode inocula. B) *Bt* lineage abundances shaded by inoculum, analogous to Fig. 2A. Rows are co-housed mice, and columns are mice inoculated with the same barcode set (blue: set 3, purple: set 4, orange: set 5). C) Frequencies of thousands of lineages rapidly converged across one pair of mono-colonized mice (sets 3 and 4), consistent with a combined rate of transmission and engraftment of ∼29%/day during the first two weeks of colonization (Supplementary Text). Points and error bars are median and IQR. D) Distributions of single day *Bt* transmission rates >14 days after colonization with *Bt* and/or the community, aggregated with measurements from Fig. 2C. Each point is colored by the recipient’s barcode inoculum or community-colonization condition. The mean (log) transmission rate was significantly lower in community-colonized mice than in mono-colonized mice, although the difference is much less than the variation among mono-colonized mice. E-J) Representative lineage trajectories in co-housed, community-colonized mice, analogous to Fig. 2E-J. Individual lineages were able to engraft in both co-colonized (E-G) and delay-colonized (H-J) recipient mice, but with less synchrony across recipients than under mono-colonization (Fig. 2E-J).

In contrast to mono-colonized mice, we found that mice colonized with the diverse synthetic community retained distinct *Bt* barcode compositions at 55 days post-colonization, which were biased toward their initial inocula (Fig. 5B). Previous work observed that invasion of a focal species was slowed in the presence of a diverse gut microbiota^54^; in our case, the *Bt* species still engrafts to high levels with the synthetic community (Fig. 4B). Nonetheless, subsequent transmission of *Bt* barcoded strains was slowed compared to in mono-colonized mice by some interaction between the community and the resident *Bt* population, although this difference is smaller than the variability across mice and over time (Fig. 5D, S14). Despite this overall slowdown, we nonetheless identified multiple examples of engrafting lineages. In total, 5 barcodes in the co-colonized mice and 15 barcodes in the delay-colonized mice reached >1% frequency in multiple co-housed mice, compared to 23 lineages in the mono-colonized mice. The migrating lineages showed diverse behaviors: some rapidly converged to the abundance of the donor barcode (Fig. 5E,F,H,I), while others converged more slowly, if at all (Fig. 5G,J). As in our cross-housing experiment (Fig. 2), the diversity of engraftment behaviors across barcodes contrasted with the relatively consistent, if asynchronous, behavior of each barcode across recipient mice, indicating that deterministic forces – rather than dispersal limitations or ecological drift – were the primary factors driving barcode dynamics in the host meta-population. Thus, even a community with relatively high diversity does not preclude engraftment of specific adaptive lineages in the face of ongoing transmission.

### Comparing within-host and *in vitro* evolution

Consistent with previous studies of *Bt* evolution *in vivo*^5,25^, we found that mutations in metabolism-related genes were rapidly selected for in the mouse gut, suggesting that the nutrient environment at least partially drives selection. The biological and chemical complexities of mice (even when mono-colonized) present challenges for identifying specific selection pressures that drove expansion of mutations. We sought to test whether *in vitro* evolution on single carbon sources would replicate the pace and targets of selection, thereby providing insights into the selection pressures faced *in vivo*. We selected ∼30 carbon sources (both simple and complex carbohydrates) that were either detected in NMR profiles of germ-free mouse ceca/feces (Fig. S15, Table S4) or are common components of human diets. We grew wild-type *Bt* in minimal media supplemented with each of the carbon sources individually at 0.25% (w/v) concentration to determine whether visible growth was supported over 48 h. From these tests, we focused on 29 conditions: four monosaccharides (glucose, glucose with supplemented iron and heme^55^, fructose, and galactose), five disaccharides (lactose, sucrose, maltose, trehalose, melibiose), and 20 oligo- and polysaccharides (including raffinose, stachyose, maltodextrin, and various commercial fibers).

We split the barcoded *Bt* library into six pools of ∼5,000 strains each. We then inoculated each well in a 96-well plate with one of the barcode pools in a given carbon source condition, and ensured that each well was surrounded by wells with different barcode pools to enable detection of contamination from neighboring wells. The *Bt* populations were passaged with a 1:200 dilution every 48 h for ∼2 months, equivalent to ∼230 generations of evolution assuming that they reached saturation in each passage (Fig. 6A). Each carbon source was associated with two disjoint barcode pools, each of which was passaged in two independent replicates. This replicated design enabled us to distinguish *de novo* mutations from pre-existing variants using the methods in Fig. 4E-G. We measured the growth dynamics of each well every 2 weeks (7 passages) to identify changes that would signify evolution (Table S5). Most populations exhibited enhanced growth through substantially increased yield, with some conditions also exhibiting increased growth rates (Fig. 6B, S16).

**Figure 6:**
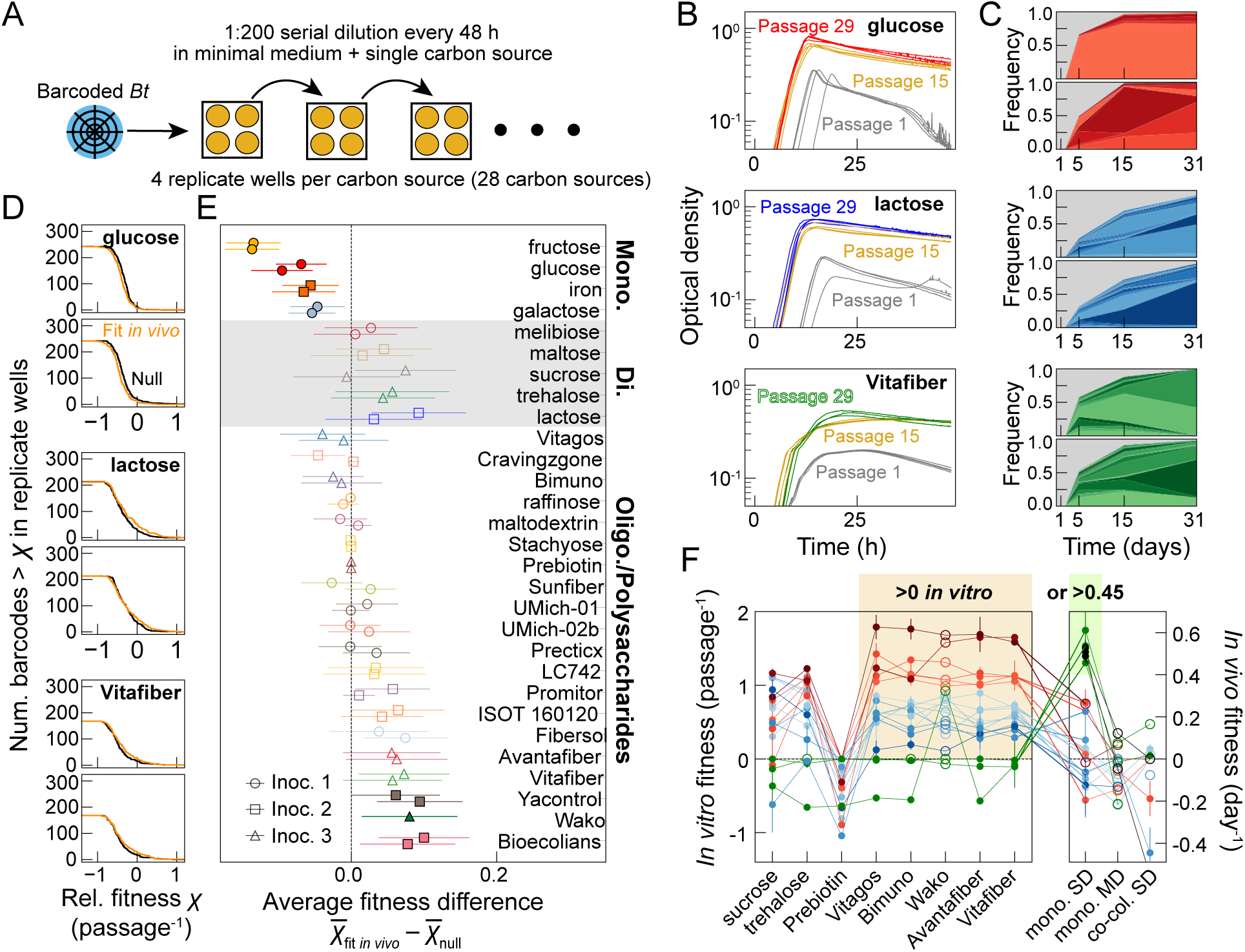
Correlation between *in vitro* and *in vivo* fitness effects increases with carbon source complexity. A) Pools of barcoded strains were passaged for ∼9 weeks in minimal medium supplemented with one of 29 carbon sources. For each carbon source, each of two pairs of wells were inoculated with the same barcode pool (four total wells). B) In three representative carbon sources, growth curves demonstrate decreased lag and increased growth rate and yield over 9 weeks of passaging. Each curve is a replicate well at a given timepoint. C) Barcode lineage dynamics in single wells representing the carbon sources in (B). D) Survival function of relative fitness over passages 1-5 of barcodes that were adaptive *in vivo (*Fig. 4E), compared to a noise-matched null set of lineages with the same distribution of frequencies after passage 1 (Supplementary Text). E) Mean fitness difference ±2 bootstrapped standard error between *in vivo*-fit barcodes and the null barcodes from (D) for every carbon source. Pairs of markers represent replicate wells, and bolded media are *q*<0.05 (two-sided *t*-test). *In vivo*-fit barcodes were consistently maladaptive in monosaccharides and frequently adaptive in polysaccharides. F) Fitness across environments of 21 barcode lineages with pre-existing variation that was consistently adaptive either in five polysaccharides over passages 1-5 or in SD mono-colonized mice (Fig. 5E). Each curve is a lineage, colored by median fitness in the five polysaccharides. Solid points and error bars are mean and range of two independent replicates and open circles represent single replicates. The ordering of lineage fitnesses is conserved across the five conditioned *in vitro* environments, but is scrambled *in vivo*.

Barcode sequencing at 6 timepoints (passages 0, 1, 2, 5, 15, and 31) revealed the rapid expansion of lineages across all the diverse carbon sources. These observations confirmed that the swift pace of evolution we observed *in vivo* was not specific to idiosyncratic features of the mouse gut (Fig. 6C, S17). We found that the apparent pace of evolution, as measured by decline in Shannon diversity, varied across carbon sources in each class of mono-, di-, oligo-, or polysaccharides (Fig. S18). There was also systematic variation between classes, with diversity loss over the first five passages faster in polysaccharides than in mono- or di-saccharides. The variability across carbon sources in otherwise identical environmental conditions suggests that *Bt* adaptation and its genetic bases were strongly influenced by the presence of specific carbon sources. We directly confirmed this genetic variation by measuring how adaptation to one carbon source affected growth on other carbon sources. We grew 12 populations evolved on diverse carbon sources in glucose, raffinose, and stachyose, all three of which are present at substantial concentration *in vivo* (Fig. S15, Table S4,S5). The evolved populations exhibited a broad range of growth behaviors in glucose (Fig. S19). Most grew faster than the ancestor, but populations evolved in glucose with iron, trehalose, and Vitafiber consistently saturated at lower maximum yields. The glucose-, fructose-, and galactose-evolved populations grew the fastest, signifying that the mutations enabled broad growth advantages across 6-carbon monosaccharides. In raffinose, the raffinose- and stachyose-evolved populations grew similarly to each other, with a much shorter lag time than any of the other populations (Fig. S19). Still, most evolved populations grew much faster and to higher yield than the ancestor, which grew only slightly. In stachyose, only the raffinose- and stachyose-evolved populations grew substantially within 36 h (Fig. S19). Thus, adaptive evolution of *Bt* in one carbon source can strongly impact growth (both positively and negatively) in other carbon sources. These findings suggest *Bt* may face both synergistic and antagonistic pleiotropies in adapting to the complex metabolic environment of the gut.

We next assessed whether overlapping selection pressures across *in vitro* and *in vivo* environments could drive expansions of the same mutations. To do so, we asked how barcodes with pre-existing adaptive variation in SD mice (Fig. 4E) behaved across our panel of *in vitro* conditions. We found that while the fitnesses within this group of *in vivo-*adaptive barcodes were not correlated between SD mice and any *in vitro* condition, in some complex carbohydrate environments (e.g., Vitafiber, Fig. 6D,E), the fitness of the *in vivo-*adaptive barcodes was on average higher than a noise-matched null subset of barcodes (FDR-controlled *q*<0.05). Conversely, these lineages were collectively enriched for low fitness in monosaccharide environments (e.g., glucose, Fig. 6E). This finding implies that at least some of the pre-existing variation positively selected *in vivo* was present in the *in vitro* experiments, and that the first steps of adaptation to complex sugars faced tradeoffs during growth on simpler sugars, echoing the reduced correlation between SD and MD mice (Fig. 4F).

By aggregating measurements across *in vitro* environments, we were also able to identify individual barcodes with substantial synergies or tradeoffs between *in vitro* and *in vivo* environments. For example, 16 barcodes consistently expanded across replicates in five *in vitro* conditions (Avantafiber, Bimuno, Vitafiber, Vitagos, and Wako) that shared the same barcode inocula (Fig. 6F), which is significantly higher than expected by chance if the *in vitro* environments were independent of each other (*p* = 2 × 10^!#$^; Mann-Whitney U-test against permutated data; Supplementary Text). The fitness rank among these barcodes was reasonably conserved across *in vitro* conditions, suggesting some underlying phenotypic differences between the underlying mutations. However, fitness rank *in vitro* was not highly conserved *in vivo*; several of the fittest barcode lineages *in vitro* were weakly unfit in SD mice. On the other hand, the fittest lineages *in vivo* were unfit *in vitro,* as well as in other host conditions (Fig. 6F, further emphasizing the complexity of the *in vivo* nutritional landscape. These results highlight the utility of high-throughput fitness profiling^7,30,31,56^ to uncover the pleiotropic effects of individual adaptive mutations arising in gut commensals.

To directly compare the genetic determinants of selection in each carbon source with those identified *in vivo*, we performed metagenomic sequencing of passages 0, 1, 15, and 31. In a minority of wells, we were able to identify point mutations that expanded to high frequency in parallel with the growth of specific barcodes (Table S6). Most other adaptations were apparently driven by phase variation, some instances of which (e.g., *BT4520*-*BT4523*) were shared *in vivo* and *in vitro*. Outside of such phase variation, virtually no genetic loci hosting point-like or structural mutations were shared *in vivo* and *in vitro*.

Together, these experiments suggest that *in vitro* evolution in single carbon sources can partially identify selection pressures faced by *Bt* during mouse colonization. On the other hand, we detected limited overlap in the mutations that ultimately emerged to high frequencies *in vivo* versus *in vitro*. This discrepancy may be consistent with *in vivo* evolutionary dynamics that must navigate a complex landscape of pleiotropy^7^ driving fitness tradeoffs (Fig. 6E,F) across carbon sources, complexities which are obviated in the context of a single carbon source *in vitro*. It remains to be seen whether more complex *in vitro* environments, for instance by combining carbon sources, sequential passaging through different carbon sources, or spatially structuring of the *in vitro* environment^57^, better recapitulate *in vivo* selection pressures.

## Discussion

Mounting evidence has highlighted the potential for rapid within-host evolution of the gut microbiota, but the impact of these dynamics on longer term ecological and evolutionary processes across hosts is still unknown. Here, we leveraged high-resolution lineage tracking of ∼60,000 barcoded strains of a prominent gut commensal to quantify the interplay among local adaptation, inter-host transmission, and colonization resistance across multiple spatiotemporal scales.

The high rates of within-host evolution we observed (Fig. 1,4,5) were broadly consistent with previous studies of *Bacteroides* evolution *in vivo*^5–7,58^, but our lineage-tracking approach enabled a more comprehensive statistical view of the evolutionary landscape driving this adaptation. In particular, by exposing our *Bt* libraries to low- and high-diversity limits of community composition, we were able to shed light on the degree to which niche pre-emption (ecological control) or niche creation (diversity begets diversity) shapes the adaptive landscape. By comparing the fitness of pre-existing variants across these two extremes of community diversity, we found that similar rates of evolution at the population level (Fig. 4D) were accompanied by broad differences in the underlying adaptative landscape (Fig. 4E-G), indicating that niche pre-emption and niche creation both play an important role. Our finding that the overall rate of evolution is similar in different community contexts is consistent with previous observations in *E.coli*^10,17^, and is in line with theoretical predictions showing that the rate of adaptation in large populations is only weakly dependent on the population size and the supply of beneficial mutations^59^. These results suggest that broader statistical characterizations of the adaptive landscape – similar to those we have employed here – will be critical for understanding how community composition impacts the evolution of a focal species.

Our analysis should serve as a roadmap for a more thorough exploration of these effects, for instance across a larger panel of diverse communities or across a gradient of community complexities. By introducing distinct pools of barcoded strains into co- and cross-housed mice, we were able to measure transmission and engraftment of *Bt* strains across hosts and test longstanding hypotheses related to the origin and maintenance of colonization resistance in the gut. Our high-resolution view allowed us to distinguish between transient and engrafted lineages: while large numbers of cells were non-specifically transmitted into recipient hosts (presumably by coprophagy), we found that shared selection pressures could enable the rapid spread of specific genetic variants across hosts, overcoming the inherent colonization resistance of the resident strain. These results raise the intriguing possibility that the colonization priority effects that have previously been observed in *Bacteroides* species^34,60,61^ could be driven by rapid adaptation of the primary colonizer to its host environment^62^. On the other hand, we also found that continual transmission, and the emergence of rare but strongly adaptive mutations, may nonetheless override such priority effects, evolved or otherwise (Fig. 2E,F). These results suggest that the colonization resistance landscape may be much more complex than previously appreciated, particularly if the secondary colonizer has also been evolving within a host.

The broad range of transmission rates we observed for mono-colonized *Bt* (∼0.1%-3% per day) was nonetheless substantially lower than we previously observed for *E. coli* in similar housing conditions (∼10% per day)^9^. We hypothesize that the lower transmission of *Bt* was driven by physiological differences with *E. coli* (e.g., sensitivity to oxygen) that directly impact survival outside the gut; if so, the lower transmission rates we observed in a complex community (Fig. 5D) relative to mono-colonized mice (Fig. 2A,C) could be driven by the tendency for Bacteroidetes species to localize on the periphery of fecal pellets in humanized mice with a diverse microbiota^38^, entailing greater exposure to the aerobic environment when outside the host. The variation in transmission over time and across hosts may, in turn, reflect host variation in propensity and/or preferences in coprophagic behavior. Since the rate of transmission sets an upper bound on the rate of engraftment, hosts with higher rates of incoming transmission may be more susceptible to turnover and replacement of resident strains by external population reservoirs. On the other hand, we found that amplification of barcode lineages across hosts was limited not by dispersal but by acquisition of specific adaptive mutations. More experiments will be necessary to determine if such temporal and inter-host variation in transmission plays a meaningful role in shaping lineage abundances across a metapopulation of hosts, or whether the long-term winners are dictated by stochasticity in the sampling of the adaptive landscape within individual hosts. Finally, while non-engrafting lineages (Fig. 2I,J) might appear to be an evolutionarily unimportant subpopulation, these transient strains could still serve as important reservoirs of horizontal gene transfer with the resident gut microbiota^63^.

A key limitation of lineage tracking methods is that they do not directly reveal the genetic targets of adaptation. We overcame this limitation via extensive isolate-based sequencing, revealing diverse modes of mutation in mono-colonized mice, from simple point mutations, to invertible repeat-mediated phase variation, to large scale structural variants that increased the size of the genome by ∼10%. The latter adaptive amplifications (“gene accordions”) are frequently observed as rapid responses to acute stresses, that will be subsequently collapsed under RecA-mediated gene loss^64,65^. Nonetheless, ancient amplifications have also survived in modern *Bacteroides* species (including *Bt*), with the constituent copies having diversified to carry out unique functions^66^. Thus, rapid structural responses to short-term selection pressures may play an important role in setting the genomic basis for long-term evolution and diversification. These and other mutations evolved repeatedly across *in vivo* experiments, sometimes in multiple lineages in the same cage, reflecting the specificity of selection pressures for particular variants as well as the capacity of large *in vivo* populations to densely sample the adaptive landscape. In addition to isolate sequencing, we also leveraged the existence of standing genetic variants to shed light on the diversity, phenotypic consequences, and environmental specificities of adaptive mutations across *in vivo* and *in vitro* environments (Fig. 6D-F). This approach could be extended in future work by transplanting evolved populations with a high diversity of adaptive mutations^7,30,31,67^ (e.g., around day 10 in our *in vivo* experiments) into a large panel of *in vitro* or *in vivo* environments. Such comparisons can directly address the phenotypic or pleiotropic effects of mutations without the need for large-scale whole genome sequencing efforts^7^, which are especially onerous when studying focal species embedded as a minority within a diverse communities.

These results show that high-resolution lineage tracking can be a powerful tool for quantifying eco-evolutionary dynamics in complex *in vivo* environments. Our barcoded *Bt* library, along with future extensions to other gut commensals, should find a wide range of applications in interrogating host-associated evolution across multiple intra- and inter-host scales.

## Methods

### Mice

All mouse experiments were conducted in accordance with the Administrative Panel on Laboratory Animal Care, Stanford University’s IACUC. Experiments involved female Swiss-Webster mice 6-12 weeks of age. For experiments involving *Bt* libraries, each well was used to inoculate 1 mL of TYG + 25 mg/mL erythromycin and grown for 24 h. Community members were inoculated into 1 mL of Brain Heart Infusion medium supplemented with hemin, L-cysteine, tryptophan, arginine, and vitamin K3 (BHI-S) and grown for 24 h. One hundred microliters of the overnight culture of each community member were mixed to construct the community inoculum lacking *Bt*. This inoculum was combined at a 1:1 ratio with a barcoded *Bt* pool for community co-colonization experiments.

For *E. coli* and *Bt* co-colonization experiments (Fig. S8), mice were gavaged with 10^8^ cells of an equal mixture of *E. coli* barcoded strains and 10^8^ cells of *Bt* labeled with cytoplasmic GFP. SD mice were fed a standard diet of normal mouse chow (Purina LabDiet 5010) *ad libitum* throughout the experiment. For experiments involving a dietary switch, mice were first fed a standard diet (Purina LabDiet 5010) rich in MACs, and then a defined low-MAC diet (Teklad TD.150689) in which the sole carbohydrate is glucose (63.5% (w/v)).

Mice were euthanized with CO_2_ and death was confirmed via cervical dislocation.

### Bacterial strains and culturing

*E. coli* MG1655 wild-type and evolved strains were grown in LB and incubated aerobically, shaking orbitally (225 rpm) at 37 °C. *Bt* and related strains were grown by streaking out glycerol stocks on Brain Heart Infusion plates supplemented with 10% defibrinated horse blood and the appropriate antibiotics (200 mg/mL gentamycin for *Bt*, 25 mg/mL erythromycin for barcoded strains), and after 24-48 h of incubation in an anaerobic chamber (85% N_2_, 10% CO_2_, 5% H_2_) at 37 °C, a single colony was selected and grown anaerobically at 37 °C for 24-48 h in liquid tryptone-yeast extract-glucose (TYG) medium (10 g of Bacto Tryptone (Gibco, Cat. #211705), 5 g of yeast extract, 2 g of glucose, 0.5 g of L-cysteine, 100 mL of 1 M KPO_4_ [pH 7.2], 1 mL of 1 mg/mL vitamin K_3_ solution, 40 mL of TYG salts solution (0.5 g MgSO_4_·7H_2_O, 10 g NaHCO_3_, 2g NaCl in 1 L of H_2_O), 1 mL of 0.8% CaCl_2_ solution, and 1 mL of 0.4 mg/ml FeSO_4_ combined in a total volume of 1 L, autoclaved, and supplemented with 1 mL hematin-histidine solution (12 mg hematin dissolved in 10 mL of 0.2 M histidine [pH 8])) supplemented with the appropriate antibiotics (25 mg/mL erythromycin).

### Quantification of bacterial densities

*Bt* densities were quantified by spot plating on duplicate BHI agar plates supplemented with 10% defibrinated horse blood. Plates incubated at 37 °C in an anaerobic chamber (Coy Laboratory Products). Plates were incubated aerobically at 37 °C for 24 h.

### Barcode library creation

We assembled two vectors (pWW3808: split Amp^R^ for *E. coli*, Erm^R^ for *Bacteroides*, R6K origin, strong promoter; and pWW3810: terminator, NBU integrase, split Amp^R^ for *E. coli*) and a random 26-nucleotide DNA barcode (**NNNNN**TT**NNNNN**AC**NNNNN**AA**NNNNN**, where the Ns represent random nucleotides) into a barcoding vector pool using BsaI. A mixture of double-stranded random DNA barcode sequences was ordered from IDT at a concentration of 100 µM. These three components were mixed equimolarly and assembled using a Golden Gate reaction according to standard procedures^68^.

One microliter of the Golden Gate reaction was mixed with 40 µL of electrocompetent S17-1 *E. coli* cells. Cells were transformed using 1-mm cuvettes at 1.8 kV, and 960 µL of LB were added immediately for cell recovery. For each electroporation, 100 µL of recovered cells were plated on LB supplemented with 150 µg/mL carbenicillin and grown overnight aerobically at 37 °C. A total of 20 electroporations were performed and then split into 192 pools, with the expectation that each electroporation would result in ∼200-400 colonies based on pilot experiments. On the same day, a single *Bt* VPI-5482 colony was used to inoculate a 200-mL liquid culture that was grown overnight anaerobically in TYG medium.

The next day, *E. coli* colonies were scraped from each of the 192 plates into an individual pool, resuspended in 0.5 mL of LB, and mixed 1:2 with the overnight *Bt* culture. Each mixture was spun down, resuspended in 100 µL of LB, plated on BHI agar plates supplemented with 10% horse blood, and grown aerobically for 24 h at 37 °C. Growth of the transformed *E. coli* depletes the oxygen so that *Bt* can grow anaerobically beneath the *E. coli* biomass and serve as the conjugation recipient. After 24 h, cells were scraped, added to 1 mL of LB, and diluted 1:5 before plating on BHI agar plates supplemented with 10% horse blood, 200 mg/mL gentamycin, and 25 mg/mL erythromycin. The gentamycin selects against *E. coli* and the erythromycin selects for *Bt* cells containing the newly integrated sequence. These plates were grown anaerobically for 24 h at 37 °C. Finally, transformed *Bt* colonies were scraped and added to 500 µL of TYG medium supplemented with 25% glycerol. Each of the 192 plates was scraped and maintained separately, resulting in 192 pools of barcoded strains that were split into two 96-well plates and stored at -80 °C.

In preparation for gavage into mice, all frozen stocks of the 192 pools of barcoded strains were grown separately for 24 hours in 1 mL of TYG medium supplemented with 200 mg/mL gentamycin and 25 mg/mL erythromycin. After 24 h of growth, certain collections of wells wells (2 pools of 96 wells each for experiment 1 and 4 pools of 48 wells each for experiment 2) were combined at equal volumes and resuspended in 500 µL of TYG.

### DNA extraction and sequencing

DNA was extracted from whole fecal pellets and bacterial cultures using the PowerSoil-htp and UltraClean 96 microbial kits (Qiagen), respectively. For barcode sequencing, extracted DNA was amplified using a two-stage Phusion High-fidelity DNA polymerase PCR (NEB). In a three-cycle PCR, adapter regions and unique molecular identifier (UMI) barcodes were added to each template. This step generates one uniquely labeled functional template per initial template molecule. The initial primers were then removed using Ampure XP Beads (Beckman Coulter) at a 1:1 ratio. Thirty cycles of a second PCR were used to amplify these labeled templates. During this reaction, known DNA indices were attached to the product to enable informatic demultiplexing of pooled libraries.

The first PCR program was 98 °C for 2 min, three cycles of (98 °C for 15 s, 53 °C for 30 s, 72 °C for 30 s), 72 °C for 5 min, and hold at 4 °C. The second PCR program was 98 °C for 2 min, 30 cycles of (98 °C for 15 s, 69 °C for 30 s, 72 °C for 30 s), 72°C for 5 min, and hold at 4 °C. The primers used for the PCR to add the UMIs are 5’-ACACTCTTTCCCTACACGACGCTCTTCCGATCT**NNNNNNNN**tgcagtgtcgaaagaaaca aa-3’ and 5’-GACTGGAGTTCAGACGTGTGCTCTTCCGATCT**NNNNNNNN**aaatgctgttccatcactgg-3’, where the Ns indicate the UMI. The primers used for the PCR to add the multiplexing indices are 5’-AATGATACGGCGACCACCGAGATCTACAC**NNNNNN**AAACACTCTTTCCCTACA CGACGCTCTTCCGATCT-3’ and 5’-CAAGCAGAAGACGGCATACGAGATAA**NNNNNN**GTGACTGGAGTTCAGACGTGTGCTCTTCCGATCT-3’, where the Ns indicate the multiplexing indices.

Successful reactions were confirmed via agarose gel electrophoresis and pooled in equal volumes. The sequencing library was finalized via purification with Ampure XP Beads at a 1:1 ratio. Sequencing was performed on an Illumina NextSeq with read length 2×146 bp and an average of ∼10^6^ reads per sample.

For metagenomic and whole-genome sequencing, extracted DNA was arrayed into 384-well plates and concentrations were quantified and normalized using a PicoGreen dsDNA quantitation kit (ThermoFisher). DNA was added to a tagmentation reaction, incubated for 10 min at 55 °C, and immediately neutralized. Mixtures were then added to 10 cycles of a PCR that appended Illumina primers and identification barcodes to enable mixing of samples during sequencing. Wells were mixed using 1 µL per well, and the pooled library was purified twice using Ampure XP beads to select the appropriately sized bands. Finally, library concentration was quantified using a Qubit (Thermo Fisher). Sequencing was performed on a NovaSeq S4 with read lengths of 2×146 bp.

### Barcode sequencing analysis

Barcode dynamics were analyzed using *Bartender*^69^ with default parameters. *Bartender* first extracts a list of putative barcodes consistent with the given template sequence (including non-variable spacers). To account for sequencing errors, putative barcodes in this list are clustered into a consensus barcode based on sequence overlap and relative read frequencies, with a maximum cluster distance of 2. Barcode frequencies were estimated by *Bartender* based on barcode read counts and ignoring UMIs. We also developed a custom method leveraging UMIs to infer the effective limit of detection in each sequencing library, which is described in the Supplementary Text. In brief, unique UMI-barcode pairs with sufficient read counts (to rule out sequencing artefacts like template switching) were used to estimate the number of unique templates input to PCR. This number was typically much less than the number of sequenced reads in a library, and thus represented the effective frequency resolution in a library.

## 16S rRNA gene sequencing

16S rRNA amplicons were generated using the Earth Microbiome Project-recommended 515F/806R primer pairs. PCR products were cleaned, quantified, and pooled using the UltraClean 96 PCR Cleanup kit (Qiagen) and Quant-It dsDNA High Sensitivity Assay kit (Invitrogen). Samples were sequenced with 250- or 300-bp reads on a MiSeq (Illumina). Samples were de-multiplexed and analyzed using DADA2^70^ and QIIME v. 1.8^71^ as previously described. Custom MATLAB (MathWorks) scripts were used to analyze amplicon sequence variant (ASV, a proxy for species) distributions^72^. Relative abundances are reported in Table S3.

### *In vitro* passaging

To initiate *in vitro* evolution experiments, the 192 wells in P1 and P2 containing *Bt* barcoded strains were grown separately in 96-well plates for 24 h in TYG + 25 mg/mL erythromycin. The 192 wells were split and mixed at equal volume in 6 pools of 32 wells each. Each pool was washed twice with minimal medium (no carbon source) and resuspended in various minimal media with one carbon source. Each carbon source was inoculated in 96-well plates with two pools, each in duplicate, for a total of four replicates per carbon source. Each well was passaged in a cycle of 1:200 dilution and 48 h of growth, for 31 passages. After every two weeks of passaging, plates were mixed with glycerol (25% final concentration) and stored at -80 °C.

For measuring the growth of evolved populations, frozen stocks of all populations from a given passage as well as the wild-type ancestor were grown in glucose minimal medium for 24 h and then diluted 1:200 into the appropriate carbon source and grown anaerobically for 48 h at 37 °C in a 96-well plate. Optical density at 600 nm (OD_600_) was measured in an Epoch 2 Microplate Spectrophotometer (Biotek Instruments) with continuous orbital shaking. All growth curves are reported in Table S5.

### Picking *Bt* isolates

One microliter of frozen fecal pellet was resuspended in 200 µL of PBS and diluted three times 1:10 in PBS. One hundred microliters of the final dilution were plated on TYG-agar supplemented with 200 mg/mL gentamycin and 25 mg/mL erythromycin and grown anaerobically for at least 48 h at 37 °C. Using a colony picker, approximately 96 colonies per mouse were picked in aerobic conditions individually into 200 µL of TYG medium supplemented with 200 mg/mL gentamycin and 25 mg/mL erythromycin. The cellular resuspension was added to 800 µL of oxygen-depleted TYG medium supplemented with 200 mg/mL gentamycin and 25 mg/mL erythromycin. Cultures were grown anaerobically for 48 h at 37 °C. Cultures were then split in half for glycerol stocks and to isolate DNA for sequencing.

### Preparation of mouse gut contents for metabolomics

Cecum and fecal contents of germ-free mice were collected and quickly stored at -80 °C. Samples were processed as reported previously^10^. Briefly, aqueous extraction (in D_2_O) was performed by homogenization with glass beads in a QIAGEN Tissuelyser II (Retsch) for 2 min with 30 rev/s pulses, followed by two centrifugation steps. The first centrifugation, for removing large debris and glass beads, was at 14000 rpm for 30 min and 4° C, followed by supernatant filtration (0.22-mm filter, Milipore). The second centrifugation was carried out with filtration (3-kDa filters, Vivaspin500) at 15000*g* and 4° C for 3 h or until 150 µL of filtrate was obtained. For acquisition, samples were mixed with phosphate buffer (pH 7) with 2% NaN_3_, 3-(Trimethylsilyl)propionic-2,2,3,3-d4 (TSP-d4, Sigma-Aldrich) as external standard and D_2_O to a total volume of 600 µL and transferred to 5-mm glass tubes for acquisition on a Bruker AVANCE II+ 500 MHz instrument equipped with Cryo TCI (F) (Prodigy) 5-mm probehead with *z*-gradients. ^1^H-NMR spectra were acquired using a 1D NOESY pulse sequence with pre-saturation (noesypr1d) as previously described^10^. Spectra were processed using Chenomx NMRSuite v. 8.1 and compounds were identified by manually fitting reference peaks to spectra in the database Chenomx 500 MHz v. 10. Quantification was based on standard (TSP-d4) peak integration. Metabolomics are reported in Table S4.

### Computational methods and analyses

Models of within-host selection and inter-host transmission and engraftment that were used to interpret or fit to data (Fig. 1,2 and 5), as well as statistical analyses of sequencing data, are described in the Supplementary Text.

### Data and Code Availability

Scripts to generate all figures, along with post-processed barcode and whole genome sequencing data, are available on Github: https://github.com/DanielWongPGH/btheta_barcoded_evolution.

## Acknowledgments

The authors thank the Huang, Good, and Sonnenburg labs for helpful discussions. NMR data were acquired at CERMAX, ITQB-NOVA, Oeiras, Portugal with equipment funded by FCT, project AAC 01/SAICT/2016. The authors acknowledge support from NIH RM1 Award GM135102 and R01 AI147023 (to K.C.H.), NSF Awards EF-2125383 and IOS-2032985 (to K.C.H.), NIH R35 GM146949 (to B.H.G.), Alfred P. Sloan Foundation grant FG-2021-15708 (to B.H.G.), and a Friedrich Wilhelm Bessel Award from the Humboldt Foundation (to K.C.H.). J.L.S., B.H.G., and K.C.H. are Chan Zuckerberg Biohub Investigators. This work was also supported in part by the National Science Foundation under Grant PHYS-1066293 and the hospitality of the Aspen Center for Physics. We thank the Stanford Research Computing Center for use of computational resources on the Sherlock cluster.

