## Supplementary Text and Figures for "High-resolution lineage tracking of within-host evolution and strain transmission in a human gut symbiont across ecological scales"

### Computational methods and analyses

#### *Noise profiling of barcode sequencing with UMIs*

Barcode amplicon libraries were prepared using forward and reverse primers designed with random 8-base unique molecular identifiers (UMIs). In principle, this strategy associates each template barcode with two 8-base UMIs. After PCR and sequencing, UMIs associate paired-end reads with a specific template from DNA extraction. One can then correct for PCR amplification noise (i.e., jackpotting) or amplification biases by de-replicating multiple reads associated with the same barcode and (pair of) UMIs, since these reads were likely derived from the same template[1].

In most of our sequencing libraries, the vast majority of paired reads carried the reverse primer-associated UMI (rUMI), but lacked the forward primer-associated UMI (fUMI). Moreover, in each library lacking the fUMI, we typically found that most reads belonging to a barcode were represented by a much smaller number of rUMIs. That is, most of the library was represented by barcode-rUMI pairs with tens to hundreds of reads (of some characteristic size in each library), indicating PCR jackpotting. This behavior was likely linked to absence/loss of the forward primer, as libraries in which both the fUMI and rUMI were present showed less skew in the barcode-rUMI distribution. We did not seek to determine which step(s) in our library sequencing protocol (from extraction through sequencing) drove fUMI loss, because, as we discuss below, while these issues limited frequency resolution in some samples, their impact could not drive spurious signatures of selection or transmission that are the primary focus of our study.

Nonetheless, we still sought to use the information encoded in the UMIs to estimate the magnitude of technical noise in each library. In libraries carrying paired reads with only rUMIs, it is not straightforward to de-replicate reads associated with high frequency lineages, for which the number of input templates may approach or exceed the diversity of the 8-base UMIs. In this case, de-replicating reads sharing the same UMI but originating from different templates of a barcode would systematically underestimate the counts of high-frequency barcodes, thereby underestimating their true frequencies and boosting the apparent frequencies of low-frequency barcodes[2]. Instead, we sought an approach that could be uniformly applied to all libraries.

To this end, we first defined the estimated frequency of barcode lineage *l* in mouse *m* at time point *t* based on the associated number of reads $R_{l,m}^{(t)}$ as

$$\hat{f}_{l,m}^{(t)} = \frac{R_{l,m}^{(t)}}{\sum_{l'} R_{l',m}^{(t)}} \equiv \frac{R_{l,m}^{(t)}}{D_m^{(t)}}, \tag{1}$$

which could be uniformly applied to all sequencing libraries and was sufficient for most downstream analyses. This metric is an unbiased estimator of the true barcode frequency in the fecal sample, $f_{l,m}(t)$, as long as the probability of PCR jackpotting for a given template is independent of barcode identity.

On the other hand, the amount of technical noise is controlled not by the total reads but by the number of unique templates represented in each sequencing library. In a typical library with total mapped reads $D_m^{(t)}$, barcode-rUMI pairs were associated with either order 1 ($O(1)$) or ≫10 reads. Despite the high frequency of barcode-rUMI pairs with $O(1)$ reads (typically >90% of all unique barcode-rUMI pairs, composing 1-70% of reads), these were almost always associated with barcodes that also had jackpot UMIs. This result suggests that the barcode-rUMI pairs with $O(1)$ reads did not derive from independent templates, but rather from amplification errors in the rUMIs of other "jackpotting" amplicons, e.g., due to template switching during PCR[3]. Since our frequency estimator ignores the UMIs, such errors should not drive substantial biases in the estimated frequencies. On the other hand, this result implied that the effective sampling depth was represented not by the nominal sequencing depth $D_m^{(t)}$, but rather by a smaller underlying number $\widetilde{D}_m^{(t)}$ of templates contributing jackpot reads to the library.

To estimate $\widetilde{D}_m^{(t)}$, we took advantage of the substantial separation of read counts of the "jackpot" UMIs compared to the "noise" UMIs to infer an effective depth based on the total number of “jackpotting” lineages. For each library, we estimated the distribution of reads associated with unique barcode-rUMI pairs among barcodes at low overall frequency (<1%), as estimated by the software package *Bartender*[4] (Methods). We assumed that the number of reads associated with a barcode-rUMI pair was drawn from a zero-truncated Poisson distribution with either a non-jackpot mean $Z_1 \sim 1$ or jackpot mean $\tilde{Z} \gg 1$. In turn, the total read depth $D_m^{(t)} = Z_1\,D_1 + \tilde{Z}\widetilde{D}$, where $\widetilde{D}$ and $D_1$ are the number of barcode-rUMI pairs associated with jackpot or non-jackpot templates, respectively. (We ignore, for brevity, the sample dependence of $Z_1, D_1, \tilde{Z}$, and $\widetilde{D}$). We fit a two-component mixture model, via expectation-maximization, to the empirical distribution to estimate $Z_1, D_1, \tilde{Z}$, and $\widetilde{D}$.

We then modeled the reads associated with barcode $l$ as

$$R_{l,m}^{(t)} \sim \mathrm{Poisson}\left(\left(\frac{Z_1 D_1}{\widetilde{D}} + \tilde{Z}\right) k_{l,m}^{(t)}\right), \tag{2a}$$

$$k_{l,m}^{(t)} \sim \mathrm{Poisson}\left(\widetilde{D} f_{l,m}^{(t)}\right), \tag{2b}$$

where $k_{l,m}^{(t)}$ is the number of templates associated with barcode $l$. While Eq. 1, involving only the nominal sequencing depth, is an unbiased estimator of a barcode’s frequency under this model, the probability of *detecting* a barcode at true frequency $f_{l,m}(t)$ is controlled by $\widetilde{D}$ (when $\frac{Z_1 D_1}{\widetilde{D}} + \tilde{Z} \gg 1$):

$$P\left[R_{l,m}^{(t)} > 0 | f_{l,m}(t)\right] = \exp\left[-f_{l,m}(t)\widetilde{D}\left(1 + e^{-(\frac{Z_1 D_1}{\widetilde{D}} + \tilde{Z})}\right)\right] \equiv \exp\left[-f_{l,m}(t) D_{m,\text{eff}}^{(t)}\right]. \quad (3)$$

In turn, the (inverse of) the effective depth, ${D_{m,\text{eff}}^{(t)}}^{-1}$, sets the scale of the minimum frequency at which a barcode will be detected in a library, although PCR amplification fluctuations (Eq. 2a) mean that barcodes may be measured below ${D_{m,\text{eff}}^{(t)}}^{-1}$. We use the effective depth $D_{m,\text{eff}}^{(t)}$ in downstream analyses and plotting. The exception was for libraries without strong jackpotting, such that $Z_1 D_1 \geq O(1) \cdot \tilde{Z}\widetilde{D}$. In these cases, the majority of singleton barcode-UMIs likely represented unique templates; for simplicity, we assumed $D_{m,\text{eff}}^{(t)} = D_m^{(t)}$ if $Z_1 D_1 \geq 3\tilde{Z}\widetilde{D}$. Fig. S2 shows the agreement between the typical frequency of detected barcode lineages in a library and $1/D_{m,\text{eff}}^{(t)}$ – inferred without any information about lineage sizes – implying our procedure reliably infers the effective sequencing depth in each library. (We note here that many of the barcode-rUMI read count distributions used to fit $Z_1, D_1, \tilde{Z}$, and $\widetilde{D}$ were over-dispersed compared to Poisson. However, properly fitting the amplification variance – e.g., with negative binomial or branching process distributions[5] – was unnecessary to estimate the noise in barcode frequency, since the degree of over-dispersion was always small compared to the reduction in templates, $D_m^{(t)}/\widetilde{D}$.

In principle, barcode-dependent amplification (e.g., if $\tilde{Z}$ depends on the barcode) could bias the estimator in Eq. 1 away from the true frequency. However, these biases could not drive the strongest signatures of selection and transmission—sweeps of the fittest barcodes to frequencies >1% (Fig. 1,2,4-6) and their transmission and engraftment between hosts (Fig. 2,5)—since the consistency of the changes across multiple days are difficult to explain by random and atemporal technical sources. Moreover, our single-day transmission inferences average over barcodes (Fig. 2B), and so should not be strongly influenced by barcode-dependent amplification biases. Finally, the observed environment-dependent fitness correlations across experiments (Fig. 4E-G) would not be expected from technical biases alone.

***Assigning barcode lineages to specific plates/inocula***

A total of 59,205 barcodes were detected in at least two *in vivo* samples (either in two mice or at two time points in the same mouse), or at frequency $>10^{-5}$ in at least one library. Here, we describe our procedure to assign barcodes in the strain library to their respective inocula. We used these inoculum assignments to visualize lineage transmission across mice, and in downstream quantitative analyses of transmission.

*Assigning barcodes to P1 or P2.* We stratified barcodes into P1 and P2 inocula (used to inoculate mice in Fig. 1,2), based on their abundances in the P1 and P2 input libraries prior to gavage, and in the libraries of mice gavaged with P1 or P2. Barcodes were assigned to the P1 inoculum if: (i) they were detected in at least two samples across the

P1 input library and fecal samples from all P1-inoculated mice (5 SD, 3 MD) collected 1-4 days after gavage; and (ii) they were detected in fewer than two samples across the P2 input library and fecal samples from all P2-inoculated mice (5 SD), or were measured at >10-fold higher frequency in the P1 input versus the P2 input library. P2 barcodes were defined reciprocally. This procedure assigned 28,021 and 24,197 barcodes to the P1 and P2 inocula, respectively. The remaining 6,987 barcodes of ambiguous origin represented ~2-3×$10^{-3}$ of P1 or P2 input inocula, and collectively represented ≤1% of the population in all but two mice, in which 1-2 of these barcodes grew above 1% frequency. This ambiguous set could be cases in which the same barcode was involved in multiple lineage-establishing insertion events across different plates or wells; cross-contamination of wells or plates during library creation and/or cultivation prior to evolution experiments; or misidentified barcodes due to contamination during sequencing library preparation, sequencing errors, or index hopping.

*Assigning barcodes to sets 1-5.* For the second set of colonization experiments (Fig. 4,5), we split the barcode library into five distinct sets. Sets 1 and 2 were created by combining non-overlapping wells of P1, sets 3 and 4 were created by combining non-overlapping wells of P2, and the low-diversity set 5 was created by picking ~100 colonies from different wells in P1.

Sets 1 and 2 were only used to inoculate singly housed mice, and did not need to be tracked across hosts. We thus simply assigned a barcode to set 1 or 2 if it was previously assigned to P1 and was measured at least twice in feces from mice inoculated with set 1 or 2, respectively, collected 1-4 days after gavage. This strategy assigned 12,672 and 11,551 distinct barcodes to set 1 and 2, respectively, while 28 barcodes were shared between the sets. All other barcodes collectively represented <3% abundance across mice inoculated with either set 1 or set 2. Thus, we were able to associate set 1 and set 2 with almost entirely non-overlapping sets of P1 barcodes.

Sets 3, 4, and 5 were inoculated into mice that were always immediately co-housed with each other. Thus, unlike our approach to disambiguate P1/P2 and set 1/2 lineages, we could not leverage sequencing libraries from mice exclusively exposed to set 3, 4, or 5. Instead, we first identified 109 barcodes measured at $>10^{-3}$ in two replicate input libraries representing the low-diversity set 5 inoculum. As expected, most (86) of these barcodes were associated with P1, while 23 were ambiguous; 0 derived strictly from P2. We then assigned barcodes with set 3 or 4 if they satisfied the following conditions: (i) they were assigned to P2 or the ambiguous set; (ii) they were not assigned to set 5; (iii) they were measured at >10-fold higher frequency in set 3 compared with set 4, or vice versa. This conditioning was stricter than that used to identify set 1 and set 2 barcodes, and assigned 6,624 and 7,725 non-overlapping barcodes to set 3 and set 4, respectively.

We identified 16 barcodes, not yet assigned to set 3, 4, or 5, that rose to 1-15% frequency in at least one, and typically multiple, co-housed mice in one cage (and in no other cages). Since these adaptive barcodes and their patterns of transmission were of particular interest, we applied additional filtering to disambiguate these barcodes into their respective inocula. For each, we determined the time at which the barcode first reached >1% frequency in a mouse. If at this time the same barcode was measured at <1/10 the frequency in a second mouse, then we rationalized that this adaptive lineage likely spread from the first mouse. We thus assigned the barcode to the inoculum set of the first mouse. This strategy specified 15 of the 16 barcodes to set 3, 4, or 5. The remaining barcode was treated as ambiguous and ignored in downstream analyses of transmission.

*Assigning barcodes to* in vitro *inocula.* For *in vitro* evolution experiments, wells were inoculated with one of six barcode pools, to permit detection of contamination or mislabeled samples. To this end, we first assigned barcodes to an inoculum if it was measured above $10^{-6}$ in the input library or above $10^{-3}$ in at least one library sampled at any time point from a well intended to contain that inoculum. This process designated 6,189-7,838 barcodes across inocula. 658 of these barcodes were shared across at least two inocula, and 34 were measured above $10^{-2}$ in wells with nominally distinct inocula, suggesting some incidental transfer of adaptive barcodes between wells. Each of the 658 ambiguous barcodes was assigned to the inoculum with the most samples in which the barcode was above $10^{-3}$, and appear as mismatched colors in Fig. S17.

**Model of evolutionary dynamics**

Here, we introduce a minimal model of within-host lineage dynamics, neglecting inter-host transmission. We also apply this model to *in vitro* lineage dynamics. In subsequent sections, we introduce more complex models including transmission (Fig. 2,5).

In the simplest version of the model, we assume that barcode lineages within a mouse *m* compete with each other as a well-mixed population. Previous work has shown that selection pressures on *Bacteroides* strains can shift within days after colonization of germ-free mice[6,7], motivating the development of a model of lineage dynamics with time-varying selection. In such a model, the frequencies of rare barcode lineages ($f_{l,m} \ll 1$) are described by a system of coupled stochastic differential equations

$$\frac{\partial f_{l,m}}{\partial t} = \left[s_{l,m}(t) - \overline{X}_m(t)\right] f_{l,m} + \sqrt{\Lambda_m(t) f_{l,m}} \cdot \eta_{l,m}(t) \,, \tag{4}$$

where $\eta_{l,m}(t)$ is a Brownian noise term with mean zero and variance one, and $\Lambda_m(t)$ is the strength of genetic drift in mouse *m* at time *t*. Each lineage *l* has time-dependent fitness $s_{l,m}(t)$, and $\overline{X}_m(t)$ is the instantaneous mean fitness of the population

$$\overline{X}_m(t) = \sum_l s_{l,m}(t) f_{l,m}(t). \tag{5}$$

Eq. 4 is a time-dependent generalization of the branching process model employed in previous *in vitro* lineage tracking studies[8,9]. The time dependence of $s_{l,m}$ and $\Lambda_m$ allows us to account for shifting selection pressures and population bottlenecks, respectively. Time-dependent selection can also implicitly account for intra-barcode diversity, for instance as an adaptive mutation arises and sweeps through its barcode lineage.

*Defining and measuring fitness*. From Eq. 4, we define the relative fitness of a lineage over a time interval $t_1$ to $t_2$ as its log-fold change in average frequency (per time):

$$\chi_{l,m}(t_2; t_1) \equiv \frac{1}{t_2 - t_1} \log \langle f_{l,m}(t_t)\rangle / \langle f_{l,m}(t_1)\rangle = \int_{t_1}^{t_2} dt' \left( s_{l,m}(t') - \overline{X}_m(t') \right). \tag{6}$$

We estimate $\chi_l(t_2; t_1)$ using the shrinkage estimator we previously developed[7]:

$$\hat{\chi}_{l,m}(t_2; t_1) \equiv \frac{1}{t_2 - t_1} \log \frac{\max\left\{\hat{f}_{l,m}^{(t_2)}, \left\{(D_{m,\mathrm{eff}}^{(t_2)})^{-1}, \hat{f}_{l,m}^{(t_1)}\right\}\right\}}{\max\left\{\hat{f}_{l,m}^{(t_1)}, \min\{(D_{m,\mathrm{eff}}^{(t_1)})^{-1}, \hat{f}_{l,m}^{(t_2)}\}\right\}}. \tag{7}$$

When a lineage is measured at both time points, this estimator reduces to $\frac{1}{t_2 - t_1} \log \frac{\hat{f}_{l,m}^{(t_2)}}{\hat{f}_{l,m}^{(t_1)}}$. When a lineage is only measured at one of the two time points, this estimator still allows us to assign a finite and conservative (zero-biased) fitness estimate consistent with the sampling depths at the two time points.

*Collective behavior of low-frequency lineages.* We compared the model in Eq. 4 to our observed *in vivo* lineage dynamics using several metrics of population diversity. We report the dynamics of two common measures, Shannon diversity (Fig. 1D, 4D) and lineage richness (the number of detected barcodes; Fig. S4, S12). Shannon diversity is robust to sequencing depth but is dominated by the behavior of high-frequency barcode lineages representing only a minority of the adaptive variation in the population. Lineage richness is maximally sensitive to low-frequency lineages, but is restricted by sequencing depth, which strongly varied across our samples. Additionally, both measures are dependent on the initial distribution of lineages, which limits comparisons across experiments with non-identical input libraries.

Thus, we sought to additionally measure the mean fitness of the population $\overline{X}_m(t)$ in Eq. 4, which controls the rate of neutral diversity loss in the population. Related methods to estimate the mean fitness over time in barcode lineage-tracking experiments have been developed[8-10]. These methods were designed with *in vitro* serial dilution experiments in mind, and assume a fixed environment and selection pressure ($s_{l,m}(t) = s_{l,m}$). However, *in vivo* selection pressures in mice appear to change over the time scales of our experiments[7,11,12]. These complications, as well as large and variable amounts of technical noise, limited the utility of previous methods to our data in the present study.

We instead took a cruder but more robust approach to measure the time-integrated mean fitness of the population, $\int \overline{X}_m(t)dt$. The estimate of $\chi_{l,m}(t;0)$ from a single lineage, $\hat{\chi}_{l,m}^{(t,0)}$, will be noisy as well as biased if the lineage is not neutral. Instead, we calculate the median among all lineages with initial frequencies in a narrow frequency range $\hat{f}_{l,m}^{(0)} \in (a_0, b_0)$:

$$\tilde{\chi}_{(a_0,b_0),m}^{(t)} = \text{median}\left\{\hat{\chi}_{l,m}^{(t,0)}; a_0 < \hat{f}_{l,m}^{(0)} < b_0\right\}. \quad (8)$$

Measuring across many barcode lineages averages over the uncertainty associated with a single lineage, and the median is robust to the inclusion of a minority of adaptive lineages. Fig. 1E reports the median and interquartile range of $\exp[t\tilde{\chi}_{(a_0,b_0),m}^{(t)}]$ for all barcodes in sets of non-overlapping $(a_0, b_0)$ spanning a total frequency range $4 \times 10^{-5}$ to $6 \times 10^{-4}$ (6000-7000 barcodes in the upper quartile of input frequencies, representing ~75% of cells in the inoculum). Fig. S12 reports similar calculations for the second cohort of mice (Fig. 4).

*Comparison to simulated data*. We compared the observed trajectory of diversity statistics to Wright-Fisher simulations corresponding to the well-mixed dynamics in Eq. 4. We considered two distinct scenarios. The first corresponded to a neutral population with a conservatively strong[7] drift strength $\Lambda_m(t) = 10^{-6}$/day. In the second, we treated the lineages highlighted in Fig. 1C as a deterministic population of total relative abundance size $F_{\text{fit}}(t)$, and simulated neutral dynamics among the remaining lineages with increasing drift strength $\Lambda_m(t) = \frac{10^{-6}}{1-F_{\text{fit}}(t)}$/day. In both cases, barcode sequencing was simulated as a single Poisson sampling step with an effective depth $D_{\text{eff}}$; introducing additional, variance-increasing steps did not meaningfully change our results. We then recalculated the diversity statistics using these simulated frequencies and compared to observations (Fig. 1D,E, S4, S12).

The discrepancy between the simulated high-frequency expectation and experimental observations implies the presence of a larger number of adaptive lineages than those highlighted in Fig. 1C, 4C. A crude lower-bound is the minimum number of plausibly adaptive barcodes – i.e., those with the largest frequencies at $t$ and with positive fold-change from 0 to $t$ – that can account for the observed $\tilde{\chi}^{(t)}_{(a_0,b_0),m}$. For example, carrying out this procedure on day 9 implies >200 adaptive barcodes in each cage in Fig. 1.

*Measuring fitnesses of high-frequency lineages*. In the main text, we report the range of growth rates among the high-frequency lineages highlighted in Fig. 1C. To generate this range, we estimated the relative fitness, Eq. 7, of each lineage $l$ in mouse $m$ using the minimum time $t_2$ such that $\hat{f}^{(t_2)}_{l,m} > 1\%$ and $t_2 \leq 21$ days, and the maximum time $t_1 < t_2$ such that $\hat{f}^{(t_1)}_{l,m} < 0.1\%$. This procedure allowed us to crudely estimate the exponential growth phase of a lineage (after establishment of an adaptive mutation). Since we do not use the same intervals for every barcode, these measurements do not approximate the relative fitnesses among high-frequency barcodes, but rather the fitness relative to the population at the time of maximal growth for each barcode. Summary statistics are reported for the 70 and 16 lineages satisfying these criteria in at least one SD or MD mouse, respectively. For barcodes reaching $> 1\%$ in multiple mice, we used the median growth rate.

*Using lineages with pre-existing variation to measure fitness across environments*. In the main text (Fig. 4, 6), we leveraged lineages with pre-existing genetic variation – commonly observed in neutral barcoding experiments[9,13] – to probe the joint distribution of fitness effects (JDFE) across *in vivo* and *in vitro* environments.

We established the existence of *in vivo*-adaptive pre-existing mutations in our library based on the common behavior of a lineage in separate experiments sharing the same host conditions. Each half of the P1 barcodes colonizing mono-colonized SD mice (Fig. 1C, middle row) were also introduced to singly housed mice (Fig. 4), which allowed us to compare their relative frequency trajectories across both experiments. Since the similarities between these trajectories can be obscured by sequencing noise, we focused on the subset of 2896 lineages with (i) $\hat{f}_{l,m}^{(3)} D_{m,\mathrm{eff}}^{(3)} > 20$ in the singly housed mono-colonized mice in Fig. 4 ($\hat{f}_{l,m}^{(3)} D_{m,\mathrm{eff}}^{(3)} > 20$), and (ii) $R_{l,\mathrm{cage}}^{(2)}$>20 reads in the P1 mono-colonized mice in Fig. 1, where we define the cage-wide read count as

$$R_{l,\mathrm{cage}}^{(t)} = \hat{f}_{l,m}^{(t)} D_{m,\mathrm{eff}}^{(t)}. \tag{9}$$

In the absence of pre-existing variation, the same lineage should exhibit uncorrelated behavior in separate experiments. We tested this hypothesis by comparing the measured fitness of lineages over the next approximately one week (days 3-8 or 2-9; the intervals vary due to differences in sampling times and/or to minimize technical noise). For the P1-colonized (Fig. 1) mice, we input noise-weighted cage-wide frequencies $f_{l,\mathrm{cage}}^{(t)} = R_{l,\mathrm{cage}}^{(t)} / D_{\mathrm{cage}}^{(t)} \equiv R_{l,\mathrm{cage}}^{(t)} / \sum_l R_{l,\mathrm{cage}}^{(t)}$ in our estimator, Eq. 7.

Conditioning on the expansion of these lineages in the cohort of mice in Fig. 1, $\hat{\chi}_l(9;2) > 0$, we found strong correlation with their measured fitnesses over a similar interval in the second cohort $\hat{\chi}_l(8;3)$ (Fig. 4E). We termed these 718 lineages the standard-diet, mono-colonized fit (SMF) lineages. We then measured SMF lineage fitness in the cage of mono-colonized MD mice from days 2-9 (Fig. 1, bottom row); in community colonized mice (from days 3 to 8 or 9); and *in vitro* from passages 1 to 5. These comparisons are shown in Fig. 4F,G and 6D,E, and discussed in the main text.

Fig. S13 also reports the full JDFE in a larger number of (noisier) barcodes across multiple pairs of environments, unconditioned on high fitness. These comparisons show strong correlation among low-fitness barcodes in the SD mono-colonized environment, indicating strong purifying selection on deleterious pre-existing variation. Additionally, we observed measurable correlations between co-colonized and *Bt* delay-colonized conditions, consistent with shared selection pressures from the presence of the community that are not entirely dependent on the timing of *Bt* colonization.

*Constructing noise-matched null sets of lineages for comparison*. Despite broad de-correlation in fitness among the SMF lineages across non-identical environments (Fig. 4F,G), we sought to test whether the SMF lineages were still enriched for high fitness compared to the larger population. To do so, we compared the SMF lineages to a non-overlapping subset of lineages of the same size, with the same distribution of initial frequencies in the non-identical environment to control for variable strengths of drift. We report the results of one-sided Kolmogorov-Smirnov tests (Fig. 4E, 6D) and the difference in mean relative fitness (Fig. 6E) between the SMF and noise-matched null lineages. In Fig. 6E, we report significant FDR-corrected $p$-values ($q$-values) for a positive difference in mean by first computing two-sided $p$-values in individual wells under a paired sample $t$-test, and then combining $p$-values across replicate wells (identical carbon sources) using Fisher's method. Carbon sources with $q < 0.05$ are bolded, while error bars represent $\pm 2$ SEM from $10^4$ bootstrapped resamplings of the SMF and noise-matched null lineages.

In Fig. 6F, we also identified individual, strongly adaptive lineages *in vitro* as those barcodes that consistently expanded over passages 1 to 5 across $r$ wells ($\hat{\chi}_{l,e_1}(5;1)$>0 for wells $e_1, \dots e_r$). Since we consider thousands of lineages, it is important to generate a null expectation of false positives. To this end, we created fake barcode lineages $l'$ by permuting barcode identities within each well while preserving the overall distribution of initial (passage 1) frequency vectors across wells, $\mathbf{F}_{\boldsymbol{l}}^{(1)} = \left(\hat{f}_{l,e_1}^{(1)}, \hat{f}_{l,e_2}^{(1)} \dots \hat{f}_{l,e_r}^{(1)}\right)$. That is, for each $\mathbf{F}_{\boldsymbol{l}}^{(1)}$, we generated a corresponding $\mathbf{F}_{\boldsymbol{l'}}^{(1)}$ where $\max_{e_i}\left|\log_2\left(\frac{\hat{f}_{l,e_i}^{(1)}}{\hat{f}_{l',e_i}^{(1)}}\right)\right| < 0.5$ for >98% of lineages. We used these scrambled barcodes $l'$ to generate a noise-matched distribution of fitness profiles, $\boldsymbol{X}_l'(5;1) = \left(\hat{\chi}_{l',e_1}(5;1),\ \hat{\chi}_{l',e_2}(5;1), \dots \hat{\chi}_{l',e_r}(5;1)\right)$ that would be expected in the absence of pre-existing variation. We then measured the number of fitness profiles with $\hat{\chi}_{l',e_i}(5;1)$>0 for all $e_i$ across many permutations. Fig. 6F provides an example of this procedure for $r = 9$, representing 5 distinct environments: while we observed 16 barcodes that consistently expanded across wells, 0 were identified across 100 permutations.

***Model of transmission and engraftment of barcodes***

Here, we introduce a model of joint intra- and inter-host lineage dynamics that is consistent with the diversity of barcode lineage trajectories observed across hosts in Fig. 2, 5. We then describe our approach to infer model parameters from data.

In brief, we generalize Eq. 4 above by assuming that the barcode population in each mouse resides in *R* distinct, individually well-mixed niches. Every niche contributes to a well-mixed transient population that combines with newly arriving cells (presumably via coprophagy) to constitute the daily passaging population represented in feces. We assume that turnover of cells in a niche is dominated by engraftment from the (much larger) daily passaging population, and neglect inter-niche migration.

In more detail, we denote the frequency on day *t* of a focal barcode *l* in niche *k* in mouse $m$ by $\phi_{l,k,m}^{(t)}$, and its respective frequency in the feces by $f_{l,m}^{(t)}$. For clarity, we define the model in discrete time and neglect demographic noise. The dynamics in mice are then given by

$$\phi_{l,k,m}^{(t+1)} = \left(1 - g_{k,m}^{(t)}\right)\Psi_{l,k,m}^{(t)}\phi_{l,k,m}^{(t)} + g_{k,m}^{(t)}\gamma_{l,k,m}^{(t)}f_{l,m}^{(t)}, \qquad k = 1, \ldots, R \tag{10a}$$

$$f_{l,m}^{(t+1)} = \left(1 - \tau_m^{(t)}\right)\Upsilon_{l,m}^{(t)}\sum_k \nu_{k,m}^{(t+1)}\phi_{l,k,m}^{(t+1)} + \tau_m^{(t)}\kappa_{l,m}^{(t)}\bar{f}_l^{(t)}. \tag{10b}$$

Eq. 10a reflects the dynamics in a resident niche *k*: a fraction $g_{k,m}^{(t)}$ of the resident population is replaced by engrafting cells arriving from the daily transient population. Thus, $g_{k,m}^{(t)}$ represents the engraftability of the niche. Barcodes may differ in their competition- and engraftment-related (exponential) fitnesses, $\Psi_{l,k,m}^{(t)}$ and $\gamma_{l,k,m}^{(t)} \geq 0$ (with $\sum_l \Psi_{l,k,m}^{(t)}\phi_{l,k,m}^{(t)} \equiv 1$ and $\sum_l \gamma_{l,k,m}^{(t)}f_{l,m}^{(t)} \equiv 1$), owing to mutations accumulated over time. In

Eq. 10b, each niche contributes a fraction $\nu_{k,m}^{(t)}$ ($\sum_k \nu_{k,m}^{(t)} \equiv 1$) to the daily passaging population, which mixes with a transmission fraction $\tau_m^{(t)}$ of cells from the feces of the metapopulation of $M$ hosts, in which the average frequency of $l$ is $\overline{f}_l^{(t)} = \frac{1}{M}\sum_m f_{l,m}^{(t)}$. Analogous to the niche-specific values of $\Psi_{l,k,m}^{(t)}$ and $\gamma_{l,k,m}^{(t)}$, the parameters $\Upsilon_{l,m}^{(t)}$ and $\kappa_{l,m}^{(t)}$ are the growth- and transmission-related fitnesses of the lineage in the transient (fecal) population.

In general, the fitnesses, transmission and engraftment rates, and niche sizes may vary across mice and over time, due to variation in the host environment, barcode lineage composition, and/or the larger microbial community. Below, we consider simplifications of this model to enable testable predictions of our data.

*Measuring transmission across co-housed mice.* We used mice inoculated with distinct barcode populations to measure transmission rates immediately upon cross-housing. We denote the day of cross-housing as $t = t_0$ (with sampling at $t_0$ immediately prior to cross-housing if $t_0 > 0$). For each barcode lineage, we define the donor mice in a cage with $M$ mice as those that were colonized with the inoculum including that barcode, and the recipient mice as those colonized with a different inoculum. In Fig. 2 ($t_0 = 14$), for each lineage, there are two donors and two recipients, and in Fig. 5 ($t_0 = 0$), there is a single donor and two recipients. For a barcode $l$, its resident and transient frequencies in a donor $d$ are $\phi_{l,k,d}^{(t_0)}$ and $f_{l,d}^{(t_0)}$, and in a recipient $r$ are $\phi_{l,k,r}^{(t_0)} = f_{l,r}^{(t_0)} = 0$. Rather than the resident frequency, we are only able to track the transient frequency $f_{l,m}^{(t)}$ of each barcode in feces. At $t_0 + 1$, the transient frequency in a recipient will reflect transmission from the donors over the previous day. It is straightforward to show that the fitness-weighted transmission is given by the ratio of recipient to donor frequencies:

$$\tau_r^{(t_0)} \kappa_{l,r}^{(t_0)} = \frac{f_{l,r}^{(t_0+1)}}{\frac{1}{M}\sum_d f_{l,d}^{(t_0)}}. \tag{11}$$

In principle, substituting different barcodes in Eq. 11 could reveal barcode-specific transmission rates, for instance reflecting variability in survival outside of the host. However, Fig. 2B shows that, owing to low rates of transmission relative to sequencing noise, most detected barcodes are not measured at their true frequency in the recipient, but rather at the effective limit of detection. Hence, the rate of transmission of an individual barcode is not well estimated by the single shot-measurement of Eq. 11. Instead, we assumed that donor barcodes have the same rate of transmission into a given mouse, and any apparent variation is driven by sequencing noise. Under our sequencing model in Eq. 2, the maximum likelihood estimate $\hat{\tau}_r^{(T)}$ is given by the weighted average

$$\hat{\tau}_r^{(T)} = \frac{\sum_l \hat{f}_{l,r}^{(t+1)}}{\sum_l \bar{f}_l^{(t)}}. \tag{12}$$

This measure is most sensitive to the behavior of high-frequency barcodes, which are anomalously fit in the donor mouse. To test the assumption that lineages did not differ in their transmission based on their frequency (potentially reflecting their underlying within-host fitness differences), we also estimated $\tau_m^{(t)}$ based on the detection probability of individual barcodes. This procedure more strongly weights the behavior of typical, low-frequency barcode lineages. From Eq. 2, the probability of detecting an individual barcode in the recipient *r* is

$$P\left(\hat{f}_{l,r}^{(t+1)}>0\middle|\bar{f}_l^{(t)}\right) = 1 - P\left(\hat{f}_{l,r}^{(t+1)}=0\middle|\bar{f}_l^{(t)}\right) = 1 - \exp\left(-\tau_r^{(t)} D_{r,\text{eff}}^{(t+1)} \bar{f}_l^{(t)}\right). \quad (13)$$

We maximized the log-likelihood of the data with respect to the joint parameter $\tilde{\tau}_r^{(t)} = \tau_r^{(t)} D_{r,\text{eff}}^{(t+1)}$,

$$\hat{\tilde{\tau}}_r^{(t)} = \text{argmax}_{\hat{\tilde{\tau}}_r^{(t)}} \sum_{\hat{f}_{l,r}^{(t+1)}=0} \left(-\tilde{\tau}_r^{(t)} \hat{f}_{l,d}^{(t)}\right) + \sum_{\hat{f}_{l,r}^{(t+1)}>0} \log\left(1 - e^{-\tilde{\tau}_r^{(t)} \hat{f}_{l,d}^{(t)}}\right). \quad (14)$$

We then estimated $\hat{\tau}_r^{(t)}$ as $\hat{\tilde{\tau}}_r^{(t)}/D_{r,\text{eff}}^{(t+1)}$. The results in Fig. 2B are consistent with a uniform transmission rate across frequencies.

*Transmission after the first day of cross-housing*. We also sought to estimate rates of ongoing transmission multiple days after cross-housing (Fig. 2C, 5D, S5, S14). Beyond the first day of cross-housing, the frequency of *l* in the recipient can reflect both transmission and engrafted cells from previous transmission events. Two days after cross-housing,

$$f_{l,r}^{(t_0+2)} = \left(1 - \tau_r^{(t_0+1)}\right) \Upsilon_{l,r}^{(t_0+1)} \sum_k \nu_{k,r}^{(t_0+2)} \underbrace{g_{k,r}^{(t_0+1)} \Upsilon_{l,k,r}^{(t_0+1)} f_{l,r}^{(t_0+1)}}_{=\phi_{l,k,r}^{(t_0+2)}} + \tau_r^{(t_0+1)} \bar{f}_l^{(t_0+1)}, \qquad (15)$$

where we continue to assume that transmission is lineage independent. To measure the rate of transmission alone beyond $t_0$, we focus on barcode lineages unable to engraft in recipient mice, i.e. $g_{k,r}^{(t)} \gamma_{l,k,r}^{(t)} = 0$ for every niche *k*. Note that our model does not distinguish whether a barcode cannot colonize a niche because the niche is uninvadable after colonization ($g_{k,r}^{(t)} = 0$), or because it has strongly negative engraftment fitness in the niche ($\gamma_{l,k,r}^{(t)} = 0$). For these barcodes, their frequency in recipient hosts is

$$f_{l,r}^{(t+1)} = \tau_r^{(t)} \bar{f}_l^{(t)}, \qquad (16)$$

for all times $t \geq t_0$. We did not know *a priori* which barcodes were unable to engraft in recipient mice. Thus, for each recipient mouse and each transmission interval $(t_1, t_2)$, we considered the (donor) barcodes that were at substantial frequency in the donor, $f_{l,d}^{(t_1)} > 2/D_{\text{eff},d}$, and undetected in the recipient, $f_{l,r}^{(t_2)} = 0$, as a candidate set of non-engrafting barcodes. In the cross-housing scenario with two donors (Fig. 2), we condition on the average donor frequency being larger than the minimum value of $D_{\text{eff},d}^{-1}$ among the donor mice that were sampled at time $t_1$. We used this set of barcodes to estimate transmission using both maximum likelihood estimates (Eq. 15, 16), which were broadly correlated in all experiments (Fig. S5, S14). When mice were not consistently sampled at single day intervals, we estimated donor frequencies $f_{l,d}(t-1)$ prior to detection in the recipient at *t* using $\hat{f}_{l,d}(t - \Delta t)$ with $\Delta t$=2 or 3. This strategy should minimally bias transmission measurements relative to the apparent scale of genuine day-to-day variation, as most lineage frequencies changed less than two-fold in donor mice over 2-3 day intervals.

In principle, previously engrafted barcodes that went undetected at the first time point due to finite sequencing depth could more substantially bias our inferences. However, our results are most consistent with low rates of engraftment ($g \ll 1$) for most barcodes. For instance, in Fig. 2C, the increasing recipient-to-donor ratio across barcodes detected in mouse 6 over days 15 to 19 might naively suggest that ongoing transmission is adding to an engrafted subpopulation. However, this ratio increased by over an order of magnitude over the short interval, which is quantitatively inconsistent with the measured rate of transmission over the first day of cross-housing and the strength of positive selection. Moreover, the recipient-to-donor ratio in mouse 4 declined over days 16 to 18, clearly inconsistent with engraftment and long-term persistence. At later times, inferred transmission rates were approximately stable in an individual mouse, especially compared to the variation across mice. This observation is also inconsistent with the expectation that lineages transmitting and successfully engrafting should approach similar frequencies in the donor and recipient (compare to Fig. 5C). Instead, it is consistent with roughly balanced replacement with new migrants and wash out, in which case our measure would reflect a weighted average of transmission over the previous days. However, the strong fluctuations we observed suggest the measure is dominated by transmission over single days.

*Predicting barcode frequencies over time*. After inferring transmission, we sought to distinguish genuine engraftment from ongoing transmission using the frequency trajectories of donor barcodes in recipient hosts. To this end, we considered barcode lineage behaviors at two limits of engraftment in our model. In the non-engraftment limit $g_{k,r}^{(T)}\gamma_{l,k,r}^{(T)} = 0$ (for every niche *k*) described above, the frequency of a barcode in a recipient mouse is described by Eq. 16. This theoretical expectation of the "transmission floor" is plotted for individual barcodes in Fig. 2E-J, 5E-J, using transmission inferences from Fig. 2C, 5D, S5, S14.

We next consider a class of engrafting barcodes for which we can make straightforward predictions of dynamics in recipients. This class strictly colonizes niches with complete daily turnover, $g_{k,m}^{(t)}$ =1 for all $\gamma_{l,k,r}^{(t)} > 0$. For instance, these barcode lineages could colonize niches with no spatial distinction from the transient population (e.g., in the lumen). As in the case of non-engrafting barcodes, the dynamics simplify to an effective one-compartment model involving only the observable fecal frequency $f_{l,m}^{(t)}$:

$$f_{l,m}^{(t+1)} = \left(1-\tau_m^{(t)}\right)\Upsilon_{l,m}^{(t)}\sum_k \nu_{k,m}^{(t+1)}\gamma_{l,k,m}^{(t)} f_{l,m}^{(t)} + \tau_m^{(t)}\bar{f}_l^{(t)}. \quad (17)$$

We sought to predict the trajectory of a barcode in recipient mice, given only its frequency $f_{l,d}^{(t)}$ in the donor and the typical transmission rate into recipients. This problem is under-determined given our data, so we made the additional assumption that the compound parameter $\Upsilon_{l,m}^{(t)} \sum_k \nu_{k,m}^{(t+1)} \gamma_{l,k,m}^{(t)}$ could be summarized by the effective, mouse-independent parameter $\Lambda_{l,\mathrm{eff}}^{(t)}$. With this assumption, the dynamics become

$$f_{l,m}^{(t+1)} = \left(1 - \tau_m^{(t)}\right)\Upsilon_{l,eff}^{(t)} f_{l,m}^{(t)} + \tau_m^{(t)} \bar{f}_l^{(t)}. \quad (18)$$

Eq. 18 jointly predicts $\Upsilon_{l,\mathrm{eff}}^{(t)}$, as well as lineage frequencies in recipient mice, $f_{l,r}^{(t)}$, from the dynamics in the donor(s), $f_{l,d}^{(t)}$, and the daily transmission measurements $\tau_r^{(t)}, \tau_d^{(t)}$ described above (Fig. 2C, 5D, S5, S14). The "transmission floor" and "complete engraftment" predictions are plotted for individual barcodes in Fig. 2E-J, 5E-J, using the initial conditions from day 14. In the cross-housing condition of Fig. 2, Eq. 18 is over-determined by two donor mice trajectories. In this case, we used the trajectory in the donor mouse with the larger measured frequency at day 14. In all cases, we replaced time-dependent transmission rates with averages $\bar{\tau}$ estimated from Fig. 2C, 5D. In Fig. 2E-J, we use $\bar{\tau}$ = 7% and 0.8% per day for P1- and P2-colonized mice, respectively. For clarity in Fig. 5E-J, we use $\bar{\tau}$ = ~1% per day, the average rate measured across community-colonized mice and time points, despite some variability between mice.

*The recipient:donor ratio*. We also summarized the engraftment behavior of barcodes via the ratio $\rho_l^{(t)} \equiv f_{l,r}^{(t)}/f_{l,d}^{(t)}$ (Fig. 2K, 5C), which is a convenient measure to compare the engraftment rates of barcodes at different donor (and recipient) frequencies. Without engraftment, $\rho_l^{(t)}$ is proportional to the daily transmission between mice. However, for the dozens of engrafting barcodes in Fig. 2 and thousands of engrafting barcodes between two of the three immediately co-housed, mono-colonized mice in Fig. 5B,C, the ratio rises above the daily transmission floor. In this scenario,

$$\rho_l^{(t+1)} = \frac{\left(1-\tau_r^{(t)}\right)\Upsilon_{l,\mathrm{eff}}^{(t)} f_{l,r}^{(t)} + \tau_r^{(t)} \bar{f}_l^{(t)}}{\left(1-\tau_d^{(t)}\right)\Upsilon_{l,\mathrm{eff}}^{(t)} f_{l,d}^{(t)} + \tau_d^{(t)} \bar{f}_l^{(t)}}, \tag{19}$$

where, for concreteness, we continue to assume that $\Upsilon_{l,\mathrm{eff}}^{(t)}$ is mouse independent. If a lineage's fitness per day is not very large, $\Upsilon_{l,eff}^{(t)} = 1 + s_l^{(t)}$, $\left|s_l^{(t)}\right| \ll 1$, and if $\tau^{(t)} \ll 1$, it is convenient to pass to continuous time, and keeping only terms $O(\tau, s)$,

$$\frac{(\rho_l^{(t+1)} - \rho_l^{(t)})}{\Delta \mathrm{t}} \simeq \frac{d\rho_l}{dt} = (\tau_r - \tau_d)\rho_l + \frac{\bar{f}_l}{f_{l,d}}(\tau_r - \tau_d \rho_l). \tag{20}$$

For illustrative purposes, we make several additional assumptions that apply to Fig. 5B,C. First, because the third co-housed mouse (colonized with barcode set 5) uptakes comparatively negligible populations of barcodes from the first two mice (Fig. 5B), we may approximate $\bar{f}_l \simeq \frac{1}{3}\left(f_{l,d} + f_{l,r} + 0\right)$. Then, for the two remaining mice, we take $\tau_r = \tau_d = \tau$. Then, with the initial value $\rho_l(t_0)$, Eq. 20 is solved as

$$\rho_l(t) = \frac{1 + \rho_l(\mathrm{t}_0) - (1 - \rho_l(\mathrm{t}_0))\exp[-\frac{2}{3}\tau(t - t_0)]}{1 + \rho_l(\mathrm{t}_0) + (1 - \rho_l(\mathrm{t}_0))\exp[-\frac{2}{3}\tau(t - t_0)]}. \tag{21}$$

Note that since Eq. 21 is recipient and donor frequency-independent, a related form applies to Fig. 2K if $\overline{f}_l \simeq \frac{1}{4}(2f_{l,d} + 2f_{l,r})$, corresponding to two donors and two recipients for each barcode. The least-squares fit of Eq. 21 to the data in Fig. 5C gives $\tau = 29\%$ per day, implying permissive engraftment conditions during the first days and weeks of mono-colonization of the mouse gut. On the other hand, we observed similarly high rates of transmission in the same mice beyond day 14 (upper limit of the distribution in Fig. 5D), showing that high rates of transmission (not necessarily with engraftment) are possible well after colonization.

**Measuring the spatial preference of adaptive lineages**

For most mice, cecum and feces were sampled on the day of sacrifice, $T_f$, providing the opportunity to study emergent spatial structuring among lineages. Fig. S7 reports cecum frequencies $\hat{v}_{l,m}^{(T_f)}$, calculated according to Eq. 1, as well as the "cecum bias" of a barcode lineage *l* in mouse *m*, defined as

$$\beta_{l,m} = log \frac{\hat{v}_{l,m}^{(T_f)}}{\hat{f}_{l,,m}^{(T_f)}}. \tag{22}$$

**Whole genome sequencing and analyses**

*Isolate sequencing*. Fecal samples, collectively representing all 10 SD mice and 2 MD mice from our first experiment, were collected on days 9, 17, 36, and 51 and grown on plates. Four hundred eighty-nine colonies, roughly evenly distributed across mice, were selected for whole-genome sequencing. To identify mutations, we used *breseq*[14] v. 0.35.7 in consensus mode with default parameters on the paired-end reads (Methods). Ten colonies were sequenced at low coverage or were not derived from a single barcode, and hence were excluded from further analyses.

The remaining 479 clonal isolates were sequenced with median coverage of 212X (range 24X-553X). These isolates represented 161 barcodes, 60 of which reached >1% frequency in at least one mouse and hence were almost certainly adaptive. A large fraction of the other 101 barcodes were also likely to have acquired adaptive mutations, despite having only been sampled once: these barcodes represented 40% of the 188 isolates sampled at day 17, whereas our analysis of the median fold change over time suggests that the non-adaptive population comprised <20% of cells at that time (Fig. 1E).

*Metagenomic sequencing.* We performed longitudinal metagenomic sequencing of all mouse and *in vitro* samples, as well as all input libraries. To analyze these data, we used *breseq* v. 0.35.7 in polymorphism mode. We also used sequencing of the input libraries to analyze isolate genomes, as described below.

*Identifying driver mutations in isolates*. For each sequencing library, *breseq* generates a large list of short read-based evidence, which is evaluated to call a high-confidence list of variants. We used the evidence and mutation lists to identify putatively adaptive structural variants and simple point-like mutations, respectively.

*Identifying simple point-like variants*. We first collated 129 point-like variants ("SNP", "SUB", "DEL", and "INS") identified by *breseq* in >90% of isolate genomes. These mutations represent pre-existing mutations fixed in the ancestral genotype (which differed from the reference strain input to *breseq*). We augmented this list with 57 variants observed in <90% of isolates but concentrated at five loci – in *BT1042*, and in the intergenic regions of (*BT1040*/*BT1041*), (*BT3239*, *BT3240*), (*BT4054*, *BT4055*), and (*BT2672*, *BT2673*) – and 152 polymorphisms measured at >10% in at least two day 0 metagenomic libraries. These additions may represent hypermutable regions or gene copy number variation in the ancestral genotype.

After excluding pre-existing variants or variants in hypermutable regions, we were left with a list of 70 unique variants, all but two of which were represented in single barcodes). 128 barcodes harbored no point-like variants in a majority of their isolates, suggesting other sources of genetic (or otherwise heritable) variation that drove lineage dynamics.

*Identifying structural variants and phase variation*. We utilized "junction" (JC) evidence from *breseq* to identify structural variants. JC-type evidence represents reads with different portions aligning to two distant regions of a genome. Compared to simpler point-like mutations or small-scale structural variants (spanning less than the read length), more complicated structural variants—in particular, large-scale amplifications and genomic inversions—must in general be distinguished from mapping artifacts[14,15].

To this end, we first collated a list of JC evidence across all isolates. For each isolate, we included only JC evidence spanning genomic distances >10 bases; measured at >10% frequency; and supported by >5 reads in the isolate library, while ignoring all JCs with >10% frequency in at least two day 0 metagenomic libraries, or involving the same five loci mentioned above. (The lenient frequency threshold avoids ascertainment biases that would influence one step in our validation strategy, described below.)

Most types of structural variation should register two or more JCs in a sequenced genome[14]. We thus sought to pair and classify JC evidence as: inversion-mediated phase variation (“inversion”), “insertion” (e.g., of a mobile genetic element), or ambiguous structural variation (“SV”). To do so, we first identified possible invertible repeat-mediated phase-variable loci within the *Bt* genome with the *einverted*[16] tool in the EMBOSS software package, using the following search parameters: max repeat=10000; gap penalty=100; minimum score threshold=50; match score=5; mismatch score=-10.

We then paired and classified JCs in the following manner. We denote a pair of JCs as $(j_1, j_2)$, spanning genomic sites $(x_1, y_1)$ and $(x_2, y_2)$ and detected in $n_1$ and $n_2$ isolates, respectively. If $j_1$ and $j_2$ were jointly present in fewer than min $(n_1, n_2,)/2$ of isolates, or if $\max(|x_1 - x_2|, |y_1 - y_2|) > 50$, we did not consider pairing them further. Otherwise, we labelled $(j_1, j_2)$ a putative “inversion” if they were in a region spanned by invertible repeats detected by *einverted* (with $\max(|x_1 - x_2|, |y_1 - y_2|) < 50$ reflecting the maximum size of the invertible repeat), and showed the expected strand orientations for such inversions[14]. We instead labelled JC pairs as evidence of “insertion” if the paired junctions shared one break point within 50 bases and the other end points were >10 kb away. All other pairs were labelled “SV.” Finally, we labelled 30 single JCs that did not satisfy our pairing criteria with any other JC as an “unpaired JC.” Deletions could generate unpaired JCs, but in no cases did we find a corresponding absence of read coverage. On the other hand, among the 75 JC pairs, some single JCs were represented in multiple pairs, reflecting the larger fact that some structural variants involve multiple junction events. We did not seek to further merge JC pairs and/or unpaired JCs, and instead relied on manual inspection and validation to confirm more complex structural variants.

Unlike point-like mutations, many JCs were found in many barcodes. To distinguish structural variants from mapping artifacts, we tested JCs for uneven distribution across barcodes (and/or time points for barcodes sampled at multiple times), since JCs arising mapping artifacts are unlikely to be correlated with barcode identity or sampling time point. For each JC detected in $k > 1$ sampled isolates, we constructed the 193×2 contingency table denoting the number of clones with or without that JC for each (barcode, time point) combination. To test for significant non-independence in the table, we first repeatedly re-sampled the table under a multivariate hypergeometric distribution maintaining the same row and column totals. We then calculated the $p$-value of the observed tables as the number of re-samples with a probability less than or equal to the probability of the observed table. This strategy provides a Monte-Carlo approximation of Fisher's exact test for arbitrary sized tables. After calculating $p$-values for all paired and unpaired JCs observed in >1 isolate, we calculated their respective $q$-values via the Benjamini-Hochberg method to control the false detection rate.

Some inversions were found at intermediate frequencies <0.5 in most clones in which they were detected (note the necessity of initially including low frequency JC evidence to observe this pattern). These instances plausibly represented rapidly mutating and positively selected inversions arising during colony growth, rather than mutations arising *in vivo*. Nonetheless, many of these were associated with a smaller number of barcodes than expected if randomly distributed across colonies, suggesting biologically interesting epistatic interactions between inversions arising during colony growth and other, *in vivo*-arising mutations present in some barcode lineages. We additionally note that all inversions were also detected in metagenomics (see below).

Table S1 reports the putative driver mutations in each barcode, including simple variants as well as JC evidence. For each barcode, a variant was included as a putative driver if it was measured in a majority of associated isolates and, for JC evidence, if it was measured at a median frequency >0.5 across detected isolates, and either had a $q < 0.05$ or was only detected in a single isolate (such that significance could not be ascertained). Table S2 additionally reports all paired and unpaired JCs, with the number of distinct barcodes and isolates in which they were found, their $q$-values, and their median frequencies in detected isolates. Note that some JC evidence is represented in multiple merged pairs, because some complex structural variants produced more than two junction events. We did not de-replicate these cases in the tables.

*Comparing isolate mutations to metagenomic data during colonization with a diverse community and* in vitro. We next sought to compare the mutations identified from isolate sequencing to metagenomic sequencing in both community-colonized mice (Fig. 4,5) and *in vitro*-evolved populations (Fig. 6). A full analysis of mutations was especially difficult in community-colonized samples, in which coverage was typically around 10 reads. Nonetheless, we were able to address our primary goal, which was to determine if metagenomic data from community-colonized mice or *in vitro*-evolved populations showed evidence of the same mutations that repeatedly arose in isolates sampled from mono-colonized mice.

We used a previously developed *breseq*-based pipeline[5] that distinguishes likely mutations from sequencing artifacts by leveraging the correlated presence of genuine variants across longitudinal metagenomic samples. In the case of mice co-colonized with the community, we did not competitively map reads to other community members (including the other *Bacteroides* species), which permitted more true read mappings to *Bt* at the expense of a (far larger) number of false read mappings. This tradeoff was warranted given our specific interest in determining the presence/absence of mutations detected in the isolates, since we subsequently validated these mutations via manual inspection of mapped reads.

We confirmed that our pipeline could identify isolate-validated mutations in metagenomic sequencing of the mono-colonized mice in Fig. 4, 5. In particular, the same point mutation in *BT0867* and the CPS 8/PUL10 amplification were observed in 3 of 5 and 4 of 5 mono-colonized mice, respectively (Fig. S9, S10), along with junctions representing transposition of mobile elements in 3 of 5 mono-colonized mice (Fig. S9).

On the other hand, neither the *BT0867* or CPS8/PUL10 mutations that were repeatedly detected in mono-colonized mice, nor any other non-inversions reported in Table S2, were detected in any of the co-colonization mouse samples (Fig. S9, S10). Similar analysis of the *in vitro* populations found limited evidence of overlap in non-inversion mutations with mono-colonized mouse samples. Table S6 reports putative driver mutations in individual wells. Each well was sampled after 1, 15, and 31 passages. To limit false positive calls, we only report mutations detected at both passages 15 and 31 in at least one well (and that were not among the pre-existing mutations excluded above). The general lack of the same non-inversion structural variation confirms our inferences from barcode dynamics of environment- and community-specificity of the adaptive landscape of *Bt* (Fig. 4E-G).

### Supplementary Figures

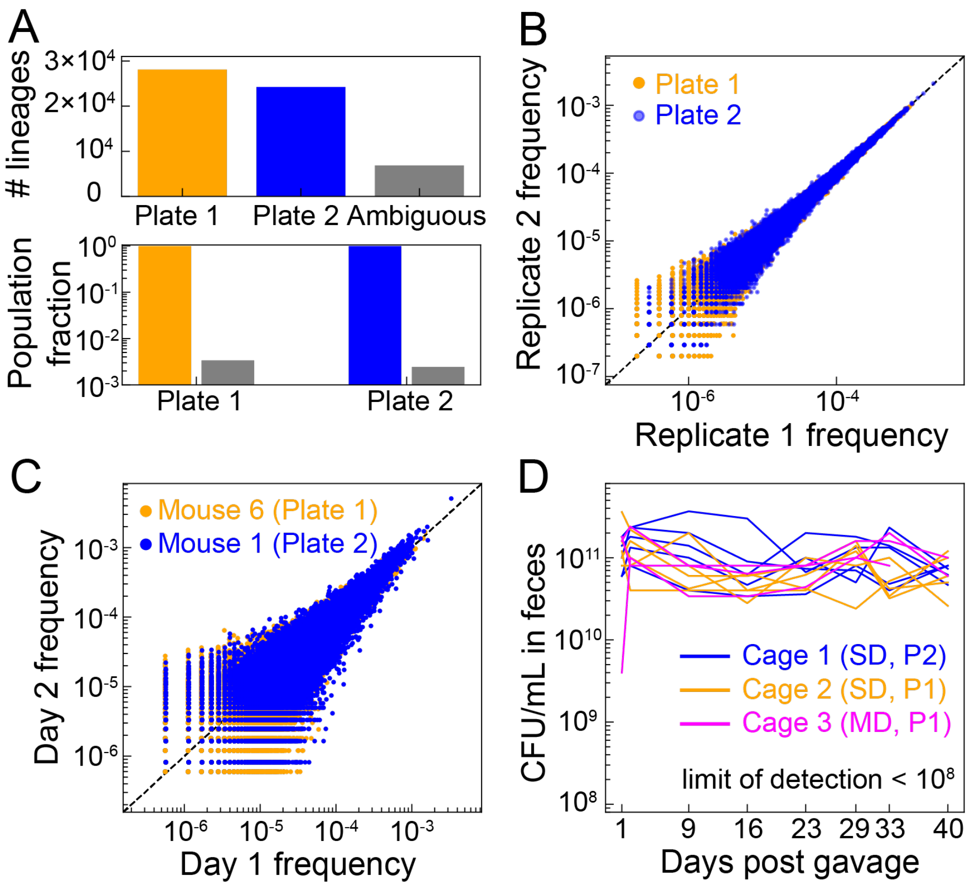


**Figure S1: Size and plate assignments of *Bt* barcode library, which stably engrafts in mice.**

A) Top: 28,093 and 24,235 barcodes were unambiguously assigned to either plate 1 (P1) or plate 2 (P2), respectively (Supplementary Text). Bottom: the ~6,856 ambiguous barcodes comprise a negligible fraction of the plate 1 and plate 2 input libraries.

B) Technical replicates of PCR amplification and sequencing of the input libraries were highly correlated.

C) P1 and P2 libraries in two representative mice, sampled at days 1 and 2, were correlated over a broad range of frequencies.

D) Barcoded *Bt* consistently and stably colonized mice within ~1-2 days, as measured by CFUs per mL of feces.

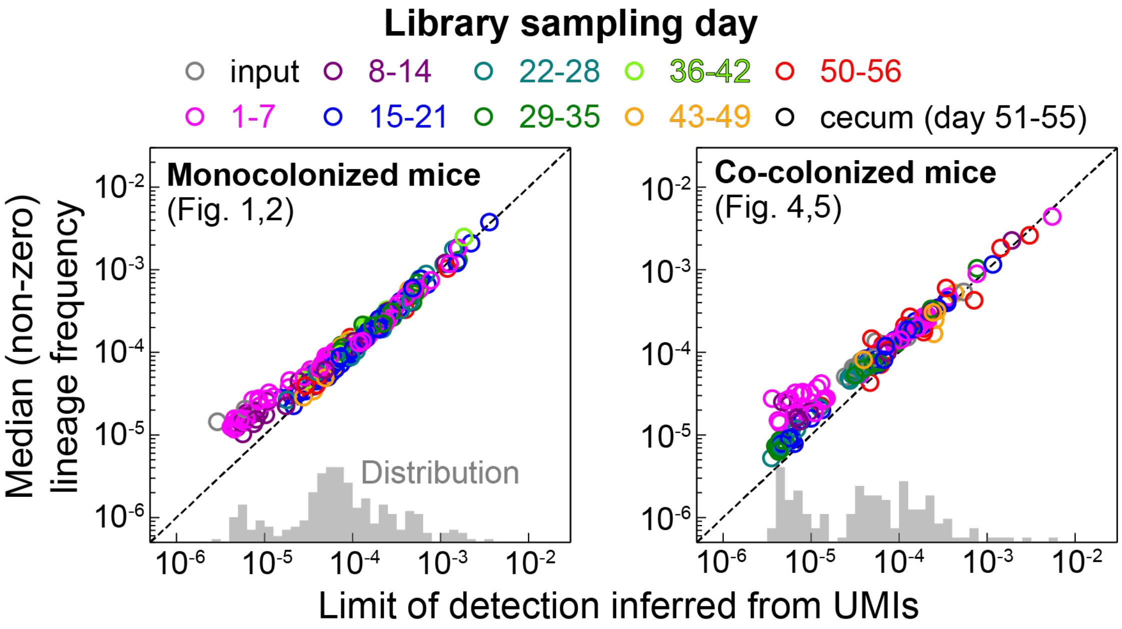


**Figure S2: Pseudo UMI-dereplication robustly estimates the effective sequencing depth in barcode libraries.** The median frequency of detected lineages *in vivo* was generally similar to the limit of detection inferred by independent means (Supplementary Text). Nonetheless, most libraries still probed >2 orders of magnitude smaller frequencies than those typically accessible by whole-genome sequencing methods (~$10^{-2}$).

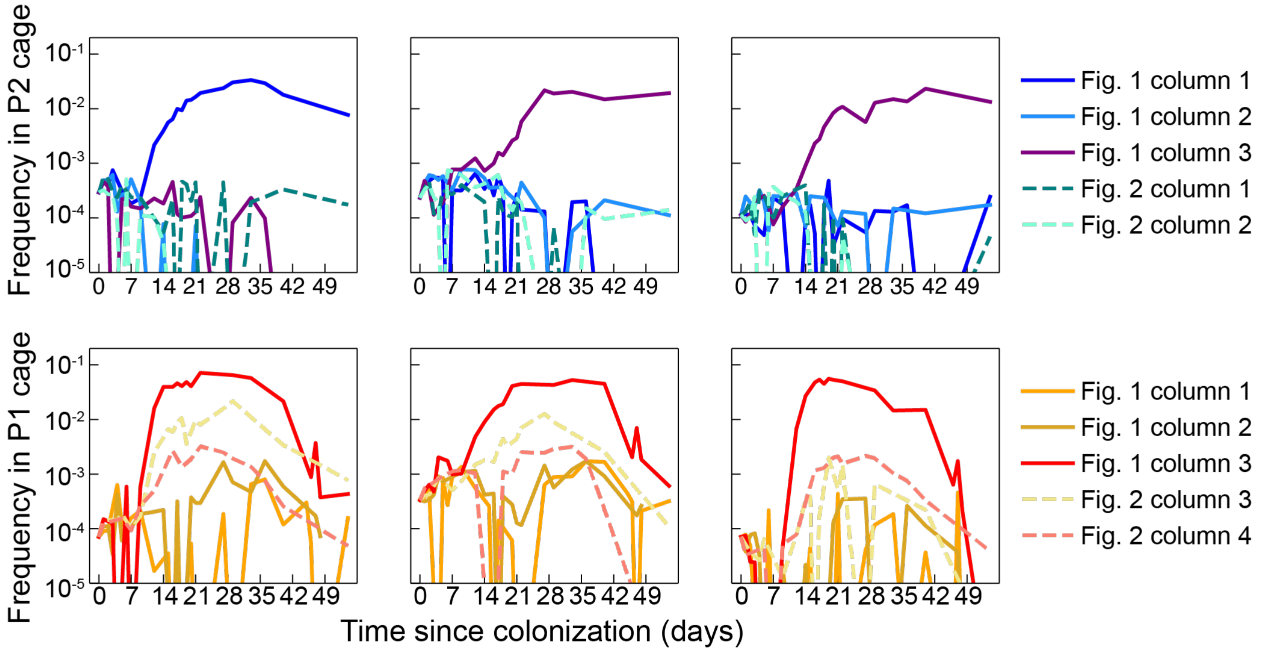


**Figure S3: Examples of barcodes that expanded in one mouse to frequencies >1% without expanding in other mice.** This divergence across mice could be driven by local adaptation to a host, an evolved inability to transmit across hosts, or by neutral processes (e.g., random colonization of spatial niches). Dashed curves represent mice that were cross-housed on day 14 (Fig. 2).

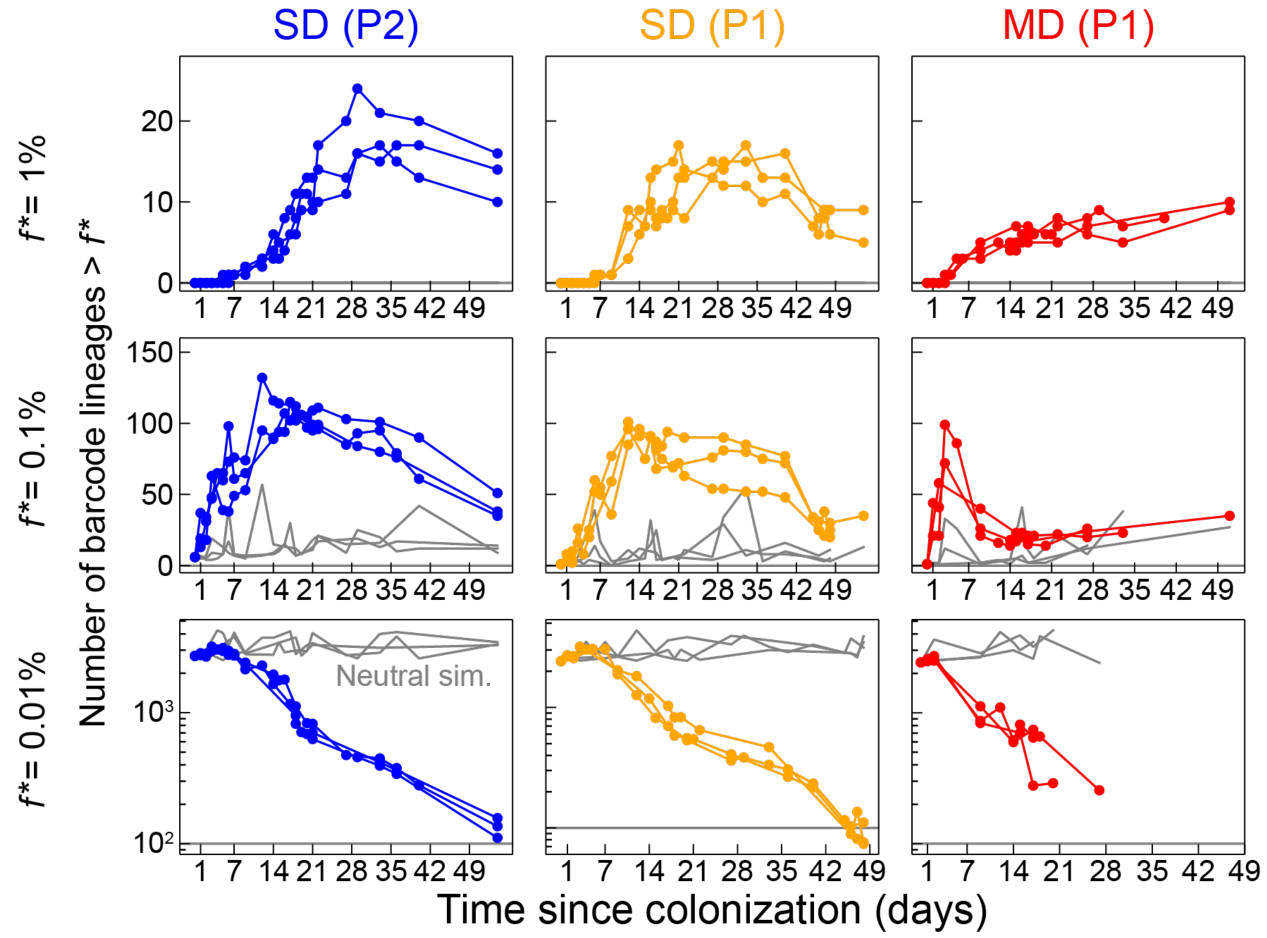


**Figure S4: Number of barcode lineages in mono-colonized mice measured above various frequency thresholds.** Each curve represents a single co-housed mouse in a cage. For clarity due to the sudden change in housing status, cross-housed mice were excluded. Libraries with effective limits of detection (Fig. S2) above the frequency threshold are excluded. The neutral simulation (sim., gray) methodology is described in the Supplementary Text.

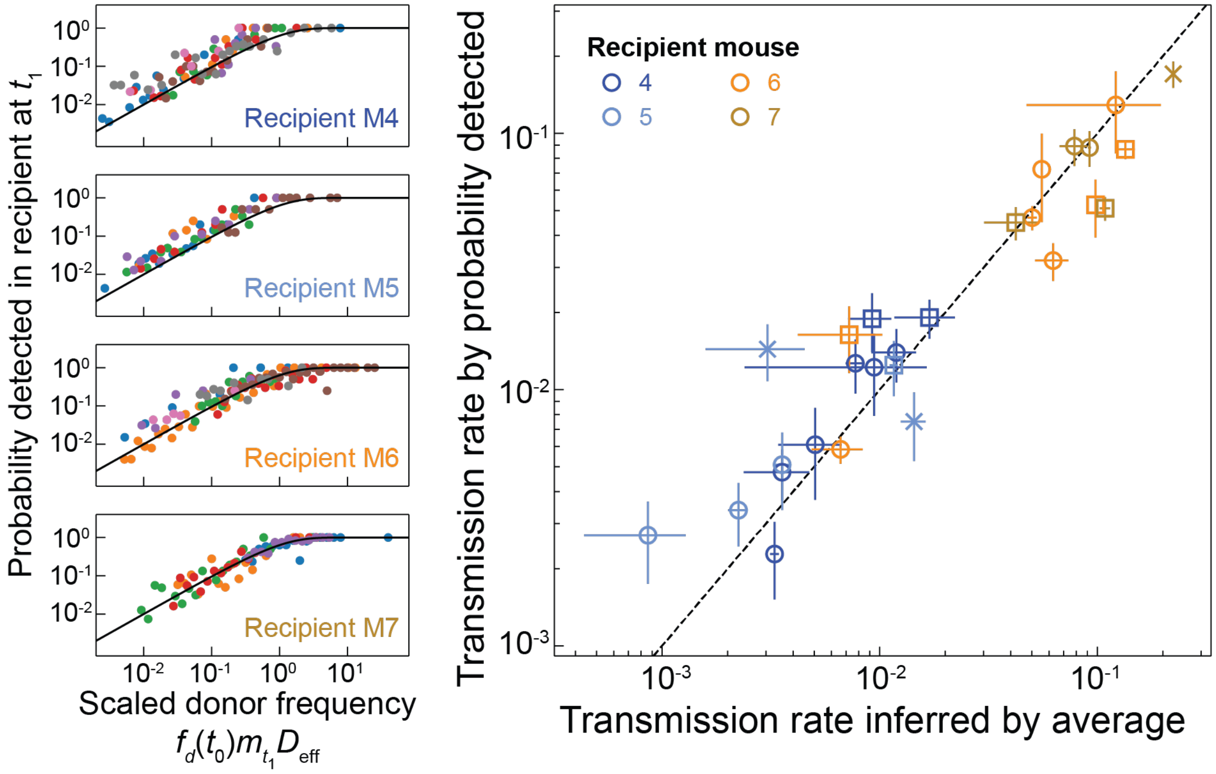


**Figure S5: Consistency of uniform daily transmission measurements in cross-housing experiment.** Left: across recipient mice (panels) and time intervals (colors), the probability of barcode detection in the recipient as a function of the donor frequency, rescaled by the inferred transmission rate and effective sequencing depth, was consistent with a Poisson distribution (black curve). Right: transmission rate inferred by the probability detected, as in left panels, versus an alternative estimate from a weighted average of recipient and donor frequencies (Supplementary Text). Error bars represent ± 1 SEM; the uncertainty across estimates is within ~3-fold, which is small compared to the variability across time and mice.

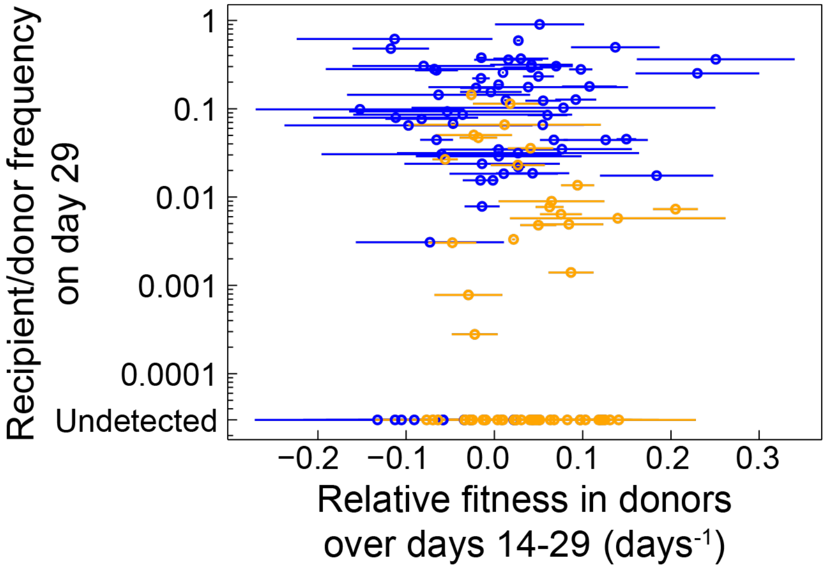


**Figure S6: Lineage growth rate in donors is not predictive of engraftment in recipients.** The relative fitness (Supplementary text) of high-frequency barcodes was estimated in each of two donor mice over the first two weeks of cross-housing with recipients. Circles and bars represent the mean and range of relative fitnesses estimated in the two donor mice, and *y*-values are the ratio of mean frequencies in the two recipients to the two donors. Orange and blue represent barcodes originating in P1- or P2-inoculated mice, respectively.

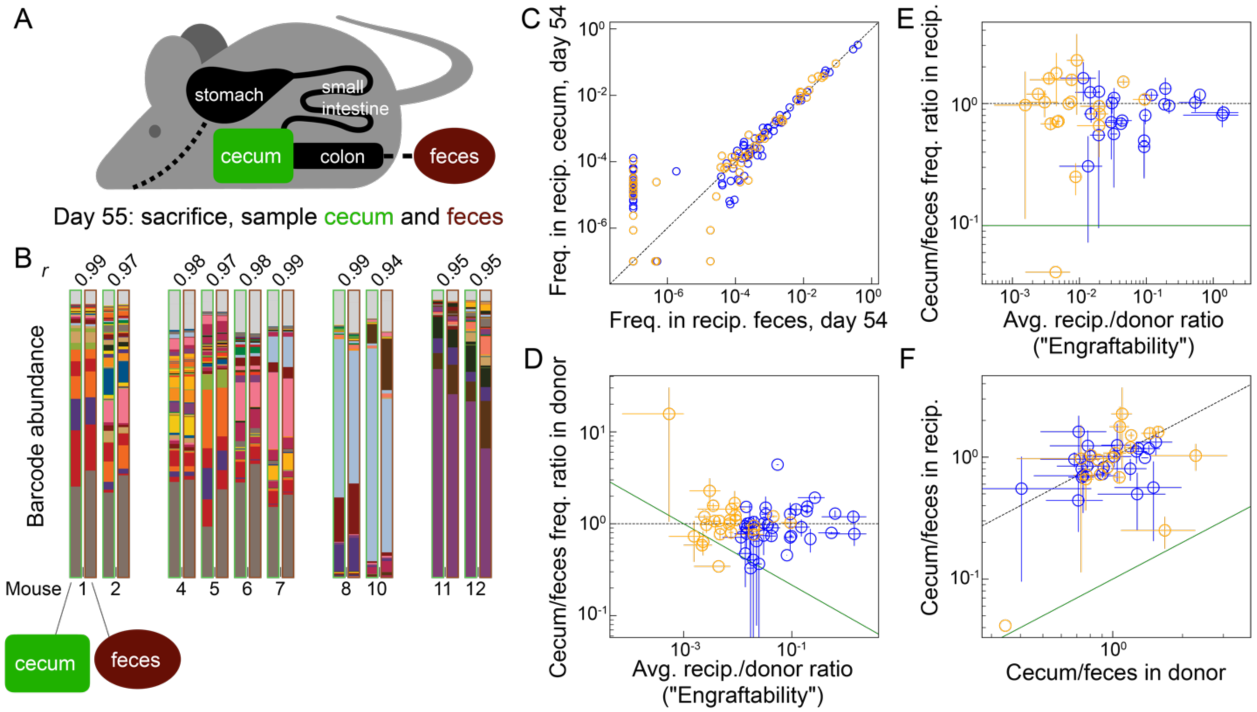


**Figure S7: Spatially resolved sampling demonstrates correlation between the cecal and fecal abundances of *Bt* barcodes.**

A) Schematic of the mouse gastrointestinal tract. On the final day of fecal sampling, mice were sacrificed and cecal contents were collected and homogenized for sequencing.

B) Barcode relative abundance was highly correlated between the cecum and feces of individual mice. Top: Pearson's correlation coefficient (*r*).

C) Cecum versus feces measurements of transmitted barcodes in recipients from Fig. 2K (orange and blue represent P1 and P2 barcodes, respectively). Each barcode is plotted twice, once for each of two recipients.

D) Ratio of cecum:feces barcode frequency in donor versus engraftability (Fig. 2K). Error bars along the $x$- and $y$-axes indicate the ranges of engraftability and cecum:feces ratios estimated across two recipients and two donors, respectively (when available). Green line is the qualitative expectation if barcodes adapted to the cecum have a more difficult time engrafting in already colonized hosts.

E) Ratio of cecum:feces frequency in recipient versus engraftability. Error bars defined as in (D). Green line is the qualitative expectation if engraftment is reduced in the cecum versus the spatial compartment representing the feces.

F) Cecum:feces ratios in the recipient versus donor. Green line combines the qualitative expectations from (D) and (E).

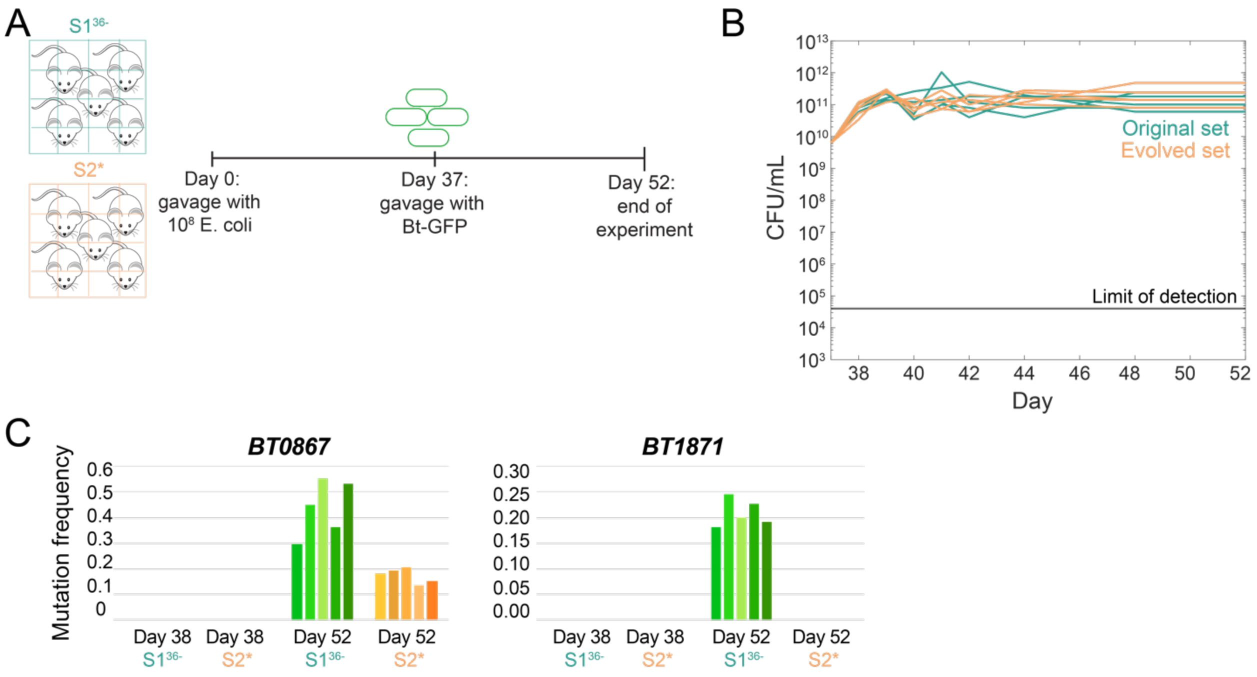


**Figure S8:** ***Bt*** **stably colonizes mice previously colonized with** ***E. coli*****.**

A) Two cages of five mice each co-colonized with (barcoded) *E. coli* from our previous study[13]; one cage was inoculated with an *E. coli* strain that had been isolated from a population evolved in ex-germ-free mice for ~2 months (S2$^{*}$), and the other cage received a distinct set of mouse-naïve lineages (S1$^{36-}$). On day 37, mice were gavaged with GFP-tagged *Bt* (distinct from our barcoded library).

B) *Bt* stably colonized mice in both cages to similar levels, regardless of the evolutionary history of the resident *E. coli*.

C) Metagenomic sequencing revealed two prominent mutations in *Bt* (in *BT0867*, discussed in the main text, and in *BT1871*) that emerged within two weeks of gavage.

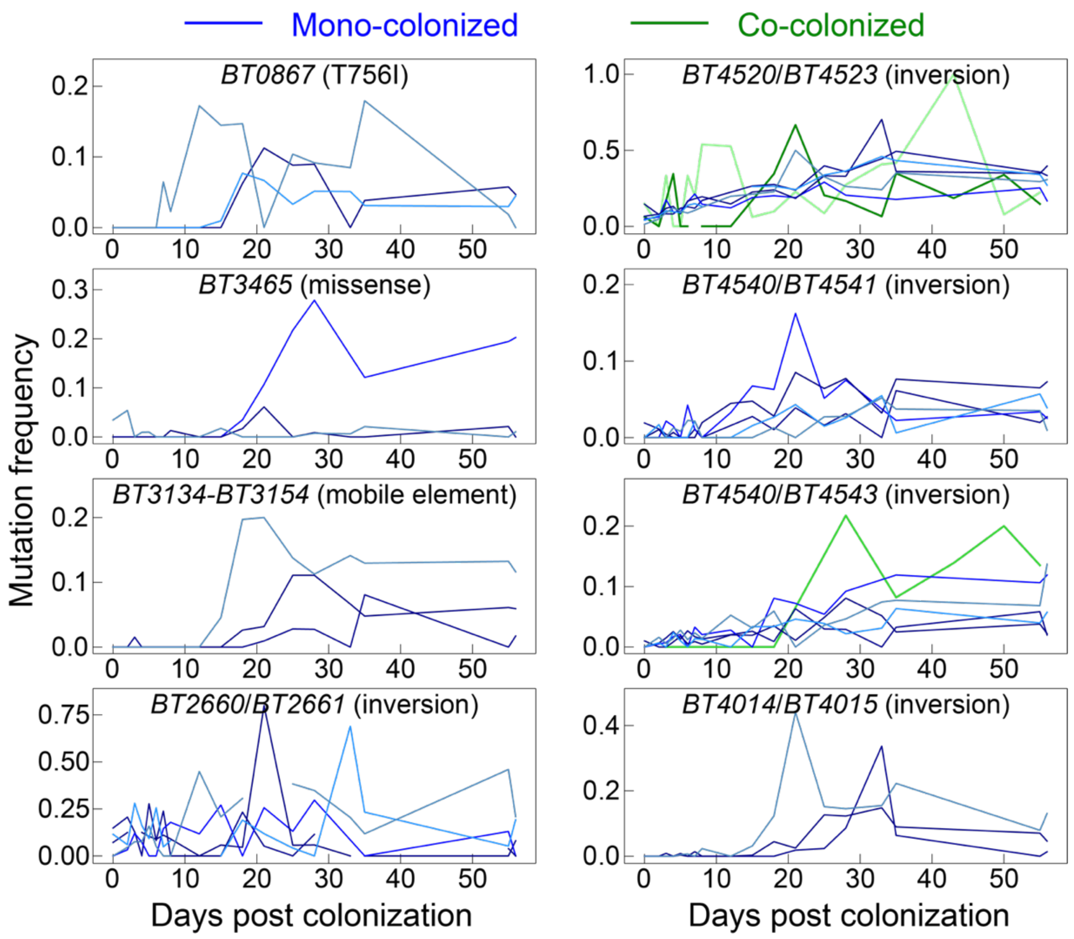


**Figure S9: Longitudinal metagenomic sequencing reveals repeated but community-dependent selection of similar mutations across evolution experiments.** Each panel represents a locus with putative driver mutations detected in isolates from the first cohort of mice (Fig. 1,2), with mutation frequency trajectories plotted in individual mice from the experiments in Fig. 4,5 (shades of blue and green represent mono-colonized and co-colonized mice, respectively). Among mutations at commonly mutated loci in mono-colonized mice, only those representing high-rate inversions in restriction-modification complexes (*BT4520/BT4523, BT4540/BT4543)* were observed in community-colonized mice.

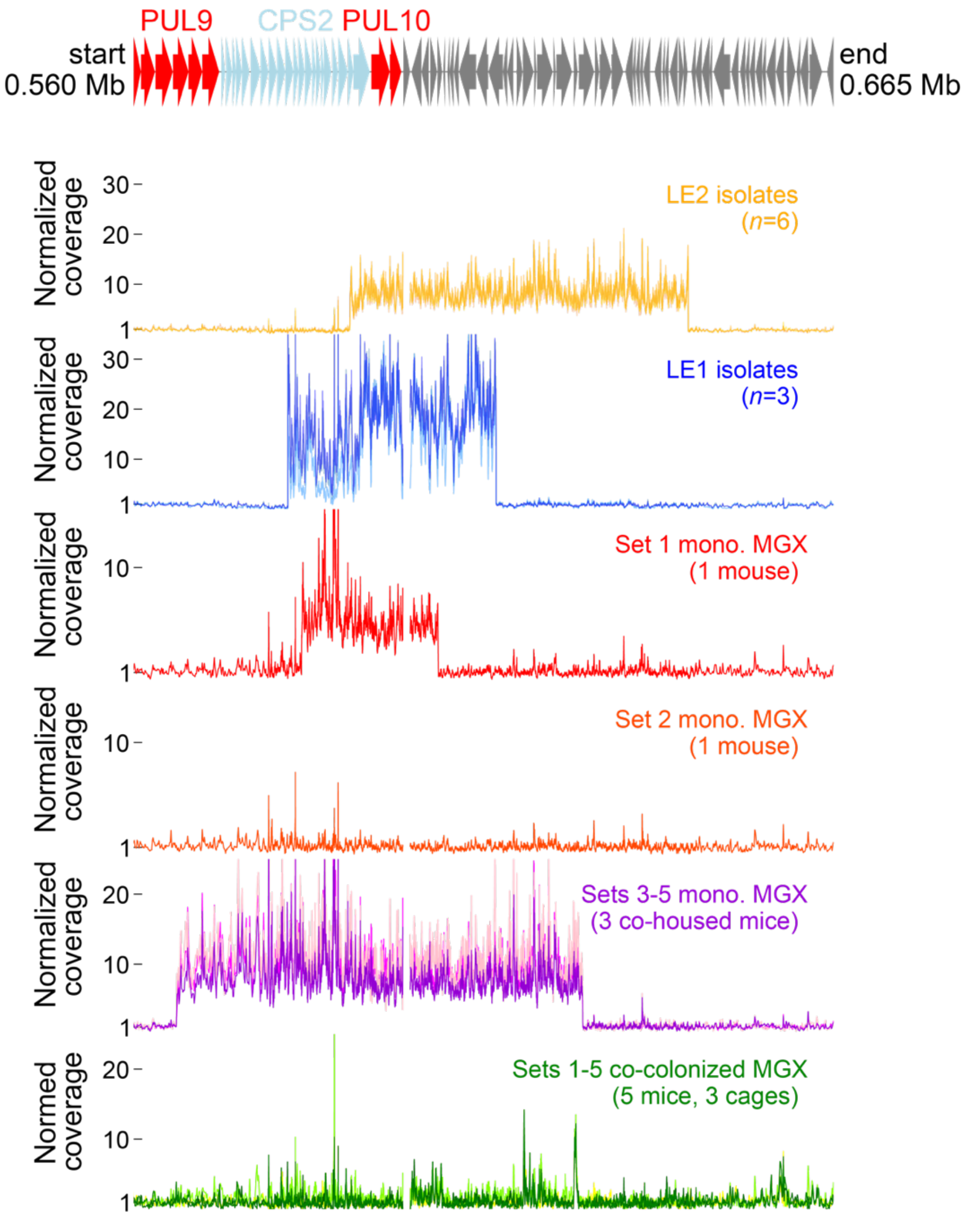


**Figure S10: Analog of Fig. 3C for metagenomic samples of mono-colonized and co-colonized mice.** Top: the 105-kb region of *Bt* containing CPS2 and PUL10. Bottom six panels: local coverage of that region in various samples, normalized by the local coverage in a metagenome of the ancestor. The top two coverage tracts reproduce the LE1 and LE2 barcode isolates shown in Fig. 3C, for reference. The next three tracts are from metagenomic stool sequencing (~100X coverage) of five mono-colonized mice (Fig. 4,5) on day 55, split by cage. The bottom tract shows the absence of evidence of amplification from local coverage in day 55 metagenomes (~10X coverage) in all five community-colonized mice (Fig. 4,5).

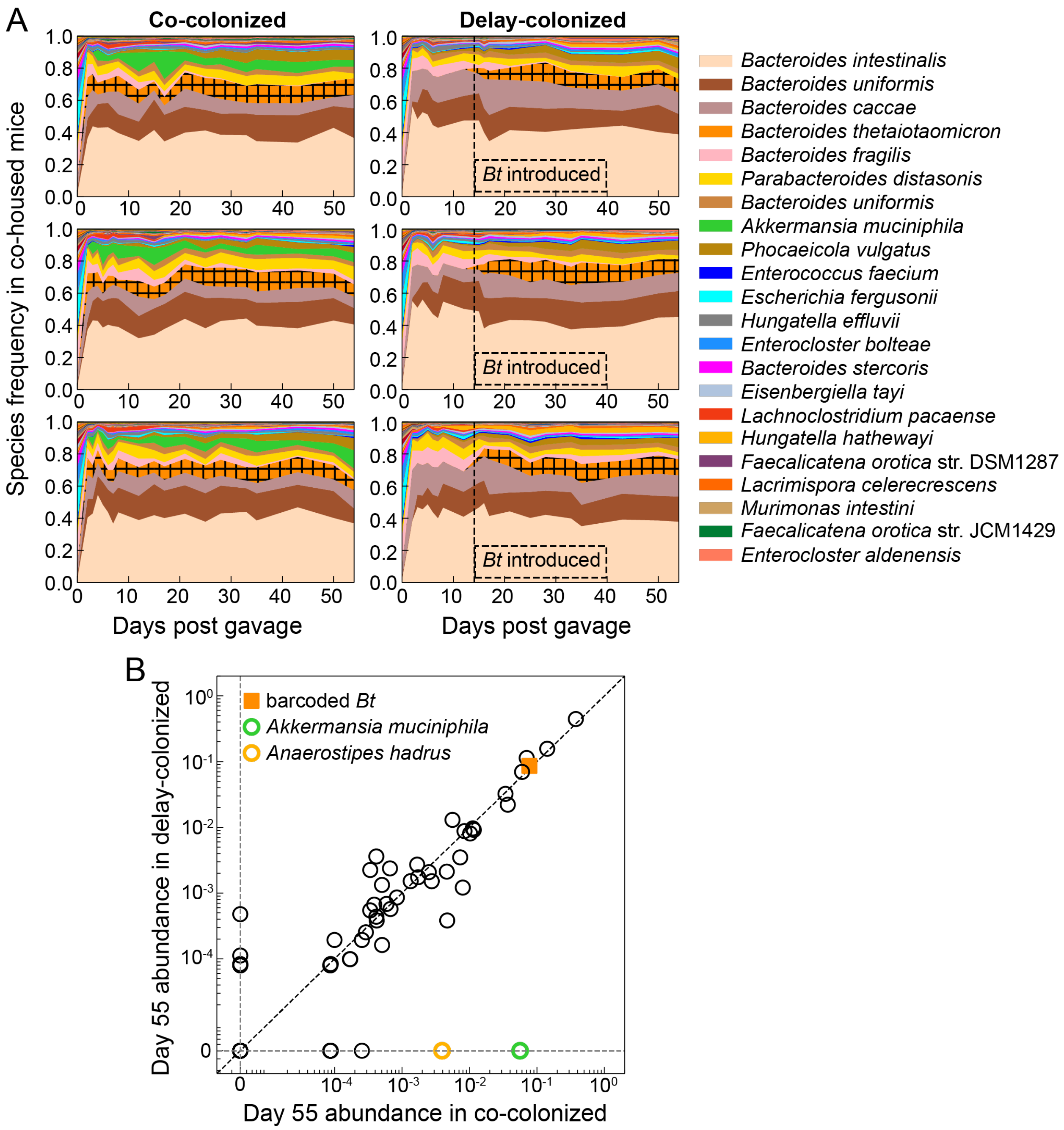


**Figure S11: Stable colonization of barcoded *Bt* with a diverse community.**

A) Analog of Fig. 4B for the three co-housed, co-colonized and three co-housed, delay-colonized mice (Fig. 5B).

B) Median ASV relative abundances in five co-colonized versus five delay-colonized mice (limit of detection ~$10^{-3.5}$). Most abundances at the end of the experiment were similar across colonization conditions. Two exceptions are *Akkermansia muciniphila* and *Anaerostipes hadrus*, which were measured above 4% and 0.2%, respectively, in co-colonized mice, but were undetected when *Bt* colonization was delayed.

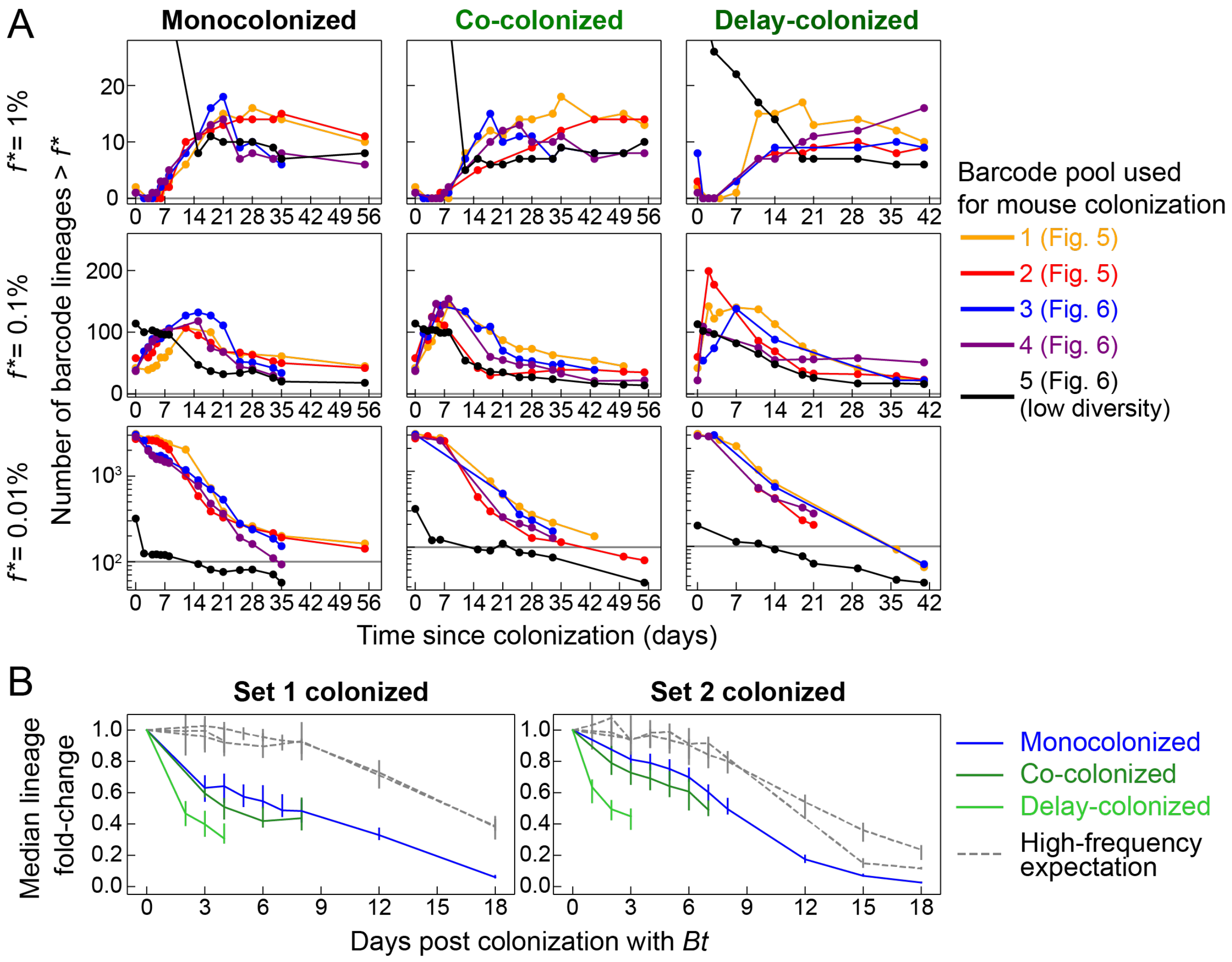


**Figure S12: Number of barcodes detected and median lineage fold-change for the second cohort of mice (Fig. 4,5).**

A) Analogous to Fig. S4, each curve in each panel represents a single mouse. Each column corresponds to a community colonization condition. Mice colonized with barcode pools 1 and 2 were singly housed, and mice colonized with pools 3, 4 and 5 were immediately co-housed. Sequencing libraries with effective limits of detection (Fig. S2) above the frequency threshold are excluded.

B) Median fold-change (MFC) over time in singly housed mice under all three community colonization conditions (Supplementary Text), analogous to Fig. 2E. Between 1900-4400 barcodes, constituting the upper half of input frequencies (and 68-82% of cells in the inoculum) were used to calculate each curve. The strength of genetic drift used to simulate the high-frequency expectation was increased ten-fold ($N\tau = 10^5$ days) relative to Fig. 1E, to account for the possibility of a reduced *Bt* effective population size in the presence of a diverse community.

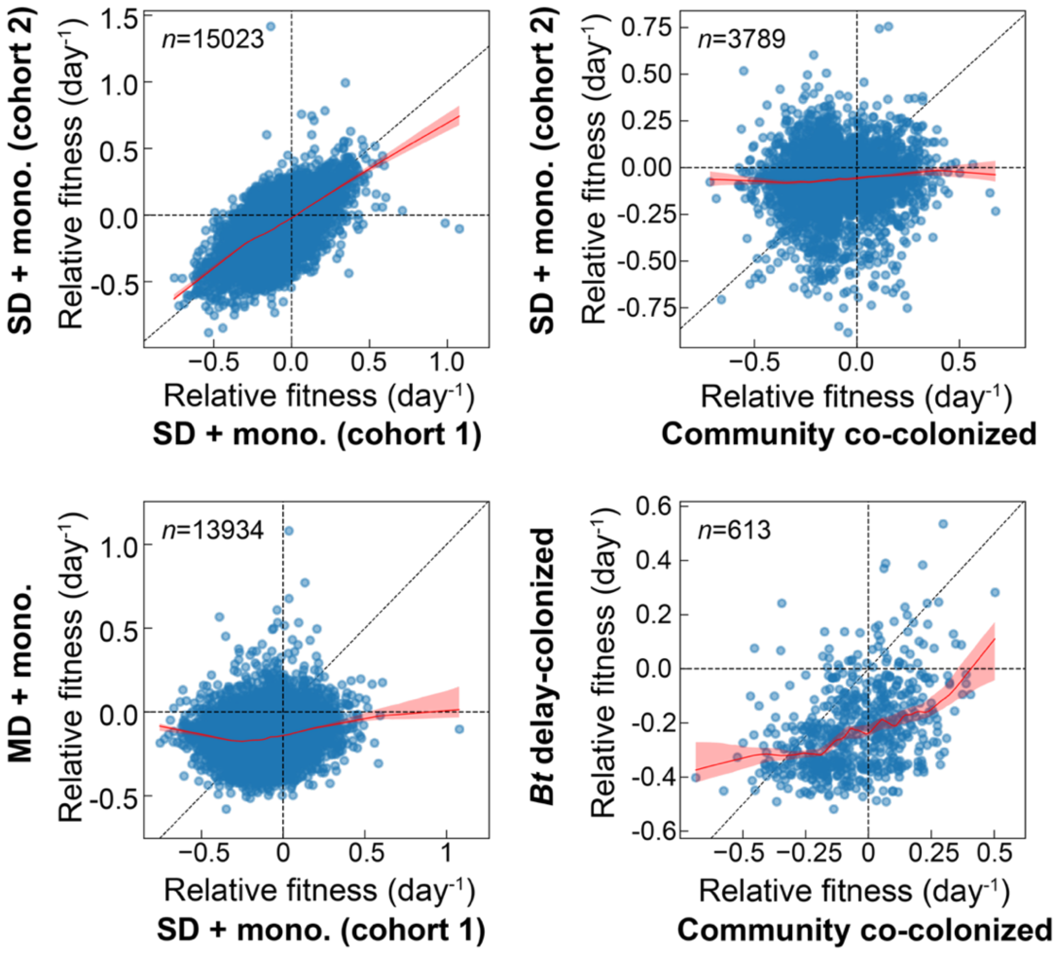


**Figure S13: Joint distribution of fitness effects across *in vivo* environments**. Fitnesses were measured over 7±2 days, beginning 2 to 3 days after initial *Bt* colonization (Supplementary Text). For each pair of environments (panel), barcodes measured at greater than 2 effective reads (Supplementary Text) at the first time point in each environment were included. The number of barcodes (*n*) included in each panel are indicated in the top left. Red curves show LOWESS regressions using 30% of the data per local estimate, and red shaded regions are 90% confidence intervals obtained by repeated LOWESS on 100 bootstraps of the data. Time intervals: SD and MD + mono-colonized (mono., cohort 1), days 2–9; SD + mono-colonized (cohort 2), days 3-8;

community co-colonized = days 3–8 (or 7 for barcode set 2-colonized mouse); delay-colonized=2–11 days after *Bt* introduced (16–25 days after community colonization).

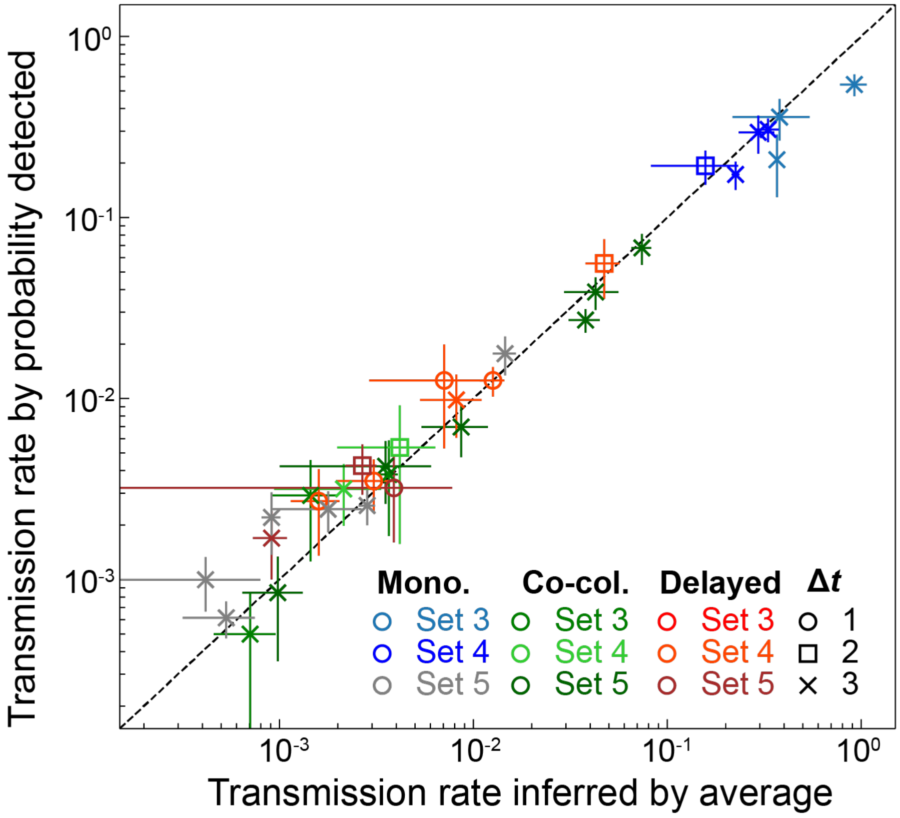


**Figure S14: Consistency of daily transmission measurements in community-colonized mice.** Transmission rates estimated by probability of detection show good agreement with a separate estimate from averaging lineages (Supplementary Text), analogous to Fig. S5.

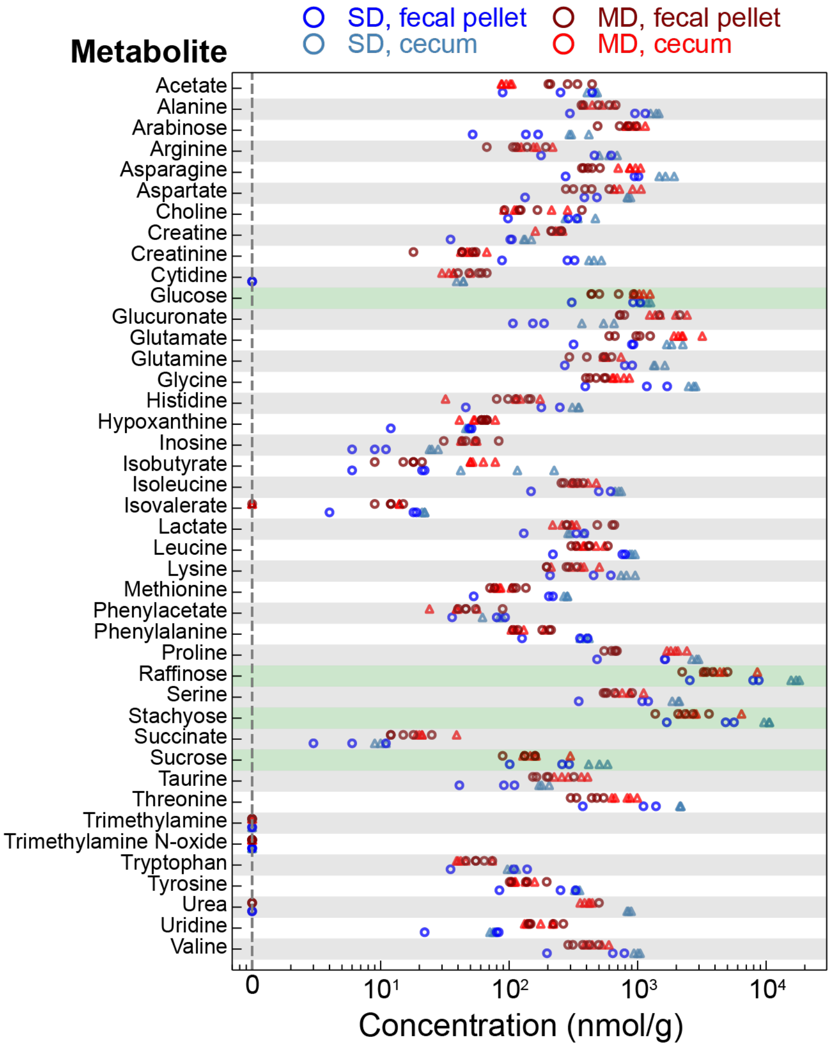


**Figure S15: Metabolomics of germ-free mice.** Three and five mice fed a standard or MAC-deficient diet, respectively, were sacrificed, and cecal contents and fecal pellets were harvested for metabolomic profiling. Metabolites highlighted in green (glucose, raffinose, stachyose, and sucrose) were used as carbon sources for *in vitro* passaging of barcoded *Bt* (Fig. 6).

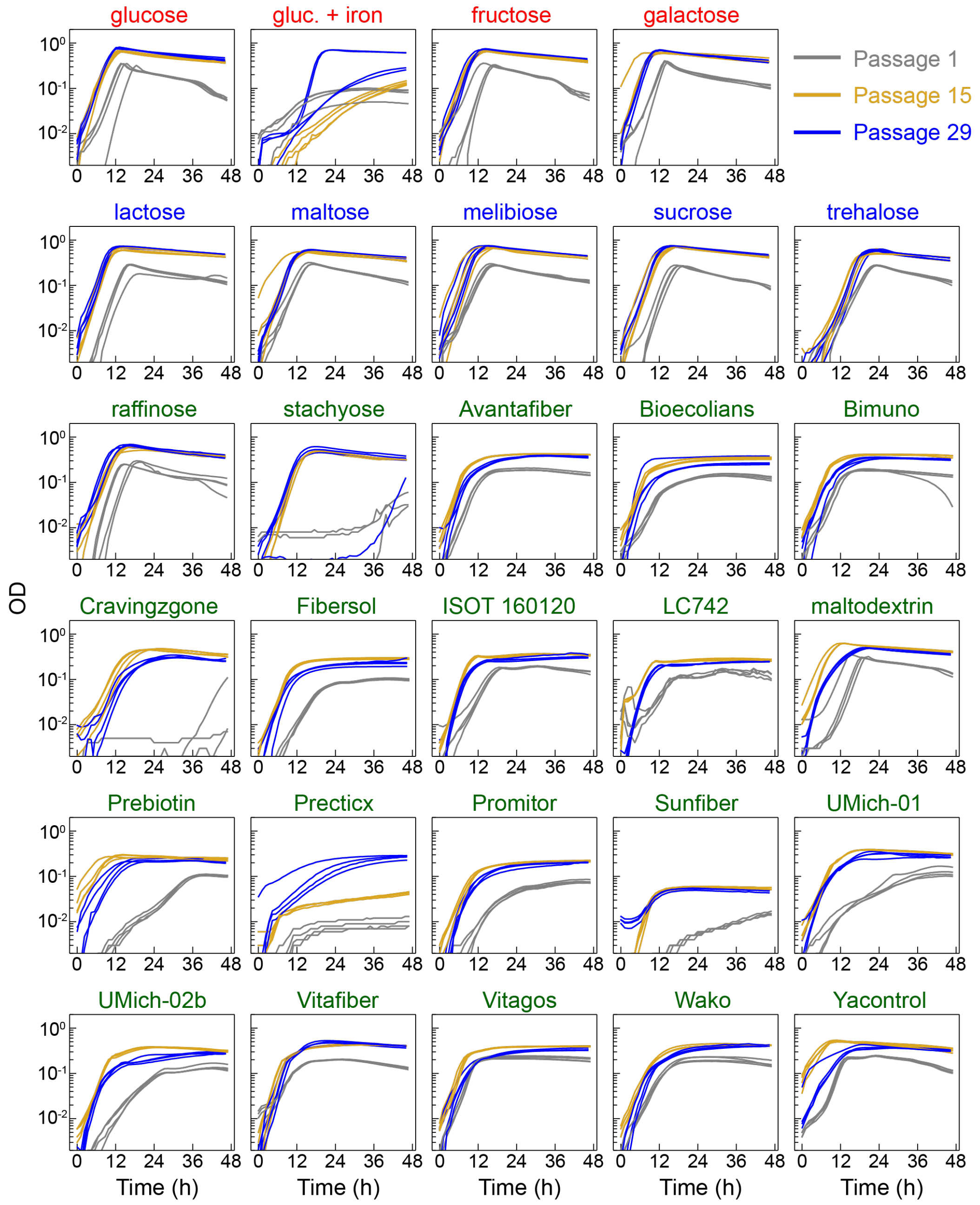


**Figure S16: Growth curves of *Bt* populations during evolution in various carbon sources.** Growth of four replicate populations in each carbon source was monitored at three time points (1, 5, and 9 weeks, corresponding to passages 1, 15, and 31). Plotted is optical density (OD) with readings from blank wells in the same plate subtracted. Red, blue, and green carbon sources are mono-, di-, and poly-saccharides, respectively.

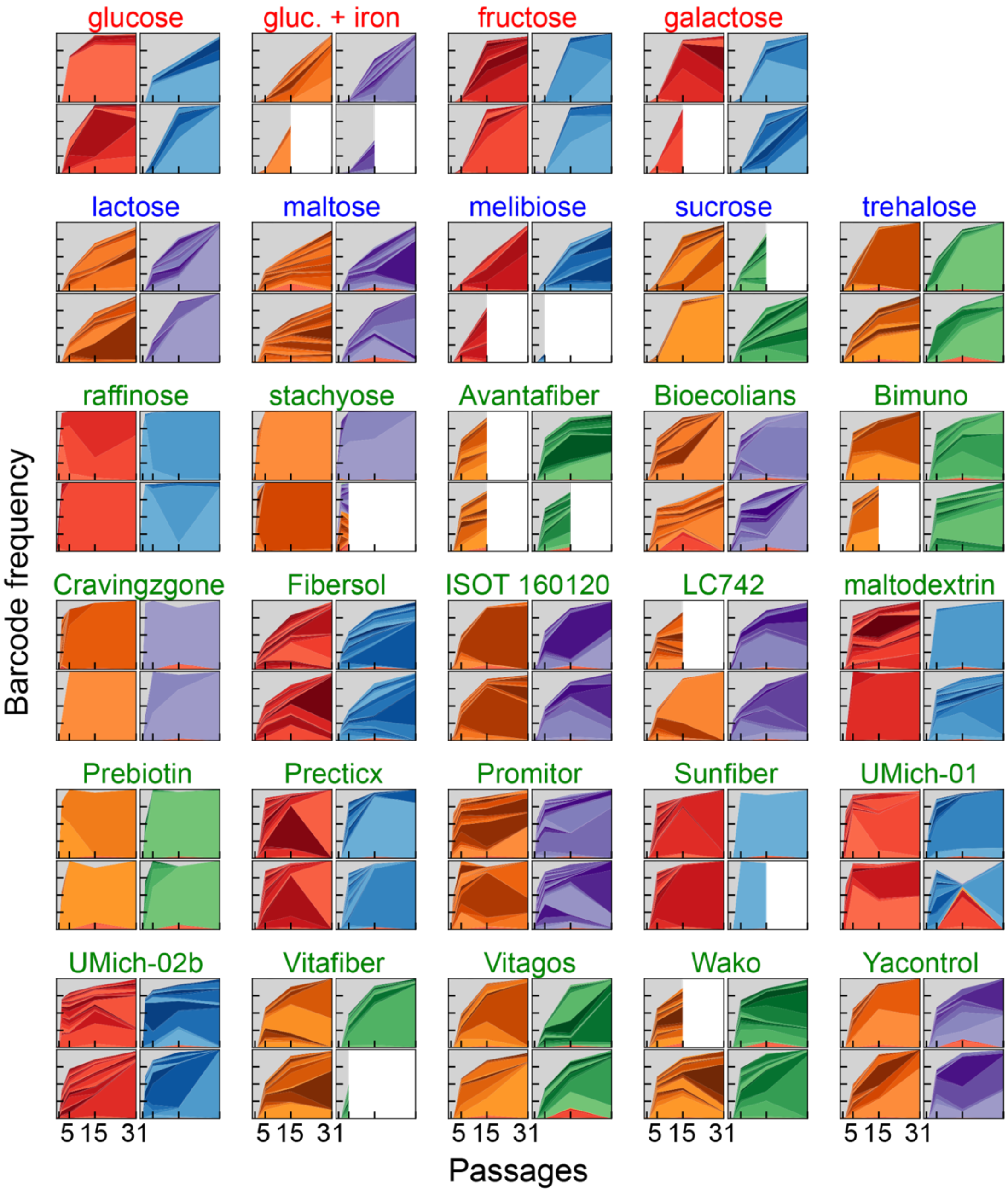


**Figure S17: Barcode lineage dynamics across all carbon sources.** As in Fig. 6C, high-frequency lineages (>1% at some time point in a well) are shaded according to their inferred inoculum (Supplementary Text). Gray bands represent the sum of lineages failing to reach 1%. Red, blue, and green carbon sources are mono-, di-, and poly-saccharides, respectively.

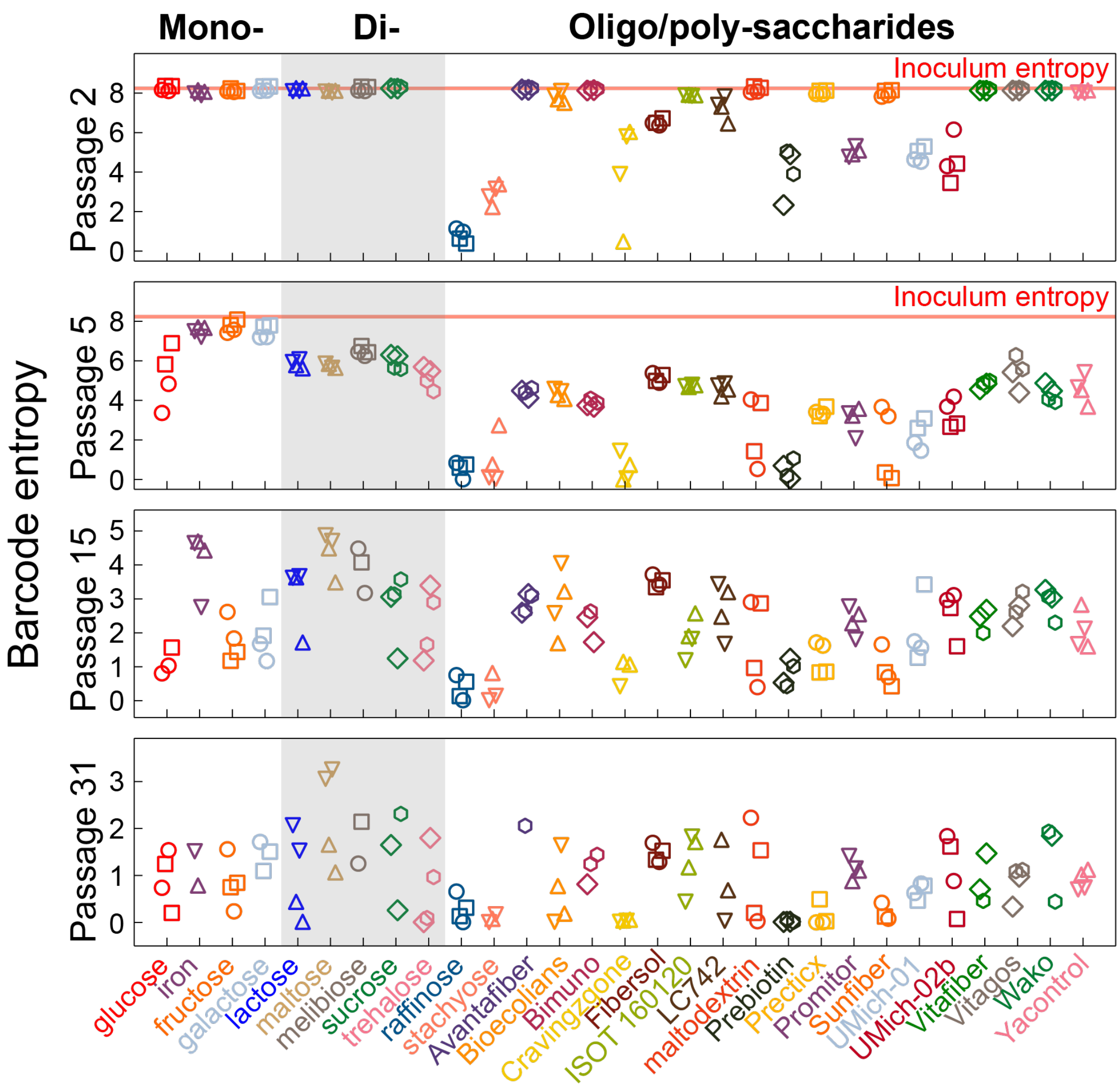


**Figure S18: Shannon entropy (diversity) over time across all carbon source evolution conditions.** Four barcode populations (wells) are shown for each carbon source, and marker shapes denote barcode inoculum. The red line in the top two panels represents the entropy of the input libraries (passage 0). At passage 5, the consistently lower entropy in polysaccharides compared with mono- and di-saccharides is not trivially explained by different population bottlenecks due to variable growth per dilution cycle (Fig. S17).

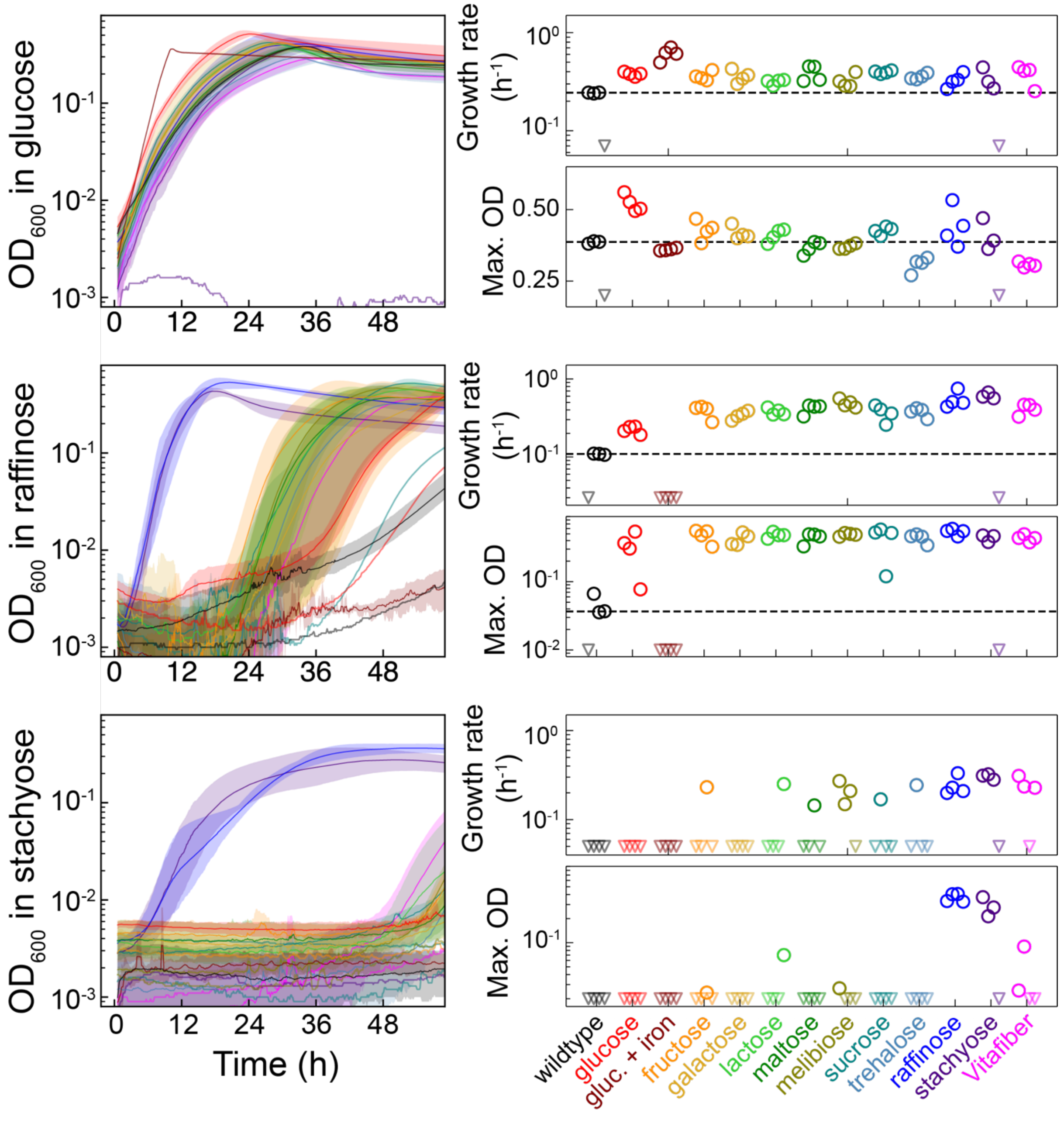


**Figure S19: Growth curves in single carbon sources of *Bt* populations passaged in various carbon sources reveal evolved pleiotropies.** *Bt* populations passaged in 12 single carbon sources for 31 48-hour passages, along with wild-type *Bt*, were then grown in glucose, raffinose, or stachyose. Colors denote passaging condition (i.e., original carbon source). Left: mean (solid line) and range (semi-transparent region) of optical density (OD) across four replicate populations per evolution condition, with the background OD from blank wells subtracted. Right: growth rate and maximum (Max.) OD for each carbon source. Growth rate was calculated as the maximum log(fold-change per hour) over a 6-h window. By both measures of growth, evolved populations can perform better or worse than wild type in a new carbon source (most notably in glucose). Each marker is a single population; triangles denote populations with negligible growth. Dashed black lines represent the median of the four wild-type replicates.

### Supplementary Table Legends

**Table S1: Putative simple point mutations, short indels, and complex structural variants driving adaptation in at least one barcode.**

**Table S2: Junction evidence for structural variants driving adaptation.**

**Table S3: 16S rRNA gene sequences and abundances for the synthetic community of 49 strains isolated from a single human donor used to inoculate germ-free mice.**

**Table S4: NMR profiles of metabolites detected in germ-free mouse ceca and feces.**

**Table S5: Growth curves for *Bt* populations during *in vitro* evolution in various carbon sources.**

**Table S6: Mutations that expanded to high frequency in parallel with the growth of specific barcodes *in vitro*.**
